# OmniTCR: a foundation model unifying T cell receptor recognition prediction and conditional sequence generation

**DOI:** 10.64898/2026.09.10.750588

**Authors:** Feiran Zeng, Duanyu Feng, Dandan Song, Li Ding, Zhenlin Tan, Qian Lei, Wenqiang Lei, An-Yuan Guo

## Abstract

T cell receptor (TCR) recognition prediction and receptor generation are traditionally modelled separately, leaving vast TCR sequence collections disconnected from smaller TCR–peptide–MHC datasets. Here we present OmniTCR, a 113-million-parameter autoregressive foundation model pretrained on 328 million formatted human immune-sequence records. Sequence-type tokens and complementary component orders enable joint learning from individual TCR chains and partial or complete TCR–pMHC associations. On unseen epitopes, OmniTCR achieved AUPRCs of 0.7009 for peptide– TCRβ recognition and 0.8235 for TCR–pMHC interaction prediction, exceeding the strongest evaluated comparators by 0.3396 and 0.3451, respectively. It distinguishes cancer from healthy repertoires across 11 independent pan-cancer cohorts (mean AUROC, 0.9436). The model achieved the highest sequence recovery on internal and external generation benchmarks. Structural modelling supported the plausibility of selected pMHC-conditioned CDR3β candidates. OmniTCR bridges heterogeneous immune sequence data, providing a foundation for computational immunology and receptor design.

## Introduction

T cells recognize peptide antigens presented by major histocompatibility complex (MHC) molecules through their T cell receptors (TCRs)^1–4^. Productive TCR-pMHC (peptide-MHC) recognition can expand specific T cell clones and reshape repertoire composition, creating sequence patterns associated with infection, cancer and other immune states^5, 6^. The same TCR-pMHC relationships guide the identification and engineering of antigen-reactive receptors^7–9^. These relationships are therefore relevant to both receptor-based therapeutics and blood-based immune monitoring^10, 11^. A unified computational model should learn these relationships from available sequence data and make them useful for both distinguishing disease-associated TCR repertoires and generating TCRβ sequences for specified pMHCs.

However, learning these relationships in a unified model is challenging, because TCR recognition is highly diverse and sparsely observed experimentally^12–14^. Somatic recombination generates vast diversity in TCRα- and TCRβ-chain sequences^15–18^, while MHC polymorphism further increases the range of pMHC contexts^19, 20^. At the same time, distinct receptors can engage the same pMHC^21, 22^, and an individual receptor can respond to several antigenic contexts^23^. But experimental platforms can only capture sparse subsets of these relationships and provide different levels of information^24, 25^. Repertoire sequencing provides large collections of TCR sequences^26^, but usually lacks cognate pMHC assignments. Paired-chain sequencing links TCRα and TCRβ sequences, whereas antigen-specific assays connect peptides or pMHCs to one or both chains, but at much smaller scale. Available datasets therefore contain differently sized and often incomplete combinations of peptide, MHC, TCRα (TRA) and TCRβ (TRB) records^27–30^. These incomplete records provide complementary information about recognition but are difficult to integrate within a common modelling framework.

Computational methods have largely reflected these data boundaries by focusing on isolated sub-tasks. In recognition prediction, although predicting complete TCR-pMHC interactions is the central goal, established methods typically address only pairwise relationships. NetMHCpan^31^ predicts peptide-MHC presentation, whereas TEIM^32^ models peptide-TCR interactions. More recent methods such as pMTnet^33^, UniPMT^34^ and UnifyImmun^35^ have begun to model tripartite interactions within a single framework. At the repertoire level, DeepCAT^36^ and iCanTCR^37^ use collections of TCR sequences to distinguish cancer from control repertoires, without requiring cognate pMHC information^38^. For receptor generation, TCRdesign^39^ and TCRT5^40^ rely on antigen-associated TCR sequences to generate TCR sequences for specified targets. These approaches address different parts of the recognition process, but integrating abundant repertoire sequences with sparse multicomponent annotations across prediction and generation remains challenging.

To address this separation, we developed OmniTCR, a 113-million-parameter autoregressive foundation model pretrained on 328 million human immune-sequence records. OmniTCR represents peptide, MHC, TRA and TRB as distinct sequence components, allowing individual sequences and multicomponent records to be learned within a single pretraining framework. During pretraining, OmniTCR uses complementary component orders for multicomponent records, enabling it to learn different conditional relationships among the constituent sequences. This design provides a common basis for both recognition prediction and pMHC-conditioned TCRβ generation. Across recognition prediction, cancer-associated repertoire classification and conditional TCR generation, OmniTCR showed consistent performance advantages over previous single task models across internal evaluations, external datasets and previously unseen epitopes. Across diverse benchmarks, OmniTCR consistently outperformed previous specialized models across recognition prediction, pan-cancer repertoire classification, and conditional TCR generation. In particular, OmniTCR demonstrated generalizability to previously unseen epitopes, reliably distinguished cancer-associated repertoires across independent clinical cohorts, and generated pMHC-conditioned TCRβ sequences with high structural confidence from AlphaFold 3. These results show that a single pretrained foundation model can bridge the gap between immune recognition and receptor generation, offering a general platform for decoding adaptive immunity and accelerating antigen-specific receptor design.

## Results

### OmniTCR jointly learns from immune-sequence records with different component combinations

We assembled 328,232,215 formatted pretraining sequences from human immune-sequence records containing different combinations of peptide, MHC, TRA and TRB (**Fig. 1a** and **Supplementary Tables S1** and **S2**). The corpus combined large TCR sequence collections with smaller multicomponent association datasets, whose coverage varied substantially across component combinations and antigenic contexts (**Supplementary Fig. 1**). Each record was represented using the same component-aware amino-acid sequence format, with [EPI], [HLA], [TRA] and [TRB] sequence-type tokens preserving component identity and boundaries. A 113-million-parameter, 12-layer decoder-only Transformer was pretrained across all record types by causal next-token prediction (**Fig. 1b**). Selected multicomponent records were also presented in complementary component orders, exposing the model to different conditioning relationships while preserving the native sequence within each component.

**Fig. 1.**
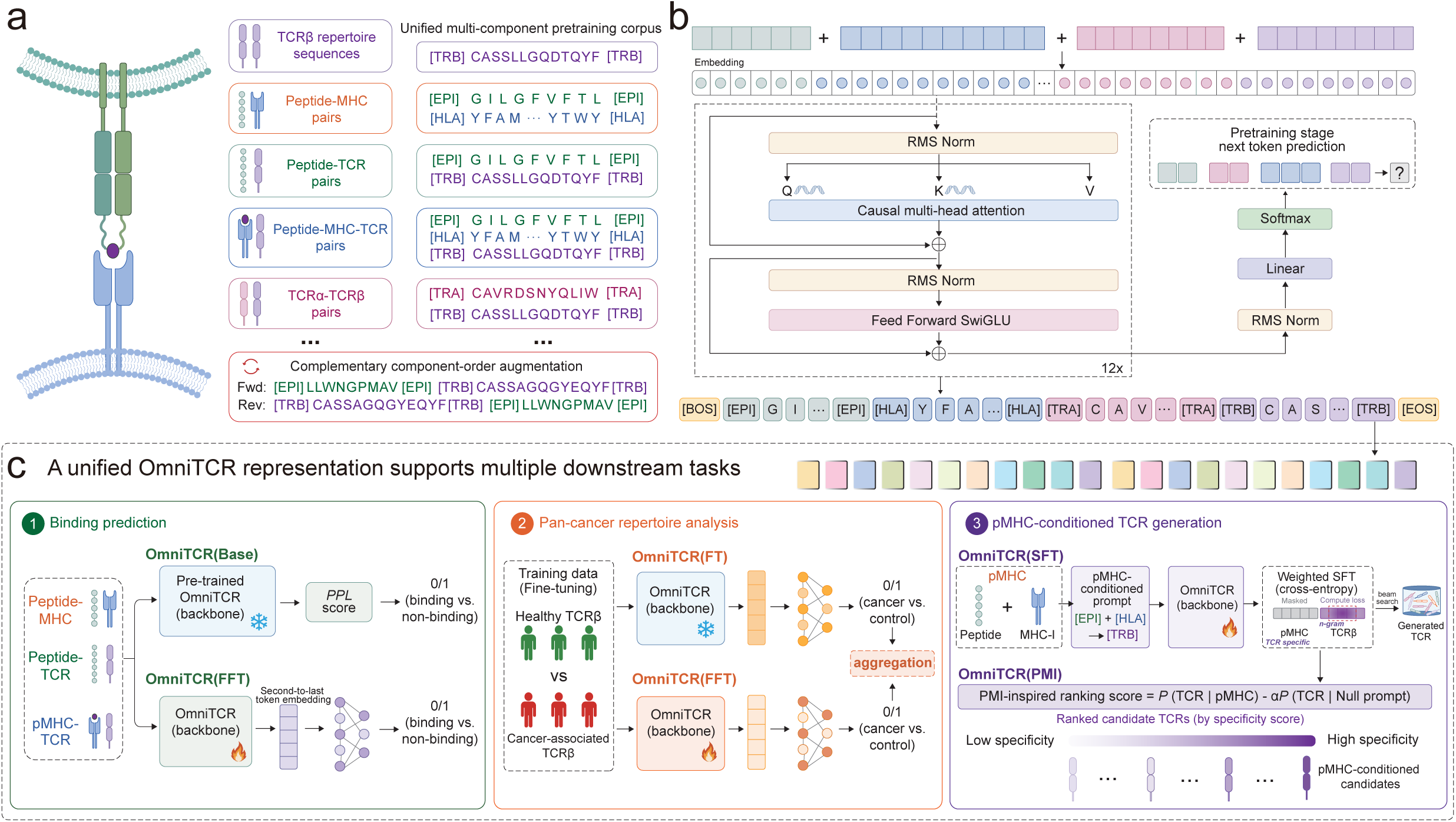
OmniTCR learns a shared autoregressive representation of TCR recognition. **a**, Construction of the 328,232,215 formatted pretraining sequences from large TCR sequence collections and available peptide-MHC, peptide-TCR, peptide-MHC-TCR and paired TCRα-TCRβ association records. Records were formatted using sequence-type tokens to preserve component identity and boundaries, and selected multicomponent records were presented in complementary component orders without reversing the native sequence within any component. **b**, Decoder-only OmniTCR architecture and causal next-token pretraining with component-aware boundary tokens. **c**, Adaptation of the pretrained backbone for recognition prediction, cancer-associated repertoire classification and pMHC-conditioned TCRβ generation. OmniTCR(Base) denotes pretrained likelihood scoring without task-specific adaptation; OmniTCR(FT) denotes frozen-backbone fine-tuning; OmniTCR(FFT) denotes full-parameter fine-tuning; OmniTCR(SFT) denotes supervised fine-tuning for pMHC-conditioned TRB generation; and OmniTCR(PMI), pointwise mutual-information-inspired reranking of the fixed TRB candidate pool generated by OmniTCR(SFT).

The same pretrained backbone was subsequently evaluated and adapted across three tasks (**Fig. 1c**): recognition prediction, including peptide-MHC presentation, peptide-TCRβ recognition and TCR-pMHC interaction prediction; cancer-associated repertoire classification; and pMHC-conditioned TRB sequence generation. These analyses tested whether a shared pretrained representation could transfer across different component combinations, biological scales and prediction and generation tasks.

### Pretraining captures recognition-related information before task-specific adaptation

We first evaluated whether shared pretraining alone had already captured the relationships underlying TCR recognition before any task-specific adaptation. We used held-out peptide-MHC (P-M), peptide-TCRβ (P-T) and peptide-MHC-TCRβ (P-M-T) test sets containing 85,506, 91,632 and 13,662 records, respectively. These records were excluded from both autoregressive pretraining and downstream fine-tuning. Positive records represented experimentally supported molecular associations. P-M negatives were generated by recombining peptides and MHC after excluding known associations, whereas P-T and P-M-T negatives were experimentally annotated non-binding records.

Without task-specific adaptation, OmniTCR(Base) assigned lower perplexity to experimentally supported records in all three settings (all adjusted *P* < 2.2 × 10^−16^; **Fig. 2a**). Rank-biserial effect sizes were 0.9022 for P-M, 0.7062 for P-T and 0.5061 for P-M-T, indicating that even the pretraining foundation model alone can intrinsically separate potential interactions from non-binding pairs. We also evaluated whether peptide-associated information was detectable from TRB representations alone. We extracted closing [TRB] token embeddings from 28,820 unique CDR3β sequences and grouped them into 50 clusters without using peptide identities or association labels. We then mapped 34,261 supported peptide-TRB associations spanning ten peptide epitopes onto the 50 clusters. Peptide identity was associated with cluster membership (bias-corrected Cramér’s V = 0.2251; permutation *P* = 9.99 × 10^−5^; **Fig. 2b**, **Supplementary Fig. 2a**). Cluster profiles were more similar among peptides from the same SARS coronavirus group than between SARS-associated and reference viral peptides (mean Jensen-Shannon similarity, 0.8876 versus 0.4862; *P* = 5.0 × 10^−4^ ; **Supplementary Fig. 2b, c**). Profile similarity also increased with peptide-sequence similarity (Spearman’s *ρ* = 0.4233, *P* = 0.0038 ; **Supplementary Fig. 2d**). Recognition-related information was therefore learned during pretraining and was detectable in both autoregressive likelihoods and TRB representations before task-specific adaptation.

**Fig. 2.**
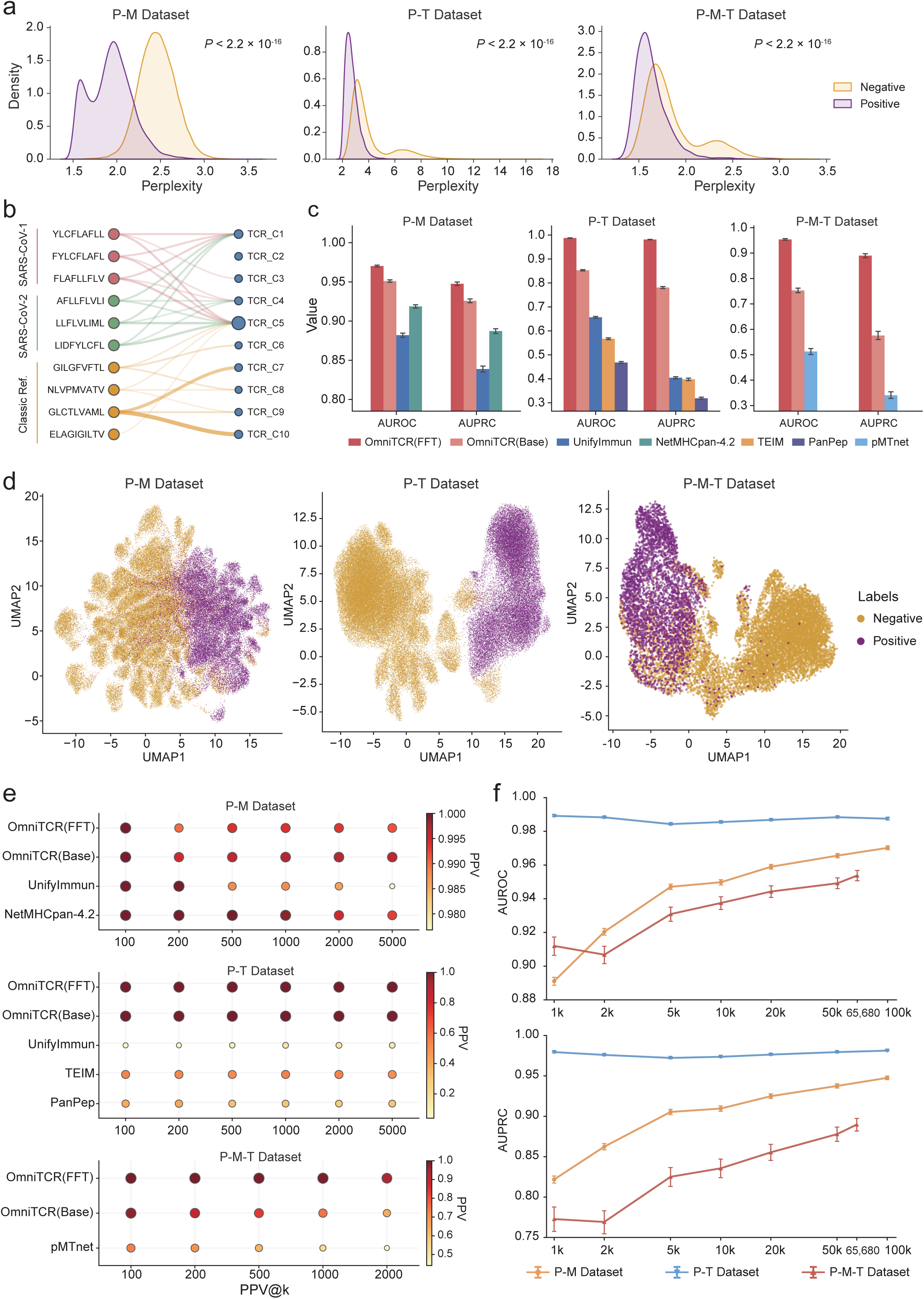
Recognition-related information learned by OmniTCR. **a**, Perplexity distributions for experimentally supported and negative P-M (n = 85,506), P-T (n = 91,632) and P-M-T (n = 13,662) records under OmniTCR(Base). P values were calculated using two-sided Mann-Whitney U tests and adjusted for multiple comparisons using the Benjamini-Hochberg procedure. **b**, Mapping of ten peptides onto ten TRB clusters selected from 50 by peptide-specific enrichment (Supplementary Section S3.2). Up to three highest-count connections per peptide are shown. Edge width reflects association-record count; cluster-node area reflects the number of unique CDR3β sequences. **c**, AUROC and AUPRC for the three internal recognition prediction tasks. **d**, UMAP projections of the corresponding representations from OmniTCR(FFT) of experimentally supported and negative records. **e**, Positive predictive value among the top-k predictions (PPV@k). **f**, AUROC and AUPRC after fine-tuning with 1,000-100,000 (P-M-T up to 65,680) experimentally supported records and an equal number of negative records. Error bars in **c** and **f** show 95% confidence intervals estimated by bootstrap with 10,000 resamples. P-M, peptide-MHC presentation; P-T, peptide-TCRβ recognition; P-M-T, peptide-MHC-TCRβ interaction prediction.

### OmniTCR enables accurate and data-efficient recognition prediction

We then tested whether the recognition-related information learned during pretraining could be translated into supervised prediction. Separate OmniTCR(FFT) models were trained for P-M, P-T and P-M-T prediction using the same pretrained backbone (**Fig. 2c**). Without task-specific fine-tuning, OmniTCR(Base) already achieved AUPRC values of 0.9259, 0.7810 and 0.5757 across the three binding prediction tasks. After full-parameter fine-tuning, OmniTCR(FFT) achieved AUROC/AUPRC values of 0.9702/0.9476 for P-M, 0.9875/0.9813 for P-T and 0.9538/0.8898 for P-M-T. For each task, we compared our method against the best-performing specialized baseline on the corresponding test set. The largest gains over specialized methods occurred in TCR-dependent tasks. AUPRC increased by 0.5767 over UnifyImmun^35^ for P-T and by 0.5492 over pMTnet^33^ for P-M-T (both *P* < 1.0 × 10^−4^). UMAP projections of OmniTCR(FFT) embeddings also showed separation between supported and negative records in the TCR-dependent settings (**Fig. 2d**).

The advantage also extended to retrieval among the highest-ranked predictions. We measured positive predictive value among the top-k predictions (PPV@k; **Fig. 2e**). OmniTCR(FFT) achieved a PPV@5,000 of 1.0000 for P-T, compared with 0.4920 for TEIM. For P-M-T, its PPV@2,000 was 0.9395, compared with 0.6430 for OmniTCR(Base) and 0.4415 for pMTnet.

We also tested how much task-specific supervision was required to obtain these gains. Models were fine-tuned with 1,000-100,000 (P-M and P-T used up to 100,000 supported records, whereas P-M-T included the full set of 65,680 supported records) supported records and an equal number of negatives. Every model was evaluated on the same fixed internal test set (**Fig. 2f**). With only 1,000 positive examples, OmniTCR achieved AUROC/AUPRC values of 0.8911/0.8218 for P-M, 0.9892/0.9795 for P-T and 0.9120/0.7728 for P-M-T. P-T performance was already close to its observed plateau. P-M and P-M-T continued to improve as more examples were introduced. These results show that the pretrained representation supported strong recognition prediction with limited task-specific supervision, including for the more sparsely observed P-M-T setting.

### OmniTCR generalizes across external datasets, paired TCRs and unseen epitopes

We next evaluated generalization to independently assembled recognition datasets. External evaluation included the NetMHCpan-4.1 P-M benchmark^31^ (6,042 records), TRAIT P-T and P-M-T sets^30^ (6,842 and 5,945 records), and independently assembled SARS-CoV-2 P-T and P-M-T sets^41^ (6,470 and 6,488 records; **Fig. 3a, b**). Exact overlaps with training data from OmniTCR were removed. Dataset compositions and negative definitions are reported in **Supplementary Table S3 and Section S1.2**.

**Fig. 3.**
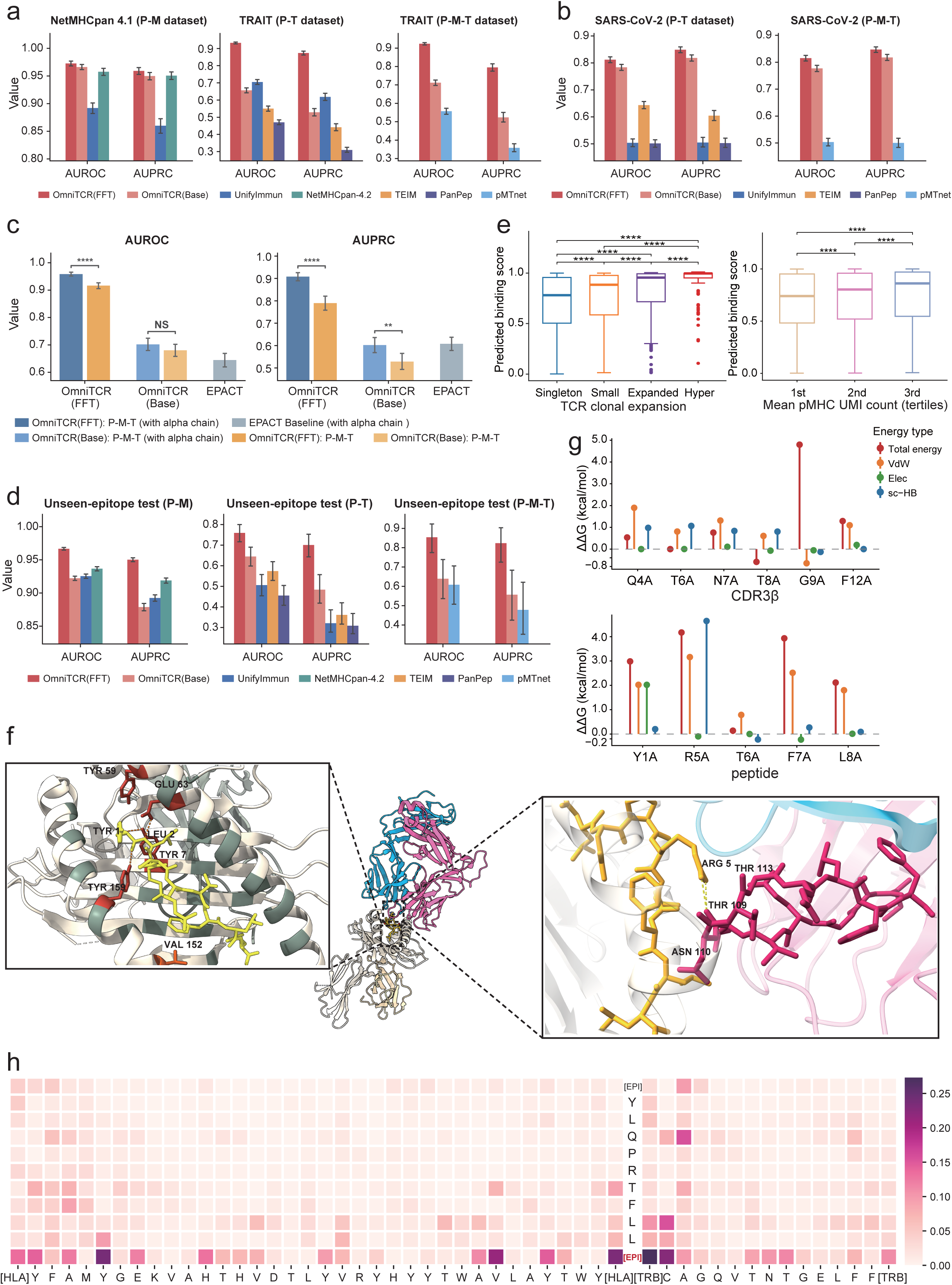
Generalization of OmniTCR and correspondence with cellular and structural evidence. **a**, Recognition prediction performance on NetMHCpan-4.1 P-M dataset (n = 6,042) and TRAIT P-T (n = 6,842) and P-M-T (n = 5,945) datasets. **b**, Recognition prediction performance on SARS-CoV-2 P-T (n = 6,470) and P-M-T (n = 6,488) datasets. **c**, P-M-T prediction with and without the paired TCRα chain in 2,535 matched TRAIT records. P values were calculated using paired bootstrap tests with 10,000 resamples. **d**, Recognition prediction performance on unseen-epitope P-M (n = 34,750), P-T (n = 642) and P-M-T (n = 141) test sets. **e**, OmniTCR(FFT) scores stratified by clonotype expansion and pMHC-barcode UMI tertile across 16,767 observations from four donors. Boxes show medians and interquartile ranges. P values were calculated using one-sided Mann-Whitney U tests and adjusted using the Benjamini-Hochberg procedure. **f**, Experimentally resolved peptide-HLA and peptide-CDR3β contacts in the TCR-pMHC complex (PDB ID: 7N6E). **g**, FoldX alanine scanning of selected CDR3β and peptide residues in the TCR-pMHC complex (PDB ID: 7N6E). **h**, Final-layer OmniTCR(Base) attention across the corresponding peptide, HLA and CDR3β sequence positions. Error bars in **a-d** show 95% confidence intervals. NS, P ≥ 0.05; **, P < 0.01; ***, P < 0.001; ****, P < 0.0001.

OmniTCR generalized across all external settings (**Fig. 3a, b**). For each task, we compared our method against the best-performing specialized baseline on the corresponding test set. On the NetMHCpan-4.1 P-M benchmark, OmniTCR(FFT) achieved an AUROC/AUPRC of 0.9727/0.9592, exceeding NetMHCpan-4.2^42^ by 0.0151/0.0086 (*P* = 3.33 × 10^−7^ and *P* = 0.0033). Larger gains were observed for receptor-dependent recognition. On TRAIT datasets, OmniTCR(FFT) achieved an AUROC/AUPRC of 0.9325/0.8736 for P-T and 0.9235/0.7949 for P-M-T. These values exceeded UnifyImmun^35^ by 0.2274/0.2552 and pMTnet by 0.3666/0.4369, respectively (all AUROC *P* < 2.2 × 10^−16^ and AUPRC *P* < 1.0 × 10^−4^ for each task) (**Fig. 3a**). On the SARS-CoV-2 datasets, OmniTCR(FFT) achieved an AUROC/AUPRC of 0.8119/0.8489 for P-T and 0.8152/0.8471 for P-M-T. The corresponding AUPRC gains were 0.2444 over TEIM and 0.3470 over pMTnet (both *P* < 1.0 × 10^−4^) (**Fig. 3b**). OmniTCR(Base) also retained AUPRC values of 0.8185 and 0.8179 on these two datasets without task-specific adaptation. UMAP projections showed corresponding separation of experimentally supported and negative records, with task-dependent organization (**Supplementary Fig. 2e, f**).

The component-aware format also enabled the same model to use paired TCRα information without architectural changes. We evaluated 2,535 matched TRAIT peptide-MHC-TRA-TRB records with or without the TRA component (**Fig. 3c**). Adding TRA increased OmniTCR(FFT) AUROC from 0.9163 to 0.9584 and AUPRC from 0.7898 to 0.9083 (both *P* < 1.0 × 10^−4^). In the paired-chain setting, OmniTCR(FFT) exceeded EPACT^43^ by 0.3138 in AUROC and 0.3004 in AUPRC (*P* < 2.2 × 10^−16^ and *P* < 1.0 × 10^−4^, respectively). Paired TRA information also increased OmniTCR(Base) AUPRC by 0.0741 (*P* = 0.0015), indicating that paired-chain information was already accessible to the pretrained model.

We evaluated generalization to unseen epitopes using epitope-disjoint test sets (**Supplementary Table S4**). To prevent data leakage, all records containing these evaluation peptide epitopes were completely excluded from both autoregressive pretraining and task-specific fine-tuning. The P-M, P-T and P-M-T test sets contained 34,750, 642 and 141 records, respectively (**Fig. 3d** and **Supplementary Table S4**). OmniTCR(FFT) achieved AUROC/AUPRC values of 0.9667/0.9502 for P-M, 0.7597/0.7009 for P-T and 0.8542/0.8235 for P-M-T. These values exceeded the strongest task-matched comparators by 0.0301/0.0314 (*P* < 2.2 × 10^−16^ and *P* < 1.0 × 10^−4^), 0.1855/0.3396 (*P* = 1.97 × 10^−11^ and *P* < 1.0 × 10^−4^) and 0.2457/0.3451 (*P* = 6.3 × 10^−13^ and *P* < 1.0 × 10^−4^), respectively. UMAP projections also showed corresponding separation of experimentally supported and negative records (**Supplementary Fig. 2g**).

### OmniTCR scores capture cellular and structural features of TCR recognition

We evaluated whether OmniTCR scores were associated with cellular measurements related to antigen recognition. We analyzed antigen-specific CD8⁺ T cells from four healthy donors in the 10x Genomics dataset^44^. Cells were retained when they contained one productive TCRβ sequence and one pMHC-multimer assignment. After quality control, 16,767 pMHC-TCRβ observations were grouped by clonotype and specificity. OmniTCR scores increased with clonotype expansion (**Fig. 3e**). Scores were higher in small than singleton clonotypes (adjusted *P* = 2.2 × 10^−9^) and in hyperexpanded than expanded clonotypes (adjusted *P* = 1.6 × 10^−8^ ; Mann-Whitney U tests with Benjamini-Hochberg correction). Scores also increased across tertiles of mean pMHC-barcode UMI abundance (adjusted *P* = 3.7 × 10^−12^ and 1.1 × 10^−12^ for successive tertiles).

We compared model attention with an experimentally resolved TCR-pMHC interface using the NR1C TCR bound to YLQPRTFLL-HLA-A*02:01 (PDB ID: 7N6E)^45^. The peptide contacted both the HLA binding groove and the CDR3β loop in the experimental structure (**Fig. 3f**). At the peptide-HLA interface (**Fig. 3f**, left inset), the HLA residues contacting peptide Tyr1 and Leu2 (such as Tyr59, Glu63, and Val152) directly coincide with polymorphic HLA pseudo-sequence positions (shown in grey-green ribbon, with red highlights indicating residues receiving high model attention). At the TCR-binding interface (**Fig. 3f**, right inset), the central peptide residue Arg5 (R5) forms direct, close contacts with the CDR3β Thr109-Asn110-Thr113 region (corresponding to the ‘T-N-T’ segment in **Fig. 3h**). To evaluate the energetic significance of these contact sites, we performed FoldX^46^ alanine scanning (**Fig. 3g**), confirming peptide R5 and its interacting partners as critical biophysical binding hotspots. Interestingly, without any structural supervision during pretraining, final-layer self-attention from OmniTCR(Base) concentrated on these implicated HLA contact residues and the central CDR3β TNT segment contacting peptide R5 (**Fig. 3h**), where [EPI] denotes the peptide epitope boundary. Layer-wise integrated gradients showed redistribution across residues and component boundaries over successive Transformer layers (**Supplementary Fig. 3a, b**). These results showed that OmniTCR attention patterns potentially capture structural features of TCR recognition.

### OmniTCR identifies cancer-associated repertoires across cohorts

To assess transfer to cancer-associated repertoire classification, we assembled receptor-level training sets from repertoire data obtained from TCRdb 2.0. We constructed a healthy reference library from the 1,504 healthy training repertoires. GLIPH2^47^ clustering of TRBs from 6,078 cancer repertoires, followed by exclusion of healthy-reference sequences, yielded 1,489,111 candidate cancer-associated CDR3β sequences. These candidates were paired with an equally sized healthy-background set matched by clonotype frequency (**Fig. 4a** and **Supplementary Section S1.3**). The cancer and healthy sets had similar global CDR3β motifs but also showed modest differences in length, amino-acid usage and physicochemical properties (**Fig. 4b** and **Supplementary Fig. 4a-c**).

**Fig. 4.**
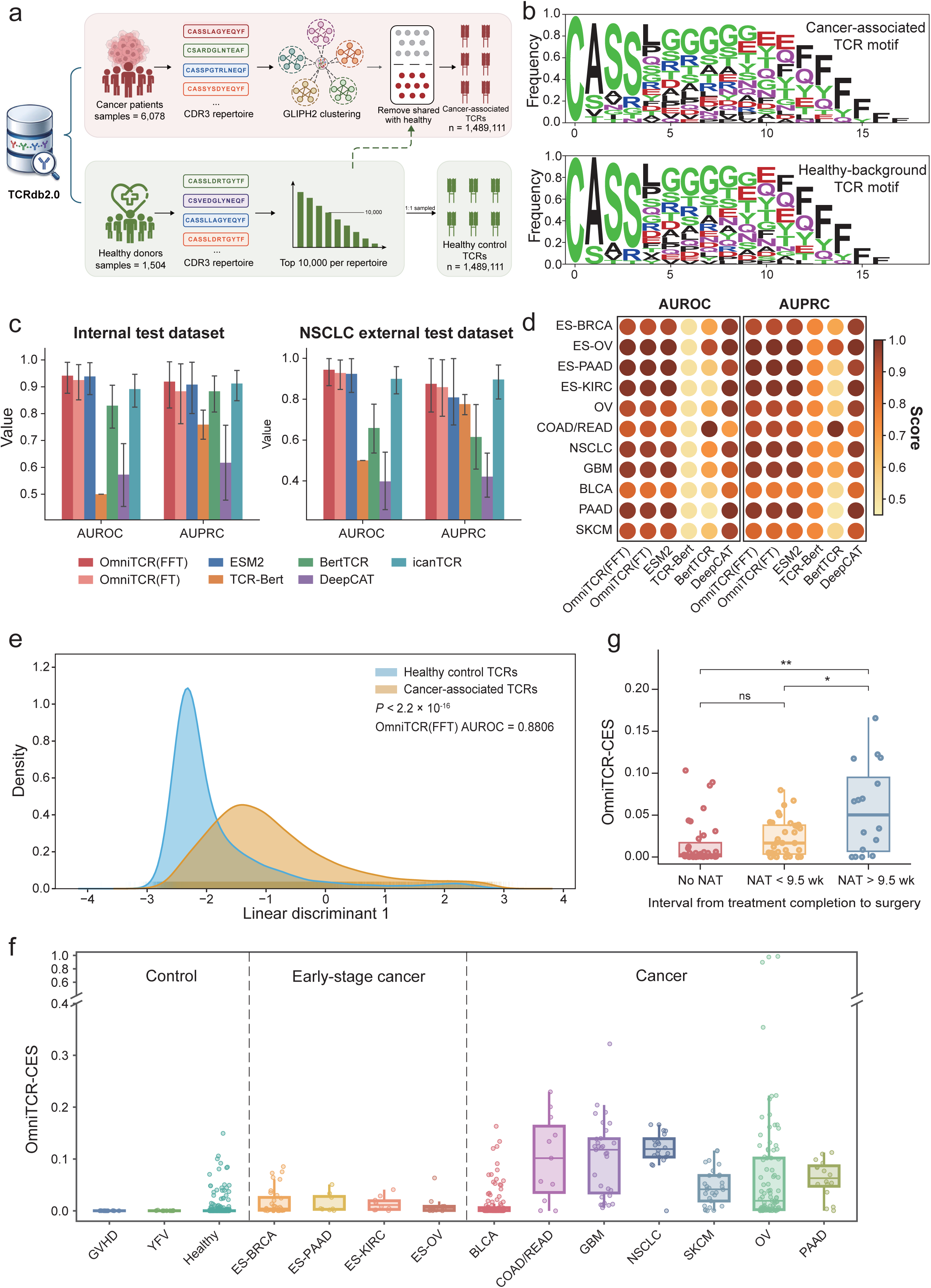
Transfer of receptor-level representations to cancer-associated repertoire classification. **a**, Construction of 1,489,111 candidate cancer-associated CDR3β sequences and an equally sized, clonotype frequency-matched healthy-background set from 6,078 cancer and 1,504 healthy repertoires. **b**, Position-specific amino-acid composition of the candidate cancer-associated and healthy-background CDR3β receptor sets. **c**, Repertoire-level classification performance on the internal set (53 cancer and 43 healthy repertoires) and the external NSCLC set (43 tumour-region repertoires from 15 patients and 40 healthy repertoires). **d**, AUROC and AUPRC across 11 independent cohorts spanning nine cancer types. **e**, Projection of 29,829 VDJdb tumour-antigen-associated sequences and 29,829 healthy-background sequences onto a fixed linear discriminant axis fitted using OmniTCR(FFT) embeddings from the training data. P values were calculated using a two-sided Welch’s t-test. AUROC was calculated separately from direct OmniTCR(FFT) predictions. **f**, OmniTCR-CES across control (n = 690), early-stage cancer (n = 60) and established cancer (n = 323) repertoires. **g**, OmniTCR-CES in OSLO-COMET repertoires grouped as untreated (n = 40), less than 9.5 weeks (n = 36) or more than 9.5 weeks (n = 16) from neoadjuvant therapy to surgery. P values were calculated using one-sided Mann-Whitney U tests. Error bars in **c** show 95% confidence intervals estimated by bootstrap with 10,000 resamples. Boxes in **f** and **g** show medians and interquartile ranges. NS, P ≥ 0.05; *, P < 0.05; **, P < 0.01.

Using these receptor sets, we trained a frozen-backbone classifier, OmniTCR(FT), and a full-parameter model, OmniTCR(FFT). We then aggregated receptor-level predictions into a cancer-associated TCR enrichment score (OmniTCR-CES, defined as the median probability among up to the 1,000 most abundant clonotypes). We used OmniTCR-CES to evaluate whether receptor-level representations learned by OmniTCR could support cancer-associated repertoire classification. On our internal held-out test dataset with 53 cancer and 43 healthy repertoires, OmniTCR(FFT) achieved the highest performance with an AUROC/AUPRC of 0.9414/0.9190, compared with 0.9251/0.8832 for OmniTCR(FT) (**Fig. 4c**). In an external multiregional NSCLC dataset^48^ containing 43 tumour-region repertoires from 15 patients, OmniTCR(FFT) achieved 0.9441/0.8750, whereas OmniTCR(FT) achieved 0.9284/0.8588.

We then evaluated different models across 11 independent external cohorts from DeepCAT^36^ spanning nine cancer types (**Supplementary Table S5**). OmniTCR(FFT) achieved mean AUROC and AUPRC values of 0.9436 and 0.9410 (**Fig. 4d**). Both mean metrics were approximately 0.02 higher than those of ESM-2, the strongest comparator by mean performance. Furthermore, across the four early-stage cohorts, OmniTCR(FFT) retained AUROC values of 0.9219-1.0000 and AUPRC values of 0.9266-1.0000. OmniTCR(FT) without full-parameter fine-tuning still achieved mean AUROC and AUPRC values of 0.9404 and 0.9388. The performance of OmniTCR(FT) showed that fixed pretrained representations supported cancer-associated classification, with full-parameter fine-tuning providing further refinement.

### Cancer-associated predictions generalize to tumour-antigen-associated TCRs and treatment-associated cohorts

We further tested whether the learned cancer-associated receptor information from repertoire-derived labels was also detectable in TCRs independently annotated by tumour-antigen recognition. We curated 29,829 unique TCRβ sequences from VDJdb^27^ with reported recognition of cancer-testis antigens, differentiation or overexpressed tumour antigens. An equally sized healthy-control set was assembled after standardization, deduplication and removal of training overlaps. A linear discriminant model was fitted to fixed OmniTCR(FFT) embeddings from the training candidate cancer-associated and healthy-background receptors. The fixed discriminant axis was then applied to the VDJdb and healthy-control sequences without refitting. Projected scores differed between tumour-antigen-reactive and healthy-control receptors (*P* < 2.2 × 10^−16^; **Fig. 4e**). Direct OmniTCR(FFT) predictions distinguished the same sets with an AUROC of 0.8806.

At the repertoire level, we applied OmniTCR-CES without cohort-specific recalibration to healthy, yellow fever vaccination (YFV) and graft-versus-host disease (GVHD) controls, four early-stage cancer cohorts and seven established-cancer cohorts. The control, early-stage and established-cancer groups contained 690, 60 and 323 repertoires, respectively. CES was low in control repertoires, increased in the early-stage cohorts and was generally higher in established malignancies (**Fig. 4f**). When repertoires were grouped by cohort category, CES increased from control to early-stage cancer (*P* < 2.2 × 10^−16^) and from early-stage to established cancer (*P* = 0.0029; **Supplementary Fig. 4d**).

Finally, we analyzed treatment-associated repertoire variation in the OSLO-COMET colorectal liver-metastasis cohort^49^. The dataset contained 92 repertoires from 85 patients, including repeated observations from some individuals. Samples were further stratified by the prespecified 9.5-week interval from completion of neoadjuvant chemotherapy (NAT) to surgery. The untreated, short-interval and long-interval groups contained 40, 36 and 16 samples, respectively (**Fig. 4g**). At the sample level, CES was higher in long-interval samples than in untreated samples (*P* = 0.0076) or short-interval samples (*P* = 0.0250). Untreated and short-interval groups did not differ significantly (*P* = 0.0605). CES was therefore associated with an independently defined post-treatment interval, extending the repertoire analysis beyond cancer-state discrimination.

### OmniTCR generates pMHC-conditioned TCRβ sequences and reranks target-dependent candidates

We tested whether the learned conditional relationships could support receptor generation for specified pMHCs. OmniTCR(SFT) was fine-tuned on peptide-MHC-TRB associations using a TCR-specific objective that emphasized central CDR3β residues and local sequence patterns (**Supplementary Section S2.3** and **Supplementary Table S8**). We evaluated the model on 20 pMHCs with 3,222 experimentally observed reference TRB sequences that did not overlap the generation training set. For each pMHC, 100 unique candidates were retained, yielding 2,000 generated TRB sequences (**Fig. 5a-d**). The generated sequences reproduced the dominant 13- and 15-residue length modes, although with a narrower distribution (**Fig. 5a**), and showed higher OLGA^50^ generation probabilities than the experimentally observed sequences (*P* < 2.2 × 10^−16^; **Fig. 5b**). Their amino-acid usage also closely matched the observed sequences (Pearson’s *r* = 0.979, *P* = 6.9 × 10^−14^ ; **Fig. 5c**), with conserved terminal patterns and greater variation across central CDR3β positions (**Fig. 5d**).

**Fig. 5.**
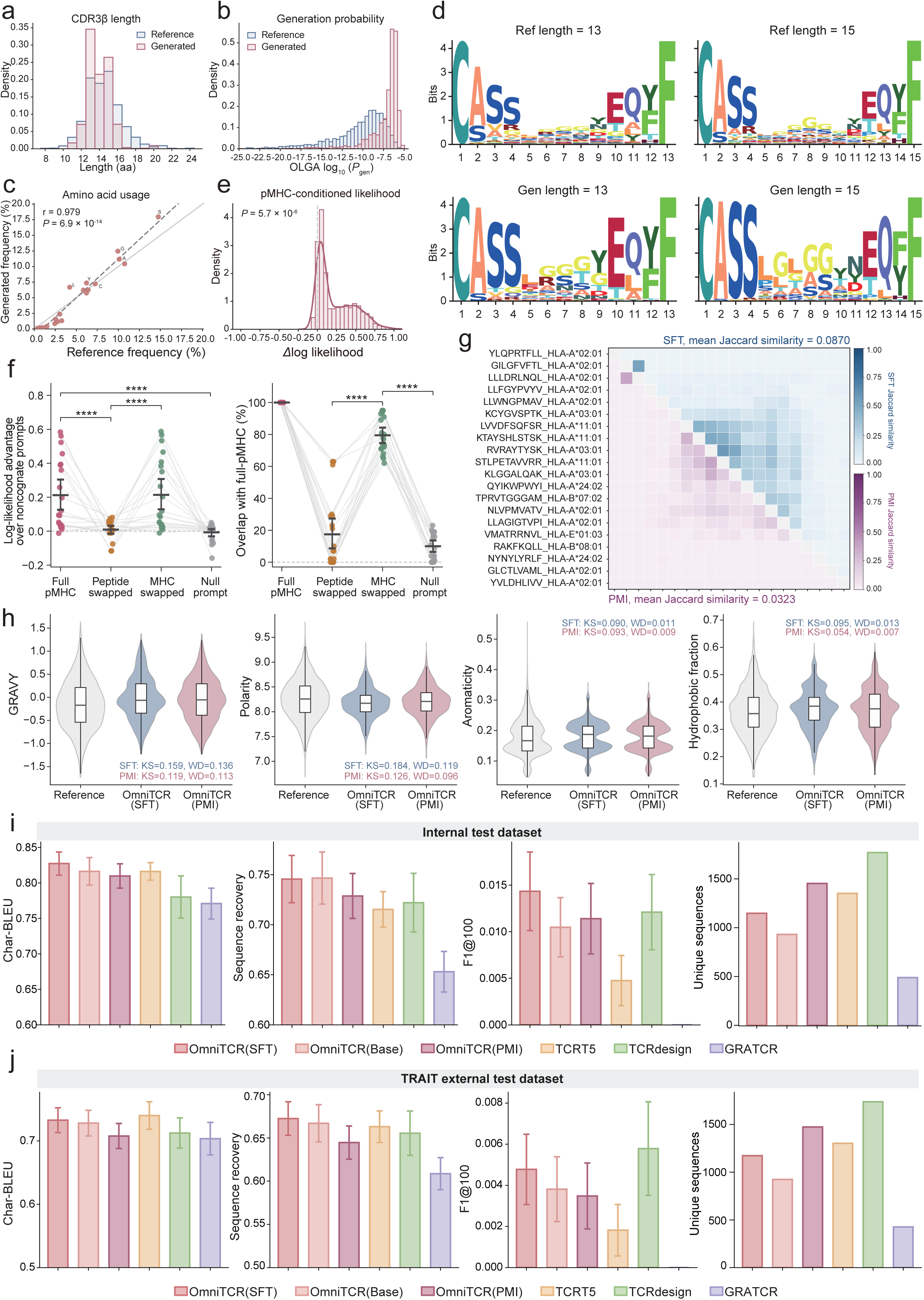
pMHC-conditioned TCRβ generation and PMI-inspired candidate prioritization. **a**, CDR3β length distributions for 3,222 experimentally observed TRB sequences and 2,000 OmniTCR(SFT) generated sequences across 20 pMHCs. **b**, OLGA generation probabilities for experimentally observed and generated CDR3β sequences. **c**, Global amino-acid frequencies for experimentally observed and generated CDR3β sequences. Pearson’s correlation was used to assess agreement in amino-acid frequencies. **d**, Sequence logos for experimentally observed and OmniTCR(SFT) generated CDR3β sequences of lengths 13 and 15. **e**, Candidate-level target-versus-mean-noncognate log-likelihood margins. Statistical inference used a two-sided Wilcoxon signed-rank test on the 20 target-level mean margins. **f**, Effects of pMHC conditioning on log-likelihood margins (left) and generated-set overlap (right). Lines connect matched pMHC targets (n = 20). Black markers and error bars show means and 95% bootstrap confidence intervals. P values were calculated using one-sided paired Wilcoxon signed-rank tests. **g**, Pairwise Jaccard similarity among the top 100 pMHC-conditioned TCRβ generation sets. The upper and lower triangles show OmniTCR(SFT) beam-ranked and OmniTCR(PMI) reranked sequences from the same generation pools, respectively. **h**, GRAVY, polarity, aromaticity and hydrophobic-residue fraction in 3,222 experimentally observed, 2,000 OmniTCR(SFT) generated and 2,000 OmniTCR(PMI) reranked sequences. KS and WD denote Kolmogorov-Smirnov and Wasserstein distances from the experimentally observed sequences, respectively. **i, j**, Char-BLEU, sequence recovery, F1@100 and pooled numbers of unique generated sequences on the internal and TRAIT benchmarks. Bars show mean ± s.e.m. across 20 pMHCs except for unique counts. ****, P < 0.0001.

We next evaluated whether generated candidates received greater model support under their designated pMHC than under alternative prompts. For each generated candidate, we compared its mean log-likelihood under the designated pMHC with its mean log-likelihood under the other 19 pMHC contexts. Candidate-level margins were positive for 90.4% of sequences. Target-level mean margins were greater than zero in a two-sided Wilcoxon signed-rank test across 20 pMHCs (*P* = 5.7 × 10^−6^ ; **Fig. 5e**). To further separate peptide and MHC contributions, we exchanged the peptide, exchanged the MHC or removed both components to form a null prompt. We measured changes in likelihood support and candidate retention relative to the complete pMHC condition (**Fig. 5f**). Peptide exchange reduced both quantities more than MHC exchange (*P_likeli_*_ℎ*ood*_ = 4.1 × 10^−5^; *P_overlap_* = 4.4 × 10^−5^), whereas the null prompt produced the weakest support and retention. Thus, likelihood scoring and regenerated-set comparisons both supported peptide-dependent conditioning within the tested pMHC panel. These patterns were consistent with peptide sequence being a major determinant of TCR antigen specificity within the composite TCR-pMHC interface, while MHC context also contributed to model support.

To prioritize candidates whose likelihood depended more strongly on the supplied pMHC, we applied pointwise mutual information (PMI)-inspired reranking to the fixed OmniTCR(SFT) candidate pools. OmniTCR(PMI) ranked each candidate by the gain in continuation log-likelihood under the target pMHC relative to an empty prompt lacking antigen sequences. This strategy effectively removes non-specific sequence bias, prioritizing candidate receptors with antigen specificity without requiring model retraining. When the top 100 candidates were retained for each pMHC, the mean pairwise Jaccard similarity among the 20 pMHC-specific candidate sets decreased from 0.0870 under the original SFT beam ranking to 0.0323 after PMI reranking, corresponding to a 62.9% reduction in cross-pMHC sequence sharing (**Fig. 5g**). Similar reductions were observed at other candidate-list depths (**Supplementary Fig. 5g**). PMI-ranked candidates retained the main CDR3β length and amino-acid distributions observed after SFT and remained close to experimentally observed TRB sequences in grand average of hydropathicity (GRAVY), polarity, aromaticity and hydrophobic-residue fraction (**Fig. 5h** and **Supplementary Fig. 5a-d**).

We then benchmarked generation on internal and external TRAIT test sets. OmniTCR(SFT), OmniTCR(PMI) and OmniTCR(Base) were compared with three specialized methods: TCRT5^40^, TCRdesign^39^ and GRATCR^51^. Metrics included Char-BLEU, sequence recovery, exact-reference F1@100 and the number of unique sequences. On the internal set, OmniTCR(SFT) exceeded the strongest specialized method by 0.0111 in Char-BLEU and 0.0236 in sequence recovery and achieved the highest F1@100 (**Fig. 5i**). On the TRAIT^30^ dataset, OmniTCR(SFT) achieved the highest sequence recovery, exceeding TCRT5 by 0.0094, while its Char-BLEU was within 0.0074 of the best method (**Fig. 5j**). On the unseen-pMHC benchmark, OmniTCR(SFT) exceeded TCRT5 by 0.0148 in Char-BLEU and 0.0145 in sequence recovery (**Supplementary Fig. 5i**). The TCR-specific objective improved Char-BLEU and sequence recovery over standard token-level SFT across all three generation benchmarks (**Supplementary Section S6.4 and Table S9**). Overall, OmniTCR(SFT) achieved strong pMHC-conditioned TRB generation across internal and external benchmarks, while PMI reranking reduced cross-pMHC sequence sharing and increased target-relative model support.

### Generated TRB candidates form plausible predicted interfaces with unseen pMHCs

We evaluated generated TRB candidates in complete predicted TCR-pMHC assemblies. We selected seven experimentally determined TCR-pMHC complexes from TCR3d database^52^ that were first released in the PDB after 8 April 2025. Each contained a target peptide absent from OmniTCR training data. For each pMHC, 100 candidates per method were inserted into the matched TCRβ framework and modeled with the experimentally reported TCRα chain, peptide, MHC class I heavy chain and β2-microglobulin using AlphaFold 3 (AF3)^53^. The experimentally observed TRB sequence for each target was modeled through the same workflow. The results were evaluated using TCRβ pTM, ipTM, mean pLDDT and interface predicted aligned error (iPAE). A candidate satisfied the composite AF3 structural-confidence criterion only if its predicted complex exceeded the matched experimentally observed TRB model in TCRβ pTM, ipTM and mean pLDDT while showing lower iPAE. Across the seven unseen pMHCs, 27.6% of OmniTCR(Base), 26.0% of OmniTCR(SFT) and 24.7% of OmniTCR(PMI) candidates met the composite criterion. The corresponding proportion was 19.9% for TCRdesign, the strongest specialized comparator (**Fig. 6a**).

**Fig. 6.**
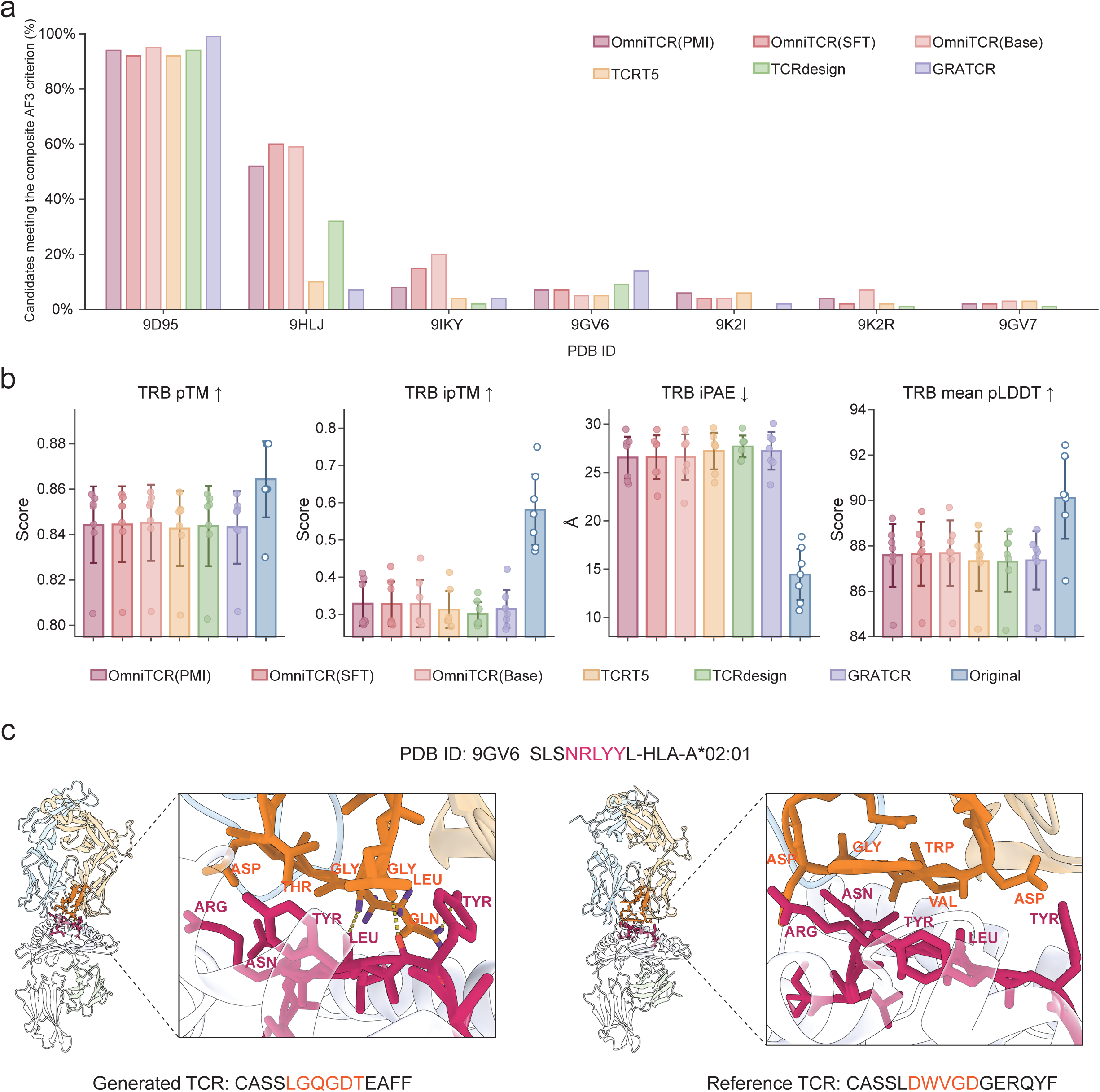
Predicted structural confidence of generated CDR3β candidates for unseen pMHCs. **a**, Percentage of 100 generated candidates per method that satisfied the composite AlphaFold 3 structural-confidence criterion for each of seven post-cutoff, epitope-excluded pMHCs. A candidate satisfied the criterion if its modeled complex showed higher TRB pTM, TRB ipTM and mean TRB pLDDT and lower TRB iPAE than the matched complex containing the experimentally observed TRB sequence. **b**, TRB structural-confidence metrics across the seven targets. Points denote target-level means; bars show mean ± s.d. across targets. Arrows indicate the favourable direction for each metric. **c**, AlphaFold 3 predicted peptide-CDR3β interfaces for the OmniTCR(PMI) reranked CASSLGQGDTEAFF candidate and the experimentally observed CASSLDWVGDGERQYF sequence in the SLSNRLYYL-HLA-A*02:01 complex (PDB ID: 9GV6). The generated or experimentally observed CDR3β loop is orange, and the peptide is magenta. Labels identify selected residues involved in predicted close contacts.

The individual AF3 metrics revealed complementary model strengths (**Fig. 6b**). OmniTCR(Base) achieved the highest mean TCRβ pTM and mean pLDDT, whereas OmniTCR(PMI) achieved the highest mean TCRβ ipTM and the lowest mean iPAE. The pretrained model thus favoured global TCRβ confidence, while PMI ranking enriched candidates with higher predicted interface confidence. For each pMHC and method, the highest TCRβ ipTM candidate from the 100 generated sequences was further compared with the matched experimentally observed sequence model (**Supplementary Fig. 6a**). OmniTCR(PMI) achieved the highest mean TCRβ ipTM among the generated methods (0.6086), compared with 0.5814 for the experimentally observed sequence models. It also exceeded the matched experimentally observed sequence in four of seven targets. OmniTCR(PMI) also achieved the lowest mean TCRβ iPAE among generated methods (16.55 Å; 17.41 Å for OmniTCR(SFT)), although the observed-sequence models had a lower mean iPAE overall (14.43 Å).

To illustrate how a sequence-divergent generated candidate could achieve high predicted interface confidence, the SLSNRLYYL-HLA-A*02:01 target (PDB ID: 9GV6) provided a representative example (**Fig. 6c**). The PMI-ranked CDR3β sequence, CASSLGQGDTEAFF, differed across most central positions from the experimentally observed CASSLDWVGDGERQYF sequence. In the AF3-predicted complex, Asp101 of the generated CDR3β was positioned within 2.78-2.83 Å of peptide Arg5, while Leu97 and Gln99 contacted peptide residues Leu6 and Tyr8. The experimentally observed CDR3β approached the same central peptide region through a different set of residues, including Asp98, Trp99, Val100 and Asp102. The sequence-divergent generated TRB can therefore retain a high-confidence predicted peptide-facing interface without reproducing the observed CDR3β sequence or its exact residue contacts.

## Discussion

OmniTCR integrates abundant TCR sequences and sparse multicomponent recognition records within a shared autoregressive foundation model. The resulting representation supports recognition prediction, cancer-associated repertoire classification and pMHC-conditioned receptor generation. The component-aware formulation provides a basis for future studies of antigen assignment, cross-reactivity assessment and receptor prioritization.

Previous efforts to unify TCR modeling have mainly shared information within closely related prediction tasks. UniPMT^34^ and UnifyImmun^35^ jointly model peptide-MHC presentation and TCR-pMHC recognition, and EPACT^43^ incorporates paired TCRα-TCRβ information and residue-level interactions. Repertoire classification and conditional TCR generation have also been developed with task-specific training objectives and data representations^36, 37, 39, 40, 54, 55^. OmniTCR differs in how information is shared. Rather than coupling specific downstream tasks on limited sets of annotated data, OmniTCR integrates diverse immune-sequence resources during pretraining, establishing a shared sequence representation before task-specific adaptation. This distinction is important. Models trained on limited TCR-pMHC recognition data may overfit to the specific features of a few well-studied peptides. In contrast, by integrating large receptor collections with multicomponent association records, OmniTCR establishes a shared representation that supports several prediction and generation tasks. This allows OmniTCR to generalize across multiple unseen scenarios without relying on exact training-set templates. This is evidenced across our evaluations, from predicting recognition on strictly epitope-disjoint benchmarks (**Fig. 3d**) and identifying cancer-associated repertoires across independent clinical cohorts (**Fig. 4c, d**), to generating candidate receptors with high AlphaFold 3 structural confidence for novel pMHCs (**Fig. 6a**).

Beyond benchmark performance, we also investigated to what extent a sequence-based model like OmniTCR can learn potential biological and structural properties of TCR recognition. At the receptor level, OmniTCR scores increased with clonal expansion and pMHC-multimer abundance (**Fig. 3e**), while model attention overlapped with experimentally resolved TCR-pMHC contact regions (**Fig. 3f-h**). In repertoire classification, OmniTCR captured sequence-level shifts distinguishing repertoire-derived candidate cancer-associated sequences and transferring to independently annotated tumour-antigen-associated receptors (**Fig. 4b, e**), and its repertoire scores reflected clinically meaningful disease stages and post-chemotherapy immune dynamics (**Fig. 4f, g**). For conditional receptor design, generated sequences preserved characteristic CDR3β patterns and broadly similar physicochemical distributions, while showing higher OLGA generation probabilities than reference sequences (**Fig. 5a-d, h**). Furthermore, perturbation analyses also revealed that sequence generation depended far more strongly on the peptide than on the MHC (**Fig. 5e, f**). Together, these analyses linked OmniTCR representations and predictions to recognition-associated sequence patterns, cellular measurements and predicted structural features.

The current scope is constrained by limited paired-chain annotations, MHC-II-associated recognition data and experimentally characterized negative TCR-pMHC interactions^14, 27, 30^. Because MHC-II molecules possess open-ended binding grooves accommodating variable peptide lengths, extending this framework to broader T-cell recognition will require richer paired datasets and tailored sequence representations. Furthermore, while AlphaFold 3 modeling supports the structural plausibility of generated receptors, prospective *in vitro* and *in vivo* experimental validation remains important to confirm the binding specificity and functional activity of generated TCRs. Looking forward, OmniTCR therefore provides a scalable sequence-based framework for studying antigen-specific TCR recognition and prioritizing candidate receptors for experimental validation.

## Methods

OmniTCR was designed to jointly learn from large TCR sequence collections and smaller TCR-pMHC association datasets containing different combinations of the peptide, MHC, TRA, and TRB. It uses a 113-million-parameter decoder-only Transformer and a component-aware sequence format that preserves component identity and boundaries, allowing records with different component combinations to be modeled within a common autoregressive input (**Fig. 1a**). Within this shared pretraining process, large TCR sequence collections provide broad coverage of receptor sequence organization, whereas multicomponent records provide the conditional relationships among peptide, MHC and TCR sequences. Because a causal decoder can condition only on preceding components, multicomponent records were also presented in complementary orders. This exposed the model to additional conditioning relationships without changing the native sequence of any component. The resulting pretrained backbone was then adapted to recognition prediction, cancer-associated repertoire classification, and pMHC-conditioned TRB sequence generation.

### Construction of the TCR-pMHC pretraining corpus

#### Data sources and molecular components

We assembled the pretraining corpus from human immune-sequence records available before 8 April 2025. Each record contained one or more of four molecular components: the peptide epitope, the major histocompatibility complex class I (MHC-I) molecule, the TCR*α* chain CDR3 sequence (TRA), and the TCR*β* chain CDR3 sequence (TRB). In humans, MHC molecules are encoded by the human leukocyte antigen (HLA) gene complex. The MHC-I component was therefore specified as an HLA-I allele, such as HLA-A*02:01. We restricted the present study to MHC-I because MHC-II-associated TCR records were substantially less abundant and require a distinct representation of peptide presentation. TRA and TRB sequences were collected primarily from TCRdb^28^ and iReceptor^26^. Multicomponent association records were obtained from IEDB^29^, VDJdb^27^, and McPAS-TCR^56^, together with datasets released with BigMHC^57^, NetMHCpan^31^, TCRAI^58^ and pMTnet^33^. Records from all sources were mapped to a common schema containing the peptide sequence, HLA-I allele, TRA CDR3 sequence, TRB CDR3 sequence, component combination, binding label when available, and data source. The components not reported by the original source were left empty.

#### Record standardization and sequence quality control

After mapping the collected records to the common schema, we standardized and filtered each component separately. Records were removed if a sequence required by the recorded component combination was missing or contained a stop codon or unsupported amino acid symbol. Peptide epitopes were restricted to 8-11 residues^59^. HLA-I allele names were standardized to two-field resolution, and records containing only serotype-level, antigen-group, or other lower-resolution HLA annotations were excluded. Each retained HLA-I allele was then mapped to the 34-residue HLA-I pseudo-sequence defined by NetMHCpan^60^, representing polymorphic positions in the MHC-I peptide-binding groove. The HLA-I allele specifies the biological MHC-I molecule, whereas its 34-residue pseudo-sequence provides the amino-acid input to OmniTCR. TRA CDR3 sequences were restricted to 7-20 residues and TRB CDR3 sequences to 7-24 residues. Both were required to begin with cysteine (C) and terminate with phenylalanine (F) or tryptophan (W)^61^. Sequences with incomplete CDR3 boundaries were excluded. Exact duplicate records were collapsed within each component combination after standardization. The single-component training partitions contained 434,143 unique peptides, 203 unique HLA-I pseudo-sequences, 5,095,003 unique TRA CDR3 sequences, and 319,410,970 unique TRB CDR3 sequences. Component-combination-specific record counts are reported in **Supplementary Section S1.1**.

#### Pretraining and held-out partitioning

We partitioned the quality-controlled records by component combination before sequence formatting or model training. Single-component peptide, HLA-I pseudo-sequence, TRA, and TRB records were assigned to autoregressive pretraining and validation without using associated disease or class labels. TRA-TRB records were partitioned in the same manner because no downstream evaluation was defined directly for this component combination. For component combinations used in downstream recognition prediction or TRB sequence generation, including peptide-MHC, peptide-TRB, peptide-MHC-TRA, peptide-MHC-TRB, and peptide-MHC-TRA-TRB, we reserved 5-9% of the records as held-out test partitions. These held-out records were excluded from both autoregressive pretraining and task-specific fine-tuning, and the remaining records were divided into pretraining and validation partitions. We additionally checked held-out associations across all training sources and component combinations using the task-specific matching procedures described in **Supplementary Sections S1.2** and **S1.4**. After partitioning, the canonical pretraining corpus contained 326,569,336 records across the component combinations. The component-order augmentation described below added 1,662,879 formatted sequences, yielding a final pretraining corpus of 328,232,215 sequences. Partition sizes for each component combination and the numbers of order-augmented sequences are reported in **Supplementary Tables S1** and **S2**, respectively.

### Component-aware sequence construction

#### Vocabulary and sequence-type tokens

OmniTCR uses amino-acid-level tokenization with a vocabulary of 34 tokens. The vocabulary contains five general special tokens, [*PAD*], [*BOS*], [*EOS*], [*UNK*], and [*MASK*] ; four sequence-type tokens, [*EPI*], [*HLA*], [*TRA*], and [*TRB*]; and 25 amino acid tokens, comprising the 20 standard amino acids and the extended residue symbols *X*, *B*, *Z*, *U*, and *O*. The [*PAD*] token is used for batch padding, whereas [*BOS*] and [*EOS*] mark the beginning and end of each formatted record. The [*UNK*] token is reserved for residues outside the defined vocabulary, and the [*MASK*] token is retained in the vocabulary. The four sequence-type tokens identify the peptide epitope, the HLA-I pseudo-sequence (MHC-I component), the TRA CDR3 sequence, and the TRB CDR3 sequence, respectively. Each sequence-type token is placed before and after its corresponding amino acid sequence, thereby identifying both the component type and its boundaries.

#### Component-aware sequence format

Let *E*, *M*, *A*, and *B* denote the amino acid sequences of the peptide epitope, HLA-I pseudo-sequence (MHC-I component), TRA CDR3, and TRB CDR3, respectively. Using the sequence-type tokens defined above, each component is represented as:

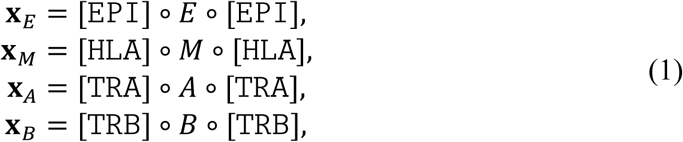

where o denotes sequence concatenation. Components present in a record are arranged according to the canonical order peptide-MHC-TRA-TRB, and components not reported in the original record are omitted with their sequence-type tokens. A record containing *k* observed components is represented as:

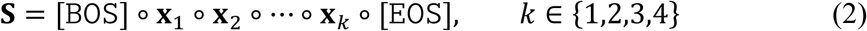

For example, a complete peptide-MHC-TRA-TRB record contains the ordered components (**x***_E_*, **x***_M_*, **x***_A_*, **x***_B_*). This format allows records with different component combinations to be represented by the same model, while preserving the identity and boundaries of the components (**Fig. 1a**).

#### Component-order augmentation

Because OmniTCR uses causal self-attention, each token is predicted only from the tokens that precede it. The order of the components determines which components serve as context for those that follow. To expose the model to complementary conditioning contexts, we retained the canonical order for each record and added one or more alternative component orders for multicomponent records. For records containing two or three components, the alternative order reverses the component order. Four-component records receive two complementary orders, allowing either TRA or TRB to provide the initial context while the other TCR chain appears after the pMHC components. The additional orders were:

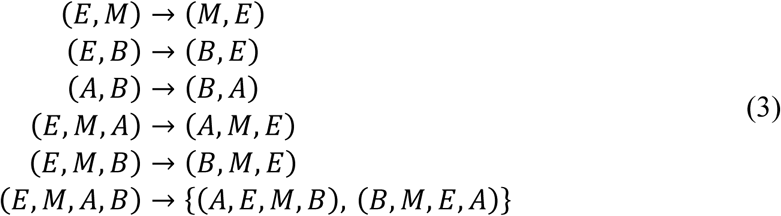

The arrows denote additional formatted sequences from the canonical order. Only the order of complete molecular components was changed; the amino acid sequence within each component retained its native N-to-C orientation. The numbers of order-augmented sequences for each component combination are reported in **Supplementary Table S2**.

### OmniTCR architecture and autoregressive pretraining

#### Model architecture

OmniTCR is a 113-million-parameter, 12-layer decoder-only Transformer^62^ (**Fig. 1b**). It consists of a token embedding layer, a stack of decoder-only Transformer blocks, and a language-model head. Each Transformer block uses RMSNorm^63^, causal self-attention, and a SwiGLU^64^ feed-forward network. A causal attention mask prevents each position from accessing subsequent tokens, and rotary positional embeddings^65^ are applied to the query and key representations to encode positional information. The language-model head projects each final hidden state to logits over the 34-token vocabulary. OmniTCR receives the component-aware sequence format described above, allowing records containing one to four components to be processed without component-specific architectural changes. The complete architecture, including the hidden dimension, number of attention heads, and feed-forward dimension, is provided in **Supplementary Section S2.1**.

#### Causal next-token prediction

OmniTCR was pretrained by predicting each token from the tokens that precede it in the formatted record. Given a tokenized sequence **x** = (*x*_0_, *x*_1_, …, *x_T_*), where *x*_0_ is [*BOS*], the model factorizes the probability of the remaining sequence as:

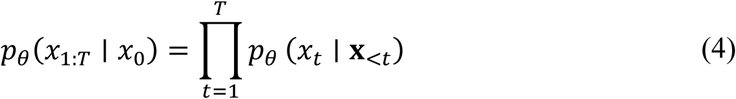

where **x**_<*t*_ = (*x*_0_, …, *x_t_*_−1_) and *θ* denotes the model parameters. This autoregressive pretraining minimizes the mean negative log-likelihood over non-padding target positions:

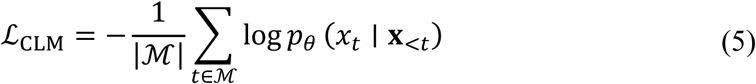

where ℳ is the set of target positions included in the loss. Amino acid tokens, sequence-type tokens, and the final [*EOS*] token are all prediction targets. The objective therefore learns sequence regularities within individual components while also capturing transitions across component boundaries and conditional relationships among components that occur in the same record. All model parameters were optimized during pretraining. The optimizer, learning-rate schedule, batch configuration and distributed-training strategy are reported in **Supplementary Table S6**.

### Downstream task adaptation

All downstream applications were initialized from the pretrained OmniTCR backbone, OmniTCR(Base). For supervised recognition prediction and cancer-associated repertoire classification, task-specific classification heads were attached to representations extracted from the pretrained backbone, with the adaptation strategy described in the corresponding sections below. pMHC-conditioned TRB sequence generation used supervised autoregressive fine-tuning followed, where indicated, by training-free PMI-inspired candidate reranking (**Fig. 1c**).

#### Recognition prediction

The component-aware sequence format allowed the same pretrained OmniTCR to be applied to different recognition tasks defined by the components present in the input: peptide-MHC presentation (P-M), peptide-TRB recognition (P-T), and peptide-MHC-TRB interaction prediction (P-M-T). The same backbone and downstream classifier architecture were used across these tasks, with each input formatted according to its corresponding component combination.

We first used the pretrained backbone directly, without a classification head or task-specific fine-tuning. This setting, denoted OmniTCR(Base), assigns a perplexity to each formatted molecular record:

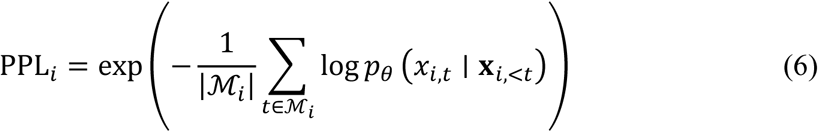

where ℳ*_i_* contains all non-padding target positions following the [*BOS*] token. A lower perplexity indicates that the formatted combination has higher probability under the pretrained model. We therefore used −PPL*_i_* as the prediction score, so that a higher score indicates a more strongly supported combination.

For supervised prediction, we added a two-layer multilayer perceptron classification head to the pretrained backbone. The input record was processed in the same component-aware sequence format, and the final-layer hidden state at the closing sequence-type token of the last molecular component was used as the record representation. This hidden state summarizes all preceding components and their sequence context. The representation was passed through the classification head with a GELU activation and a sigmoid output to obtain the predicted probability. Separate models were trained for P-M, P-T, and P-M-T using binary cross-entropy loss. Both the backbone and classification head were updated during training, yielding OmniTCR(FFT). For the paired-chain analysis, the TRA CDR3 sequence was added to the P-M-T input while retaining the same backbone and classification head. Model dimensions and optimization hyperparameters are provided in **Supplementary Table S7**.

Each OmniTCR(FFT) model was trained using supported records from the corresponding training partition and an equal number of task-specific negative records. P-M negatives were generated by recombining peptide epitopes and HLA-I alleles after removing known peptide-MHC associations. P-T and P-M-T negatives were obtained from experimentally annotated non-binding records in TRAIT^30^. Up to 100,000 supported records were used to train each task. Performance was assessed in three evaluation settings. Internal evaluation used task-specific records held out from both autoregressive pretraining and fine-tuning. External evaluation used independent benchmarks from NetMHCpan^31^, TRAIT^30^, and SARS-CoV-2^41^ datasets. Unseen-epitope evaluation used epitope-disjoint test sets in which every test peptide and all records containing that peptide were excluded from both pretraining and fine-tuning. Overlap with OmniTCR training data was assessed by matching the components required by each task across all sources and component combinations. Evaluation records were excluded if the corresponding association occurred in any training record, including records containing additional components (**Supplementary Section S1.2**). Overlaps with publicly available comparator training data were also removed. Dataset compositions are provided in **Supplementary Tables S3** and **S4**.

#### Cancer-associated repertoire classification

The same pretrained backbone was adapted to cancer-associated repertoire classification using individual TRB sequences as inputs. Each TRB CDR3 sequence was formatted as a single-component record using the component-aware sequence format. The final-layer hidden state at the closing [*TRB*] token was used as the TRB representation. This representation was passed through a two-layer multilayer perceptron with ReLU activation, followed by a softmax output over the candidate cancer-associated and healthy-background classes. The classifier was optimized using cross-entropy loss. We evaluated two adaptation strategies: OmniTCR(FT), which froze the pretrained backbone and updated only the classification head, and OmniTCR(FFT), which updated both the backbone and classification head. Complete optimization settings are provided in **Supplementary Table S7**.

To aggregate receptor-level predictions into a repertoire-level cancer score, productive TRB clonotypes passing quality control were ranked by clone frequency, and each was assigned a cancer-associated probability by OmniTCR. The OmniTCR cancer-associated TCR enrichment score (OmniTCR-CES) was defined as the median predicted probability among the 1,000 most abundant retained clonotypes in each repertoire, or among all retained clonotypes when fewer than 1,000 were available.

Receptor-level training data were constructed from 6,078 cancer and 1,504 healthy repertoires obtained from TCRdb 2.0. For the healthy-background pool, the 10,000 most abundant productive TRB clonotypes from each healthy repertoire were merged and deduplicated to form a healthy reference library. For the candidate cancer-associated pool, TRB sequences from the cancer repertoires were clustered using GLIPH2^47^. Sequences that also occurred in the healthy reference library were removed, leaving 1,489,111 unique candidate cancer-associated TRB sequences. To construct the binary classification dataset, an equal number of healthy-background sequences were sampled from the healthy reference library while matching the clonotype-frequency distribution of the candidate cancer-associated sequences. Detailed data-construction procedures are provided in **Supplementary Section S1.3**. We evaluated the receptor-level classifier directly at the receptor level and, after aggregation using OmniTCR-CES, at the repertoire level. Receptor-level transfer was evaluated using an independently annotated VDJdb^27^ collection of cancer-antigen-associated TRB sequences. Internal repertoire evaluation used 53 cancer and 43 healthy samples held out before training. External repertoire evaluation included a multiregional NSCLC cohort^48^ and pan-cancer, early-stage cancer, and immune-control cohorts from DeepCAT^36^. Treatment-associated changes in TCR repertoires were further examined using the OSLO-COMET cohort^49^. Complete cohort compositions are provided in **Supplementary Table S5**.

#### pMHC-conditioned TRB sequence generation

We formulated pMHC-conditioned TRB CDR3 generation as a conditional autoregressive language-modeling task. OmniTCR(SFT) was initialized from the pretrained OmniTCR backbone and adapted by supervised fine-tuning (SFT) on supported peptide-MHC-TRB associations. Each training sequence contained a pMHC prompt followed by the target TRB sequence:

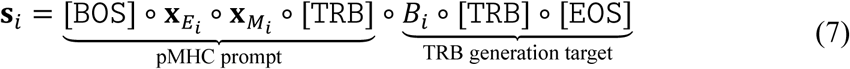

where **x***_Ei_* and **x***_Mi_* are the formatted peptide and HLA-I pseudo-sequence components, and *B_i_* is the target TRB CDR3 sequence. The opening [*TRB*] token specifies that the following component is TRB, whereas the closing [*TRB*] and [*EOS*] tokens terminate the TRB component and the complete record, respectively. During both training and inference, each TRB residue was conditioned on the pMHC prompt and the preceding TRB residues. The supervised fine-tuning loss was calculated only for the target TRB residues, the closing [*TRB*] token and the final [*EOS*] token. The pMHC prompt was excluded from the loss.

Standard supervised fine-tuning assigns equal weight to every target residue. This does not reflect the position-dependent organization of CDR3*β*. Its terminal regions contain more conserved sequence patterns, whereas its central junction is more variable and frequently contributes to peptide-facing contacts. We therefore increased the contribution of central residues using a parabolic position-weighting scheme. For a CDR3*β* sequence of length *L_i_*, let *r_i_*_,*t*_ ∈ {0, . . ., *L_i_* − 1} denote the relative position of residue *t*. The position weight was defined as:

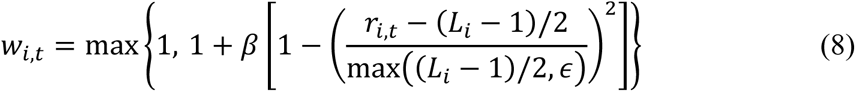

where *ε* = 10^−8^ and *β* = 3. Weights were 1 at the termini and increased towards an upper bound of 4 near the sequence midpoint. The closing [*TRB*] and [*EOS*] tokens retain unit weight. We further added a five-residue window loss calculated over all overlapping five-residue windows within the CDR3*β* sequence. This term increases the contribution of residues occurring within contiguous local sequence segments and complements the position-weighted token loss. The position-weighted and window-based losses were combined using a relative coefficient of *α* = 3.5. The full objective and optimization settings are provided in **Supplementary Section S2.3** and **Table S8**; the ablation is reported in **Section S6.4** and **Table S9**.

At inference, beam search may favor TRB sequences that have high likelihood even without peptide-MHC context. To reduce this pMHC-independent preference, we developed OmniTCR(PMI), which reranks candidates generated by OmniTCR(SFT) without additional training. For a candidate TRB sequence *y*, let l*_θ_*(*y* ∣ **x**) denote the mean log-likelihood of its CDR3*β* residues and closing [*TRB*] token under prompt *x*. The PMI-inspired score was defined as:

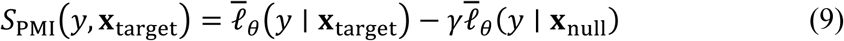

where *γ* = 0.8, **x**_target_ denotes the target pMHC prompt and:

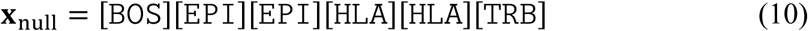

The null prompt retained empty peptide and HLA components and the opening [TRB] token, while omitting peptide and HLA-I residues. Subtracting the null-prompt likelihood reduces the score of candidates that are strongly supported by the general TRB sequence distribution even in the absence of pMHC information, while favoring candidates whose likelihood increases under the target pMHC context. More modeling settings are provided in **Supplementary Sections S2.4**.

OmniTCR(SFT) was trained on supported peptide-MHC-TRB records from the generation training partition described above. The SFT dataset combined records from the curated generation-training partition with additional TCRT5-derived records.

Sequence-level generation performance was evaluated in three evaluation settings. Internal evaluation used P-M-T records constructed from our data collection excluded from both autoregressive pretraining and supervised fine-tuning. External evaluation used retained TRAIT records, whereas unseen-pMHC evaluation used exact peptide– HLA pairs absent from pretraining and generation SFT. All these records were also completely excluded from both autoregressive pretraining and supervised fine-tuning. Within each setting, the 20 pMHCs with the largest numbers of experimentally observed TRB sequences were selected as generation targets. Each method generated 100 valid TRB CDR3 sequences per target under the same output budget. Structural evaluation used seven experimentally determined TCR-pMHC complexes from the TCR3d^52^ database, released after the 8 April 2025 pretraining cutoff: PDB ID 9GV6, 9GV7, 9IKY, 9HLJ, 9D95, 9K2R, and 9K2I. Complexes containing peptides present in any OmniTCR training and fine-tuning data were excluded. For each retained pMHC, 100 generated TRB CDR3 sequences from each method were inserted into the corresponding TRB framework and modeled with the experimentally reported TRA, peptide, MHC-I heavy chain, and *β*_2_-microglobulin using AlphaFold 3^53^. Complete dataset compositions are provided in **Supplementary Sections S1.4**.

### Evaluation

Recognition prediction was evaluated using the area under the receiver operating characteristic curve (AUROC) and the area under the precision-recall curve (AUPRC). Positive predictive value at rank *k* (PPV@*k*) was additionally used to quantify the proportion of experimentally supported records among the top-*k* predictions. Cancer-associated repertoire classification was evaluated using AUROC and AUPRC calculated from the repertoire-level scores produced by each method.

Generated TRB sequences were evaluated for sequence validity, CDR3*β* length, amino acid usage, OLGA^50^ generation probability, and physicochemical properties. Their similarity to experimentally observed reference repertoires was evaluated using Char-BLEU, sequence recovery, and exact-match F1@100, while output diversity was summarized by the number of unique generated sequences and pairwise Jaccard similarity across target pMHCs.

The structural confidence of the generated TRB sequences was evaluated using TRB pTM, TRB ipTM, mean TRB pLDDT, and TRB interface predicted aligned error (iPAE) from AlphaFold 3. Higher pTM, ipTM, and pLDDT and lower iPAE indicate greater predicted structural confidence. A generated complex satisfied the composite AlphaFold 3 structural-confidence criterion only when it simultaneously achieved higher TRB pTM, ipTM, and pLDDT and lower TRB iPAE than the matched complex modeled with the experimentally observed TRB sequence. The specific details of these evaluation procedures are provided in **Supplementary Section S3**.

## Data availability

Sequence collection and pretraining used publicly available data from TCRdb2.0 [https://guolab.wchscu.cn/TCRdb2/], iReceptor [https://gateway.ireceptor.org/], IEDB [https://www.iedb.org/], VDJdb [https://vdjdb.com/], McPAS-TCR [https://friedmanlab.weizmann.ac.il/McPAS-TCR/]. Additional association data were obtained from resources released with BigMHC [https://data.mendeley.com/datasets/dvmz6pkzvb/4], NetMHCpan-4.1 [https://services.healthtech.dtu.dk/services/NetMHCpan-4.1/], TCRAI [https://github.com/regeneron-mpds/TCRAI/tree/main/data] and pMTnet [https://github.com/tianshilu/pMTnet].

Raw SARS-CoV-2 sequencing data from Minervina et al. are available under BioProject accession PRJNA744851 [https://www.ncbi.nlm.nih.gov/bioproject/?term=PRJNA744851]. The TRAIT test datasets were obtained from TRAIT database [https://pgx.zju.edu.cn/traitdb]. Antigen-specific single-cell TCR data were obtained from the 10x Genomics CD8+ T-cell datasets for healthy donors 1, 2, 3, 4 [https://www.10xgenomics.com/datasets/cd-8-plus-t-cells-of-healthy-donor-1-1-standard-3-0-2] [https://www.10xgenomics.com/datasets/cd-8-plus-t-cells-of-healthy-donor-2-1-standard-3-0-2] [https://www.10xgenomics.com/datasets/cd-8-plus-t-cells-of-healthy-donor-3-1-standard-3-0-2] [https://www.10xgenomics.com/datasets/cd-8-plus-t-cells-of-healthy-donor-4-1-standard-3-0-2].

Cancer and healthy repertoire data for classifier training and internal evaluation were also obtained from TCRdb 2.0. External cancer repertoire cohorts benchmarks were obtained from immuneACCESS [https://clients.adaptivebiotech.com/immuneaccess]. The associated cancer, healthy and immune-control benchmark data are available in the DeepCAT dataset [https://doi.org/10.5281/zenodo.3894880]. The independent NSCLC repertoire dataset is available under BioProject accession PRJNA506151 [https://www.ncbi.nlm.nih.gov/bioproject/506151]. The OSLO-COMET repertoire data are available through immuneACCESS [https://clients.adaptivebiotech.com/pub/hoye-2023-gs]. The receptor-level transfer analysis used tumour-antigen-associated TRB annotations from VDJdb.

Experimental structures were obtained from TCR3d [https://tcrdb.ibbr.umd.edu/] and the RCSB Protein Data Bank [https://www.rcsb.org/] with PDB ID: 7N6E, 9GV6, 9GV7, 9IKY, 9HLJ, 9D95, 9K2R and 9K2I.

More details of dataset composition and processing procedures are described in the Methods and **Supplementary Sections S1.1–S1.4**.

## Code availability

The source code is available at [https://github.com/GuoBioinfoLab/OmniTCR]. The pretrained OmniTCR model, tokenizer and task-specific checkpoints for recognition prediction, repertoire analysis and pMHC-conditioned TCRβ generation are available through Hugging Face [https://huggingface.co/loveCloud/OmniTCR]. The repository includes the software-environment specification, example input files and instructions for reproducing the principal inference workflows.

## Author contributions

F.Z., D.F. and A.G. conceived the project. F.Z. and D.F. designed the method. F.Z., D.F. and L.D. designed and conducted the numerical experiments. F.Z., D.S. and D.F. visualized and illustrated the manuscript figures. F.Z. and D.F. wrote the manuscript. A.G. and W.L. supervised the study and provided support throughout the project. D.S., L.D., Z.T. and Q.L. provided technical and infrastructure guidance. All authors discussed the results and contributed to the final manuscript. All authors approved the manuscript.

## References

1. Davis, M.M. & Bjorkman, P.J. T-Cell Antigen Receptor Genes and T-Cell Recognition. Nature 334, 395–402 (1988).

2. Rossjohn, J. et al. T Cell Antigen Receptor Recognition of Antigen-Presenting Molecules. Annu Rev Immunol 33, 169–200 (2015).

3. Kearse, K.P., Roberts, J.P., Wiest, D.L. & Singer, A. Developmental Regulation of Alpha-Beta-T-Cell Antigen Receptor Assembly in Immature Cd4(+)Cd8(+) Thymocytes. Bioessays 17, 1049–1054 (1995).

4. Pishesha, N., Harmand, T.J. & Ploegh, H.L. A guide to antigen processing and presentation. Nat Rev Immunol 22, 751–764 (2022).

5. Emerson, R.O. et al. Immunosequencing identifies signatures of cytomegalovirus exposure history and HLA-mediated effects on the T cell repertoire. Nat Genet 49, 659–665 (2017).

6. Dewitt, W.S. et al. Human T cell receptor occurrence patterns encode immune history, genetic background, and receptor specificity. Elife 7, e38358 (2018).

7. Foy, S.P. et al. Non-viral precision T cell receptor replacement for personalized cell therapy. Nature 615, 687–696 (2023).

8. Gubin, M.M. et al. Checkpoint blockade cancer immunotherapy targets tumour-specific mutant antigens. Nature 515, 577–581 (2014).

9. Tran, E. et al. Cancer Immunotherapy Based on Mutation-Specific CD4+T Cells in a Patient with Epithelial Cancer. Science 344, 641–645 (2014).

10. Zaslavsky, M.E. et al. Disease diagnostics using machine learning of B cell and T cell receptor sequences. Science 387, eadp2407 (2025).

11. Nielsen, S.C.A. & Boyd, S.D. Human adaptive immune receptor repertoire analysis-Past, present, and future. Immunol Rev 284, 9–23 (2018).

12. Mason, D. A very high level of crossreactivity is an essential feature of the T-cell receptor. Immunol Today 19, 395–404 (1998).

13. Wooldridge, L. et al. A Single Autoimmune T Cell Receptor Recognizes More Than a Million Different Peptides. J Biol Chem 287, 1168–1177 (2012).

14. Gfeller, D., Racle, J., Harari, A. & Croce, G. Advances in predicting T cell epitope recognition for cancer immunotherapy. Nat Cancer 7, 711–722 (2026).

15. Arstila, T.P. et al. A direct estimate of the human αβ T cell receptor diversity. Science 286, 958–961 (1999).

16. Alt, F.W., et al. Vdj Recombination. Immunol Today 13, 306–314 (1992).

17. Krangel, M.S. Mechanics of T cell receptor gene rearrangement. Curr Opin Immunol 21, 133–139 (2009).

18. Robins, H.S. et al. Comprehensive assessment of T-cell receptor β-chain diversity in αβ T cells. Blood 114, 4099–4107 (2009).

19. Falk, K., Rotzschke, O., Stevanovic, S., Jung, G. & Rammensee, H.G. Allele-Specific Motifs Revealed by Sequencing of Self-Peptides Eluted from Mhc Molecules. Nature 351, 290–296 (1991).

20. Yewdell, J.T.W. & Bennink, J.R. Immunodominance in major histocompatibility complex class I-restricted T lymphocyte responses. Annual Review of Immunology 17, 51–88 (1999).

21. Dash, P. et al. Quantifiable predictive features define epitope-specific T cell receptor repertoires. Nature 547, 89–93 (2017).

22. Glanville, J. et al. Identifying specificity groups in the T cell receptor repertoire. Nature 547, 94–98 (2017).

23. Sewell, A.K. Why must T cells be cross-reactive? Nat Rev Immunol 12, 668–677 (2012).

24. Zhang, S.Q. et al. High-throughput determination of the antigen specificities of T cell receptors in single cells. Nat Biotechnol 36, 1156–1159 (2018).

25. Pai, J.A. & Satpathy, A.T. High-throughput and single-cell T cell receptor sequencing technologies. Nat Methods 18, 881–892 (2021).

26. Corrie, B.D. et al. iReceptor: A platform for querying and analyzing antibody/B-cell and T-cell receptor repertoire data across federated repositories. Immunol Rev 284, 24–41 (2018).

27. Shugay, M. et al. VDJdb: a curated database of T-cell receptor sequences with known antigen specificity. Nucleic Acids Res 46, D419–D427 (2018).

28. Yue, T. et al. TCRdb 2.0: an updated T-cell receptor sequence database. Nucleic Acids Res 54, D504–D510 (2026).

29. Vita, R. et al. The Immune Epitope Database (IEDB): 2018 update. Nucleic Acids Res 47, D339–D343 (2019).

30. Wei, M.M. et al. TRAIT: A Comprehensive Database for T-cell Receptor-antigen Interactions. Genom Proteom Bioinf 23, qzaf033 (2025).

31. Reynisson, B., Alvarez, B., Paul, S., Peters, B. & Nielsen, M. NetMHCpan-4.1 and NetMHCIIpan-4.0: improved predictions of MHC antigen presentation by concurrent motif deconvolution and integration of MS MHC eluted ligand data. Nucleic Acids Res 48, W449–W454 (2020).

32. Peng, X.A. et al. Characterizing the interaction conformation between T-cell receptors and epitopes with deep learning. Nat Mach Intell 5, 395–407 (2023).

33. Lu, T.S. et al. Deep learning-based prediction of the T cell receptor-antigen binding specificity. Nat Mach Intell 3, 864–875 (2021).

34. Zhao, Y.X. et al. A unified deep framework for peptide-major histocompatibility complex-T cell receptor binding prediction. Nat Mach Intell 7, 650–660 (2025).

35. Yu, C.P., Fang, X., Tian, S.Y. & Liu, H. A unified cross-attention model for predicting antigen binding specificity to both HLA and TCR molecules. Nat Mach Intell 7, 278–292 (2025).

36. Beshnova, D. et al. De novo prediction of cancer-associated T cell receptors for noninvasive cancer detection. Sci Transl Med 12, eaaz3738 (2020).

37. Cai, Y.D. et al. The Deep Learning Framework iCanTCR Enables Early Cancer Detection Using the T-cell Receptor Repertoire in Peripheral Blood. Cancer Res 84, 1915–1928 (2024).

38. Wu, K. et al. TCR-BERT: learning the grammar of T-cell receptors for flexible antigen-binding analyses. Pr Mach Learn Res 240, pp. 194–229 (2024).

39. Li, X.K. et al. TCRdesign: an antigen-specific generative language model for de novo design of T-cell receptors. Brief Bioinform 26, bbaf691 (2025).

40. Karthikeyan, D., Bennett, S.N., Reynolds, A.G., Vincent, B.G. & Rubinsteyn, A. Conditional generation of real antigen-specific T cell receptor sequences. Nat Mach Intell 7, 1494–1509 (2025).

41. Minervina, A.A. et al. SARS-CoV-2 antigen exposure history shapes phenotypes and specificity of memory CD8+ T cells. Nat Immunol 23, 781–790 (2022).

42. Nilsson, J.B., Greenbaum, J., Peters, B. & Nielsen, M. NetMHCpan-4.2: improved prediction of CD8+epitopes by use of transfer learning and structural features. Front Immunol 16, 1616113 (2025).

43. Zhang, Y.M. et al. Epitope-anchored contrastive transfer learning for paired CD8+ T cell receptor-antigen recognition. Nat Mach Intell 6, 1344–1358 (2024).

44. 10x Genomics, Edn. Cell Ranger v3.0.2 (10x Genomics, 2019).

45. Chaurasia, P. et al. Structural basis of biased T cell receptor recognition of an immunodominant HLA-A2 epitope of the SARS-CoV-2 spike protein. J Biol Chem 297 (2021).

46. Schymkowitz, J. et al. The FoldX web server: an online force field. Nucleic Acids Res 33, W382–W388 (2005).

47. Huang, H., Wang, C.L., Rubelt, F., Scriba, T.J. & Davis, M.M. Analyzing the Mycobacterium tuberculosis immune response by T-cell receptor clustering with GLIPH2 and genome-wide antigen screening. Nat Biotechnol 38, 1194–1202 (2020).

48. Jia, Q.Z. et al. Local mutational diversity drives intratumoral immune heterogeneity in non-small cell lung cancer. Nat Commun 9, 5361 (2018).

49. Hoye, E. et al. T cell receptor repertoire sequencing reveals chemotherapy-driven clonal expansion in colorectal liver metastases. Gigascience 12, giad032 (2023).

50. Sethna, Z., Elhanati, Y., Callan, C.G., Jr., Walczak, A.M. & Mora, T. OLGA: fast computation of generation probabilities of B- and T-cell receptor amino acid sequences and motifs. Bioinformatics 35, 2974–2981 (2019).

51. Zhou, Z.H. et al. GRATCR: Epitope-Specific T Cell Receptor Sequence Generation With Data-Efficient Pre-Trained Models. Ieee J Biomed Health 29, 2271–2283 (2025).

52. Lin, V.L.R. et al. TCR3d 2.0: expanding the T cell receptor structure database with new structures, tools and interactions. Nucleic Acids Res 53, D604–D608 (2025).

53. Abramson, J. et al. Accurate structure prediction of biomolecular interactions with AlphaFold 3. Nature 630, 493–500 (2024).

54. Sidhom, J.W., Larman, H.B., Pardoll, D.M. & Baras, A.S. DeepTCR is a deep learning framework for revealing sequence concepts within T-cell repertoires. Nat Commun 12, 1605 (2021).

55. Zhang, M. et al. BertTCR: a Bert-based deep learning framework for predicting cancer-related immune status based on T cell receptor repertoire. Brief Bioinform 25, bbae420 (2024).

56. Tickotsky, N., Sagiv, T., Prilusky, J., Shifrut, E. & Friedman, N. McPAS-TCR: a manually curated catalogue of pathology-associated T cell receptor sequences. Bioinformatics 33, 2924–2929 (2017).

57. Albert, B.A. et al. Deep neural networks predict class I major histocompatibility complex epitope presentation and transfer learn neoepitope immunogenicity. Nat Mach Intell 5, 861–872 (2023).

58. Zhang, W. et al. A framework for highly multiplexed dextramer mapping and prediction of T cell receptor sequences to antigen specificity. Sci Adv 7, eabf5835 (2021).

59. Momburg, F., Roelse, J., Hammerling, G.J. & Neefjes, J.J. Peptide Size Selection by the Major Histocompatibility Complex-Encoded Peptide Transporter. J Exp Med 179, 1613–1623 (1994).

60. Hoof, I. et al. NetMHCpan, a method for MHC class I binding prediction beyond humans. Immunogenetics 61, 1–13 (2009).

61. Fang, H. et al. Quantitative T cell repertoire analysis by deep cDNA sequencing of T cell receptor α and β chains using next-generation sequencing (NGS). Oncoimmunology 3, e968467 (2014).

62. Vaswani, A., et al. Attention Is All You Need. Adv Neur In 30 (2017).

63. Zhang, B. & Sennrich, R. Root Mean Square Layer Normalization. Advances in Neural Information Processing Systems 32 (Nips 2019) 32 (2019).

64. Shazeer, N. GLU Variants Improve Transformer. arXiv preprint, arXiv:2002.05202 (2020).

65. Su, J.L. et al. RoFormer: Enhanced transformer with Rotary Position Embedding. Neurocomputing 568, 127063 (2024).

