## Supplementary Information for "OmniTCR: a foundation model unifying T cell receptor recognition prediction and conditional sequence generation"

### S1. Data sources and dataset composition

#### S1.1 Construction of the pretraining corpus

The OmniTCR pretraining corpus integrates complementary sources of human HLA class I antigen presentation and TCR recognition. Large TCR sequence collections were obtained from TCRdb2.0[1] and iReceptor[2], whereas experimentally supported or previously curated multicomponent association records were assembled from IEDB[3], VDJdb[4] and McPAS-TCR[5] and from datasets distributed with BigMHC[6], NetMHCpan[7], TCRAI[8] and pMTnet[9]. Together, these sources contributed standalone peptides, HLA-I pseudo-sequences, TRA and TRB CDR3 sequences, as well as peptide-HLA, peptide-TRB, TRA-TRB, peptide-HLA-TRA, peptide-HLA-TRB and peptide-HLA-TRA-TRB records. After standardization and quality control (described in the main Methods), records were partitioned by component combination before sequence formatting. **Supplementary Table S1** reports 326,569,336 training records, 32,729 validation records and 87,005 held-out test records before component-order augmentation. Selected multicomponent training records were additionally formatted in complementary component orders. This procedure added 1,662,879 training sequences, yielding 328,232,215 formatted pretraining sequences. **Supplementary Table S2** reports the number added for each alternative component order.

To characterize the OmniTCR pretraining corpus, we further examined the abundance, annotation coverage and sequence composition of the assembled corpus. These analyses revealed pronounced imbalance across component combinations and antigen-receptor annotations, together with substantial heterogeneity in HLA-I coverage and sequence composition. These characteristics motivated the use of a shared foundation model with a component-aware sequence format, allowing records with different component combinations and levels of annotation completeness to be learned within a common autoregressive framework.

First, the large differences among multicomponent association records reflect the imbalance of currently available immune-sequence data. High-throughput repertoire sequencing provides extensive coverage of naturally occurring receptor sequences. By contrast, experimentally resolved chain pairing, peptide presentation and complete pMHC-TCR recognition remain comparatively limited.

This imbalance was also evident within the multicomponent association data. Among 2,271 peptides with annotated TRB sequences, the median number of unique TRB sequences associated with the peptide was four, whereas the 90th percentile was 390 (**Supplementary Fig. 1a**). A relatively small group of extensively characterized peptides accounted for a disproportionate fraction of known peptide-TRB associations. Retaining HLA context further reduced annotation density. Among 1,797 pMHCs, the median number of unique TRB sequences associated with the pMHC was two, and the 90th percentile was 17 (**Supplementary Fig. 1b**).

The same sparsity was apparent from the receptor perspective. Among 70,192 TRB sequences with a complete pMHC annotation, 91.2% were associated with one pMHC and 7.3% with two. Only 1.5% were associated with at least three pMHCs (**Supplementary Fig. 1c**). These distributions describe the coverage of published experiments and curated databases rather than intrinsic receptor specificity, because most TCRs have been tested against only a small fraction of possible pMHCs.

Additional analyses characterized HLA-I coverage and sequence-length composition within the pretraining corpus. Among the 20 HLA-I alleles with the greatest peptide coverage, the numbers of associated peptides and TRB sequences varied substantially, indicating that broad peptide-presentation coverage did not necessarily coincide with similarly broad TCR-recognition annotation (**Supplementary Fig. 1d**). Nine-residue peptides were the most common, whereas TRA and TRB CDR3 sequences were most frequently 14 and 15 residues long, respectively (**Supplementary Fig. 1e**). HLA-I annotation was also concentrated across peptides. Among 415,686 peptides with at least one annotated HLA-I association, 70.4% were associated with one HLA-I allele, 15.2% with two and 5.5% with three. Progressively fewer peptides were represented across four or more HLA-I alleles (**Supplementary Fig. 1f**). To visualize corpus-level residue composition, sequence logos were generated for the two most abundant CDR3 length classes in each receptor chain. These comprised 13- and 14-residue TRA CDR3 sequences and 15- and 16-residue TRB CDR3 sequences, with the number of contributing sequences reported above each logo (**Supplementary Fig. 1g, h**). These summarize corpus-level composition, including germline-derived and preprocessing-associated constraints.

**Table S1**. Record partitions before component-order augmentation

| Sequence type | Training | Validation | Test |
|---|---|---|---|
| Peptide | 434143 | 1086 | 0 |
| MHC | 203 | 203 | 0 |
| TRA | 5095003 | 12769 | 0 |
| TRB | 319410970 | 3195 | 0 |
| Peptide-MHC | 755645 | 8038 | 40190 |
| Peptide-TRB | 569294 | 6121 | 36726 |
| Peptide-MHC-TRA | 20906 | 224 | 1344 |
| Peptide-MHC-TRB | 65680 | 729 | 6561 |
| TRA-TRB | 183630 | 0 | 0 |
| Peptide-MHC-TRA-TRB | 33862 | 364 | 2184 |

**Table S2**. Additional formatted sequences generated by component-order augmentation

| Additional component order | Sequence count |
|---|---|
| MHC-Peptide | 755645 |
| TRB-Peptide | 569294 |
| TRA-MHC-Peptide | 20906 |
| TRB-MHC-Peptide | 65680 |
| TRB-TRA | 183630 |
| TRA-Peptide-MHC-TRB | 33862 |
| TRB-MHC-Peptide-TRA | 33862 |

#### S1.2 Recognition prediction datasets

Task-specific datasets for discriminative fine-tuning were constructed separately for the corresponding P-M, P-T, P-M-T and P-M-A-B tasks described in the main Methods. For each task, we randomly sampled up to 100,000 supported records from the corresponding pretraining data (**Supplementary Table S1**) to construct the fine-tuning dataset, using all available records when fewer were available. P-M negatives were generated by recombining peptides and HLA-I alleles after excluding combinations reported as supported associations. P-T and P-M-T negatives were obtained from experimentally annotated non-binding TRAIT records. All supported and negative records underwent the sequence-standardization and quality-control procedures used to construct the pretraining corpus (**Supplementary Section S1.1**). Duplicates were defined by the complete task-specific component combination and removed before model training.

For the primary OmniTCR(FFT) models, the P-M and P-T training sets each comprised 100,000 supported and 100,000 negative records. The processed P-M-T dataset contained 65,680 unique supported records, all of which were retained and combined with 65,680 negative records. The final P-M, P-T and P-M-T training sets therefore contained 200,000, 200,000 and 131,360 records, respectively.

For the data-efficiency analysis, the P-M and P-T models were trained using 1,000, 2,000, 5,000, 10,000, 20,000, 50,000 or 100,000 supported records sampled from the aforementioned training set, together with an equal number of task-specific negatives. P–M and P–T used up to 100,000 supported records, whereas P–M–T included the full set of 65,680 supported records.

Evaluation was conducted in four settings: internal, external, paired-chain and unseen-epitope evaluation (**Supplementary Tables S3** and **S4**).

Overlap was checked against the union of OmniTCR pretraining and task-specific fine-tuning records from all sources. We extracted peptide–HLA keys for P–M, peptide–TRB keys for P–T and peptide–HLA–TRB keys for P–M–T. Each key was extracted from every training record containing the required components, including records with additional components. For example, a peptide–TRB test pair was checked against peptide–MHC–TRB and peptide–MHC–TRA–TRB training records. Matching used standardized component values and was independent of data-source labels and formatted component order. Evaluation records with matching training keys were excluded from the test sets.

Internal evaluation used task-specific records held out from both autoregressive pretraining and discriminative fine-tuning. Supported records were obtained from the held-out P-M, P-T, and P-M-T partitions of the curated association corpus (**Supplementary Table S1**). P-M negatives were generated by recombining peptide epitopes and HLA-I alleles after removing known peptide-MHC associations. P-T and P-M-T negatives were obtained from experimentally annotated non-binding records in TRAIT[10]. Candidate records underwent the component-specific overlap checks against the OmniTCR training union described above. Overlaps with publicly available training data from applicable comparators, including UnifyImmun[11], pMTnet[9], and PanPep[12], were removed.

External evaluation used independently assembled recognition benchmarks from NetMHCpan[7], TRAIT[10] and SARS-CoV-2[13]. The P-M evaluation used the NetMHCpan dataset, which contained 2,014 supported and 4,028 negative records after processing. The TRAIT P-T dataset contained 2,285 supported and 4,557 experimentally annotated non-binding records, whereas the TRAIT P-M-T dataset contained 1,685 supported and 4,260 non-binding records. The SARS-CoV-2 evaluation used a separate one-to-one design in which negative TRB sequences were sampled from TCRdb rather than generated by rearranging antigen-responsive receptors within the same dataset. The resulting SARS-CoV-2 P-T and P-M-T datasets contained 3,235 supported and 3,235 negative records and 3,244 supported and 3,244 negative records, respectively. Records were standardized and deduplicated within each source dataset before applying the component-specific overlap checks described above. Overlaps with publicly available training data from applicable comparator methods were also removed.

Paired-chain evaluation examined whether the addition of TRA provided information beyond peptide, MHC-I and TRB alone. We selected 845 supported and 1,690 non-binding peptide-MHC-TRA-TRB records from TRAIT. Each record was evaluated in two matched input settings: the complete peptide-MHC-TRA-TRB input and a peptide-MHC-TRB input constructed by removing only the TRA component. The peptide, MHC-I, TRB, and recognition label therefore remained identical between the two settings, allowing the contribution of TRA to be evaluated through a record-level matched comparison.

Unseen-epitope evaluation assessed generalization to peptide epitopes absent from model training. For each P-M, P-T and P-M-T task, candidate unseen epitopes were further identified from supported records in the held-out test set from the internal dataset, TRAIT[10] and SARS-CoV-2[13]. An epitope was defined as unseen only if its sequence was absent from the complete pretraining and fine-tuning union. This check included peptide-only records and every multicomponent record containing a peptide. Supported test records containing these epitopes were then retained for unseen-epitope evaluation. P-M negatives were generated by recombining the retained test peptides and HLA-I alleles after removing known peptide-MHC associations, whereas P-T and P-M-T negatives were obtained from experimentally annotated non-binding TRAIT records containing the held-out peptides. The resulting P-M dataset contained 11,528 supported and 23,222 negative records, the P-T dataset contained 214 supported and 428 negative records, and the P-M-T dataset contained 47 supported and 94 negative records. These datasets were additionally subjected to the same standardization, deduplication, and overlap-control procedures used for the internal and external evaluations.

**Table S3**. Composition of internal and external recognition evaluation datasets

| Dataset | Task | Pos | Neg | Total |
|---|---|---|---|---|
| Internal | P-M | 28363 | 57143 | 85506 |
| Internal | P-T | 30611 | 61021 | 91632 |
| Internal | P-M-T | 3796 | 9866 | 13662 |
| NetMHCpan-4.1 | P-M | 2014 | 4028 | 6042 |
| TRAIT | P-T | 2285 | 4557 | 6842 |
| TRAIT | P-M-T | 1685 | 4260 | 5945 |
| TRAIT $\alpha - \beta$ Paired | P-M-T | 845 | 1690 | 2535 |
| TRAIT $\alpha - \beta$ Paired | P-M-TRA-TRB | 845 | 1690 | 2535 |
| SARS-CoV-2 | P-T | 3235 | 3235 | 6470 |
| SARS-CoV-2 | P-M-T | 3244 | 3244 | 6488 |

**Table S4**. Unseen-epitope dataset composition

| Task | Pos | Neg | Total |
|---|---|---|---|
| P-M | 11528 | 23222 | 34750 |
| P-T | 214 | 428 | 642 |
| P-M-T | 47 | 94 | 141 |

#### S1.3 Cancer-associated TCR and pan-cancer repertoire datasets

We collected 7,598 cancer and 1,881 healthy TRB repertoires from TCRdb2.0, which further provides cancer and healthy annotations beyond the pretraining data. Before pooling any TRB sequences, we partitioned the repertoires at the repertoire level into training, validation and test sets in an 8:1:1 ratio. The resulting training set contained 6,078 cancer and 1,504 healthy repertoires. Two receptor-level training pools were then constructed from the training repertoires. For the healthy-background pool, the 10,000 most abundant productive TRB clonotypes from each healthy repertoire were merged and deduplicated, yielding a healthy reference library of 12,364,886 unique TRB CDR3 sequences. For the cancer-associated pool, TRB sequences from the cancer repertoires were grouped using GLIPH2[14]. Clusters were retained when they satisfied $P_{\text{Fisher}} < 1 \times 10^{-4}$, $P_{\text{V}\beta} < 0.05$ and they contained at least four unique CDR3β sequences. Cluster-associated sequences present in the healthy reference library were then removed. Cancer repertoires were eligible only if more than 100 candidate cancer-associated CDR3β sequences remained after this exclusion. Sequences from eligible repertoires were pooled and deduplicated, yielding 1,489,111 unique candidate cancer-associated CDR3β sequences. An equal number of healthy-background sequences were then sampled from the healthy reference library while matching the clone-frequency distribution of the candidate cancer-associated sequences.

Cancer and healthy evaluation repertoires were processed using the same GLIPH2[14] cluster-quality criteria. For quality control and sufficient repertoire depth, only repertoires with more than 50 retained TRB clonotypes were included.

Internal evaluation assessed whether aggregated receptor-level probabilities distinguished cancer and healthy repertoires held out before classifier training. The internal test set contained 53 cancer and 43 healthy repertoires that had been separated before construction of the receptor-level training dataset. For each repertoire, receptor-level cancer-associated probabilities were aggregated using the OmniTCR cancer-associated TCR enrichment score (OmniTCR-CES) defined in the main Methods.

External evaluation assessed generalization to independently collected cancer repertoires. The multiregional non-small-cell lung cancer (NSCLC) cohort[15] initially contained 57 tumour-region repertoires from 15 patients, of which 43 passed repertoire-processing and quality-control criteria. These tumour-region repertoires were compared with 40 healthy-control repertoires obtained from held-out healthy repertoires. Because several patients contributed more than one tumour-region repertoire, each repertoire was treated as the analysis unit. Additional pan-cancer, early-stage cancer, and non-cancer immune-control cohorts were obtained from projects distributed with DeepCAT[16]. The pan-cancer evaluation comprised 11 cohorts spanning nine cancer types, including four early-stage cancer cohorts. Yellow fever vaccination and graft-versus-host disease cohorts were retained as non-cancer immune controls. All projects represented in these evaluation cohorts were excluded from construction of the receptor-level training dataset.

Treatment-associated repertoire variation was evaluated using the OSLO-COMET cohort[17]. After repertoire processing, the dataset contained 92 repertoires from 85 patients: 40 repertoires collected without neoadjuvant therapy (No NAT), 36 collected after a treatment interval shorter than 9.5 weeks, and 16 collected after an interval longer than 9.5 weeks. OmniTCR-CES was calculated for each repertoire without cohort-specific model fitting or recalibration. Because some patients contributed more than one repertoire, the repertoire was treated as the analysis unit. Cohort source, cancer types, control sources, patient numbers, and retained repertoire numbers are reported in **Supplementary Table S5**.

Receptor-level transfer was evaluated using independently annotated TCRs from VDJdb[4]. We selected TRB sequences with reported recognition of cancer-associated antigens, including cancer-testis antigens, differentiation antigens, and overexpressed tumour antigens. Antigen names were standardized, duplicate TRB sequences were removed, and sequences overlapping the receptor-level training data were excluded. The resulting dataset contained 29,829 unique tumour-antigen-associated TRB sequences and was compared with an equally sized healthy-control set. This analysis was conducted at the individual-TRB level and did not use repertoire aggregation.

**Table S5**. Composition of TCR repertoire datasets used in the pan-cancer analyses.

| Dataset | Disease type | Source | Counts |
|---|---|---|---|
| Internal (Fig. 4c) | Pan-cancer | TCRdb | 53 repertoires |
| External (Fig.4c) | NSCLC | PRJNA506151 | 43 repertoires (15 patients) |
| Control | Healthy | TCRdb | 83 repertoires |
| External Dataset (Fig4.d,f; Supplementary Fig. 4d) | ES-BRCA | DeepCAT[16] | 32 repertoires (16 patients) |
| | ES-PAAD | | 8 repertoires (8 patients) |
| | ES-KIRC | | 10 repertoires (10 patients) |
| | ES-OV | | 10 repertoires (10 patients) |
| | BLCA | | 117 repertoires (30 patients) |
| | COAD/READ | | 14 repertoires (3 patients) |
| | GBM | | 31 repertoires (15 patients) |
| | NSCLC | | 20 repertoires (20 patients) |
| | SKCM | | 29 repertoires (23 patients) |
| | OV | | 96 repertoires (5 patients) |
| | PAAD | | 16 repertoires (16 patients) |
| | GVHD | | 15 repertoires (15 patients) |
| | YFV | | 9 repertoires (9 patients) |
| | Control | | 666 repertoires (666 patients) |
| OSLO-COMET | COAD/READ | immuneACCESS | 92 repertoires (85 patients) |

#### S1.4 pMHC-conditioned TRB sequence generation datasets

To expand the dataset, the supervised fine-tuning (SFT) dataset for pMHC-conditioned TRB generation was assembled from two sources: the supported peptide-MHC-TRB records from the curated training partition (**Supplementary Section S1.1**), and a supplementary subset of SFT training data obtained from TCRT5[18]. The TCRT5 resource contains a large collection of HLA-I assignments inferred from lower-resolution annotations. Duplicate prompt-target pairs, defined by identical peptide, HLA-I representation and TRB CDR3 sequence, were removed. Because the number of associated TRB sequences varied markedly among pMHC conditions (**Supplementary Fig. 1b**), each exact peptide-HLA condition was capped at 300 associated TRB records by random subsampling with seed 42. We then applied a peptide-overlap filter within each HLA-I representation following the strategy used for TCRT5[18]. Within each HLA-I representation, peptide conditions were ranked by the number of retained records, and a condition was retained only if its peptide shared no contiguous five-residue subsequence with any previously retained peptide. After filtering, the training set contained 141,493 unique supported peptide-MHC-TRB records, comprising 1,563 peptide epitopes, 111 HLA-I representations, 5,098 peptide-HLA conditions and 90,851 TRB CDR3 sequences. Each record provided a peptide and HLA-I pseudo-sequence as the pMHC prompt and an associated TRB CDR3 sequence as the generation target.

After incorporating the additional TCRT5-derived SFT records, we repeated overlap checks against the combined pretraining and generation SFT data. Internal and external generation tests were screened using peptide–HLA–TRB keys extracted from all training records containing these components. For unseen-pMHC generation, screening used peptide–HLA keys and excluded any target pair present in either training stage. These checks included records containing additional components, irrespective of their formatted order.

Internal generation evaluation used supported peptide-MHC-TRB records held out from both autoregressive pretraining and generation fine-tuning (same as **Supplementary Table S3** (Internal P-M-T Dataset)). Held-out records were grouped by their exact peptide-HLA pair, and the associated TRB CDR3 sequences formed the experimentally observed reference collection for each pMHC. The 20 pMHCs with the largest experimentally observed reference TRB collections were selected as generation targets, comprising 3,222 unique observed TRB sequences in total. Each pMHC was treated as an independent evaluation target, and performance was summarized across pMHCs.

External generation evaluation used retained peptide-MHC-TRB records from TRAIT[10]. Records overlapping OmniTCR pretraining or generation fine-tuning data were removed before evaluation. The remaining records were grouped by exact peptide-HLA pair, with the associated unique TRB CDR3 sequences forming the reference collection for each pMHC. The 20 pMHCs with the largest retained reference collections were selected as external targets, together comprising 1,325 target-specific reference TRB CDR3 sequences.

We expanded the unseen-pMHC generation test set with additional records from IMMREP25[19]. Only exact peptide–HLA pairs absent from both autoregressive pretraining and generation supervised fine-tuning (SFT) were retained. The 20 pMHCs with the largest reference collections were selected, comprising 1,036 target-specific reference TRB CDR3 sequences. Of these reference sequences, 986 (95.2%) were associated with peptides absent from both pretraining and SFT. The remaining 50 (4.8%) were associated with peptides present during pretraining but absent from SFT.

Structural evaluation used experimentally determined TCR-pMHC complexes from the TCR3d[20] database. Complexes were eligible only if their first PDB release occurred after 8 April 2025 and their peptide was absent from all OmniTCR training stages. After deduplication, seven complexes were retained as structural evaluation targets, PDB ID: 9GV6, 9GV7, 9IKY, 9HLJ, 9D95, 9K2R, and 9K2I.

### S2. Model implementation and training

#### S2.1 OmniTCR architecture and pretraining

OmniTCR was implemented as a decoder-only Transformer with 12 layers, a hidden dimension of 768, 12 self-attention heads and an intermediate feed-forward dimension of 3,072. The model used rotary positional embeddings, RMS normalization and SwiGLU feed-forward activations, yielding 113,317,632 trainable parameters. The output language-model head projected the final hidden states to the 34-token amino-acid vocabulary.

All model parameters were optimized during autoregressive pretraining using AdamW with $\beta_1 = 0.9$, $\beta_2 = 0.95$, $\epsilon = 1 \times 10^{-8}$ and a weight decay of 0.1. The peak learning rate was $3 \times 10^{-4}$, followed by cosine decay after the warm-up period. Training used bfloat16 mixed precision, gradient checkpointing and DeepSpeed ZeRO stage 2, with the gradient norm clipped at 1.0. Validation loss was evaluated every 32,053 steps (approximately 0.1 epochs), and the checkpoint with the lowest prespecified validation loss was retained for downstream adaptation. **Supplementary Table S6** also reports detailed pretraining settings.

**Table S6**. OmniTCR architecture and pretraining

| Setting | Value |
|---|---|
| Architecture | Decoder-only Transformer |
| Trainable parameters | 113,317,632 |
| Vocabulary size | 34 |
| Decoder layers | 12 |
| Hidden dimension | 768 |
| Attention heads | 12 |
| Dimension per head | 64 |
| Feed-forward dimension | 3,072 |
| Feed-forward activation | SwiGLU |
| Normalization | RMSNorm |
| Positional encoding | RoPE |
| Attention mask | Causal |
| Optimizer | AdamW |
| Learning rate | $3 \times 10^{-4}$ |
| AdamW $\beta_1, \beta_2$ | 0.9, 0.95 |
| AdamW $\epsilon$ | $1 \times 10^{-8}$ |
| Weight decay | 0.1 |
| Learning-rate schedule | Warm-up followed by cosine decay |
| Precision | bfloat16 |
| Gradient clipping | 1.0 |
| Distributed optimization | DeepSpeed ZeRO stage 2 |
| Gradient checkpointing | Enabled |
| Training epochs | 10 |
| Random seed | 42 |

#### S2.2 Fine-tuning for recognition prediction and receptor-level cancer-associated TCR classification

To adapt the pretrained OmniTCR foundation model to recognition prediction and the receptor-level classifier underlying cancer-associated repertoire classification, we performed supervised fine-tuning using task-specific multilayer perceptron (MLP) classification heads.

Both tasks used the sequence-level hidden representation $\mathbf{h}_i \in \mathbb{R}^{d_{\text{model}}}$ (where $d_{\text{model}} = 768$) extracted from the closing sequence-type token of the final decoder layer. For recognition prediction, the extracted representation $\mathbf{h}_i$ was projected through a two-layer MLP classification head incorporating GELU activation and a dropout probability of $0.1$. The intermediate dimension of this MLP was set to $d_{\text{model}}/2 = 384$, and the final projection layer produced a single logit representing the recognition state. For the receptor-level classifier of cancer-associated repertoire, the sequence representation $\mathbf{h}_i$ was passed through a two-layer MLP classification head utilizing a ReLU activation. The first layer projected the hidden state to an intermediate dimension of $d_{\text{model}}/2 = 384$, and the second layer projected the activated state to two logits representing candidate cancer-associated and healthy-background CDR3β classes.

All fine-tuning tasks were optimized using the AdamW optimizer. For recognition prediction, we performed full-parameter fine-tuning, simultaneously updating the parameters of both the OmniTCR backbone and the MLP classification head. For the receptor-level classifier, we compared two fine-tuning configurations: (1) OmniTCR(FT), which froze the pre-trained backbone and only updated the classification head parameters, and (2) OmniTCR(FFT), which performed full-parameter fine-tuning using differential learning rates for the backbone and the classification head. The complete set of fine-tuning hyperparameters for these configurations is summarized in **Supplementary Table S7**.

**Table S7**. Fine-Tuning Hyperparameters for Discriminative Tasks

| **Settings** | **OmniTCR (FFT)** | **OmniTCR (FT)** | **OmniTCR (FFT)** |
|---|---|---|---|
| Tasks | recognition prediction | receptor-level classifier | receptor-level classifier |
| Backbone update | Full | Frozen | Full |
| Backbone learning rate | $1\times 10^{-5}$ | 0 | $1\times 10^{-5}$ |
| Head learning rate | $1\times 10^{-5}$ | $5\times 10^{-6}$ | $1\times 10^{-4}$ |
| Optimizer | AdamW | AdamW | AdamW |
| Weight decay | 0.01 | 0.01 | 0.01 |
| Batch size | 128 | 256 | 256 |
| Training epochs | 3 | 30 | 3 |
| Retained checkpoint | Epoch 3 | Epoch 29 | Epoch 3 |
| Head dimensions | 768→384→1 | 768→384→2 | 768→384→2 |
| Activation | GELU | ReLU | ReLU |
| Dropout | 0.1 | 0 | 0 |
| Loss | Binary cross-entropy | Cross-entropy | Cross-entropy |

#### S2.3 Supervised fine-tuning for pMHC-conditioned TRB generation

OmniTCR(SFT) was initialized from the pretrained OmniTCR backbone and fine-tuned on supported peptide-MHC-TRB records. Each training example consisted of a pMHC prompt followed by its associated TRB CDR3 sequence. The prompt contained the peptide epitope, HLA-I pseudo-sequence and opening [TRB] marker, whereas the generation target contained the TRB CDR3 residues, the closing [TRB] marker and the final [EOS] token. The model predicted each target token from the pMHC prompt and all preceding target tokens. For training example $i$, the conditional log-probability of token $x_{i,t}$ was defined as:

$$\ell_{i,t} = \log p_{\theta}\left(x_{i,t} \mid \mathbf{x}_{i,<t}\right) \tag{S1}$$

The loss was calculated only for the TRB CDR3 residues, the closing [TRB] marker and the [EOS] token. The peptide, HLA-I and opening [TRB] positions were excluded from the loss and served only as conditioning context. All OmniTCR parameters were updated during supervised fine-tuning.

Standard supervised fine-tuning assigns equal weight to every target position. However, the terminal regions of CDR3β contain relatively conserved sequence patterns[21], whereas the central junction is more variable and frequently contributes to peptide-facing contacts[22]. We combined parabolic position weighting with an overlapping-window term that further reweighted autoregressive residue losses. This objective increased supervision of the central CDR3β region and its local sequence windows. For a CDR3β sequence of length $L_\mathrm{i}$, let $j \in \{1, \dots, L_\mathrm{i}\}$ denote the relative residue position. The position weight was defined as:

$$w_{i,j} = \max\left\{1,\ 1 + \beta\left[1 - \left(\frac{(j-1)-(L_i-1)/2}{\max((L_i-1)/2, \epsilon)}\right)^2\right]\right\} \tag{S2}$$

Here, $\varepsilon = 10^{-8}$ prevents division by zero and $\beta = 3$ controls the additional weight assigned near the sequence center. The weight equals 1 at the terminal residues and increases towards an upper bound of 4 near the midpoint. The closing [TRB] and [EOS] tokens retained unit weight. Let $\mathcal{M}_i$ denote all active target positions, including the CDR3β residues and the two termination tokens. The position-weighted token-loss numerator and its normalizing weight were defined as:

$$A_i = \sum_{t \in \mathcal{M}_i} w_{i,t}\,(-\ell_{i,t}), \qquad B_i = \sum_{t \in \mathcal{M}_i} w_{i,t} \tag{S3}$$

For CDR3β residues, $w_{i,t}$ was determined by Equation S2. The closing [TRB] and [EOS] tokens were assigned a weight of 1.

To increase the contribution of contiguous local sequence patterns, we additionally aggregated the autoregressive likelihood over every overlapping n-residue window within the CDR3β sequence. Let $\mathcal{G}_i$ denote the set of all overlapping windows of length n. For each window $g \in \mathcal{G}_i$, its mean position weight was calculated as:

$$\overline{w}_{i,g} = \frac{1}{n}\sum_{j\in g} w_{i,j} \tag{S4}$$

The weighted window-loss numerator and its normalizing term were then defined as:

$$C_i = \sum_{g\in\mathcal{G}_i} \overline{w}_{i,g}\left(-\frac{1}{n}\sum_{j\in g} \ell_{i,j}\right), \qquad D_i = \sum_{g\in\mathcal{G}_i} \overline{w}_{i,g} \tag{S5}$$

Only CDR3β residues were included in the contiguous-window term. The closing [TRB] and [EOS] tokens were excluded.

The position-weighted token term and contiguous-window term were combined and normalized across N training examples:

$$\mathcal{L}_{\mathrm{SFT}} = \frac{\sum_{i=1}^{N}(A_i+\alpha C_i)}{\sum_{i=1}^{N}(B_i+\alpha D_i)} \tag{S6}$$

We used $n = 5$ for the window length and $\alpha = 3.5$ for the relative contribution of the contiguous-window term. The final objective therefore combines residue-level autoregressive supervision with increased weighting of the central CDR3β region and its overlapping local subsequences.

OmniTCR(SFT) was trained using the AdamW optimizer with a maximum learning rate of $2 \times 10^{-4}$ and a weight decay of 0.05. The learning rate was scaled using a cosine scheduler with a warmup ratio of 0.04. To manage the memory constraints of deep autoregressive modeling and maximize training throughput, we employed FlashAttention-2[23] and the DeepSpeed ZeRO stage 2 distributed optimization framework[24, 25]. Gradient clipping was applied with a threshold norm of 0.5. The complete set of training hyperparameters and computational configurations is summarized in **Supplementary Table S8**.

**Table S8**. Supervised Fine-Tuning Hyperparameters for OmniTCR(SFT)

| Setting | Value |
|---|---|
| Initialization | Pretrained OmniTCR |
| Parameter update | Full |
| Optimizer | AdamW |
| Learning rate | $2 \times 10^{-4}$ |
| AdamW $\beta_1, \beta_2$ | 0.9, 0.95 |
| Weight decay | 0.05 |
| Learning-rate schedule | Cosine |
| Warm-up fraction | 0.04 |
| Batch size | 128 |
| Training epochs | 1 |
| Gradient clipping | 0.5 |
| Precision | bfloat16 |
| Attention implementation | FlashAttention-2 |
| Distributed optimization | DeepSpeed ZeRO stage 2 |
| $n$-gram length $n$ | 5 |
| $n$-gram coefficient $\alpha$ | 3.5 |
| Position-weight coefficient $\beta$ | 3.0 |

#### S2.4 Candidate generation and PMI-inspired reranking

Generation was performed using the pMHC prompt and autoregressive continuation format defined in the main Methods. For each target pMHC, beam search was run with a beam width of 400, a maximum generation length of 40 tokens and early stopping. Up to 200 unique CDR3β candidates were retained after exact-sequence deduplication. The opening prompt and generated component-boundary tokens were not included in the reported CDR3β sequence.

OmniTCR(SFT) and OmniTCR(PMI) were evaluated using the same candidate pool. OmniTCR(SFT) ranked candidates by their mean continuation log-likelihood under the target pMHC prompt. OmniTCR(PMI) reranked these candidates using the PMI-inspired score defined in the main Methods. PMI scoring was restricted to the CDR3β residues and the closing [TRB] token and used a null-prior coefficient of 0.8. The 100 highest-ranked unique candidates per pMHC were retained for downstream evaluation. Equal PMI scores were resolved first by the target-prompt continuation likelihood and then by the original beam rank.

### S3. Evaluation settings and comparator implementations

This section expands the evaluation procedures summarized in the main Methods.

#### S3.1 General evaluation protocol and comparator implementation

All evaluations used the records or repertoires retained after quality control, deduplication, and the overlap-removal procedures described in Section S1. Within each evaluation setting, OmniTCR and all applicable comparators were evaluated on the same retained examples. Official code, released checkpoints, and documented inference settings of comparator methods were used directly when available.

For pretrained foundation models without a task-specific predictor, or for comparator methods that required additional training, we trained the corresponding task-specific model using the same training records and labels used for OmniTCR in that task. Official training configurations were used when available; otherwise, hyperparameters were selected using the common validation partition.

Comparator models were selected separately for each downstream evaluation task. For recognition prediction, NetMHCpan-4.2[26] and UnifyImmun[11] were used for P-M; UnifyImmun, TEIM[27] and PanPep[12] were used for P-T; pMTnet[9] was used for P-M-T; EPACT[28] was used for paired-chain P-M-T evaluation including TRA. For cancer-associated repertoire classification, comparators included ESM-2[29], TCR-BERT[30],

BertTCR[31], iCanTCR[32] and DeepCAT[16]. ESM-2 and TCR-BERT were equipped with the same classification head as OmniTCR(FFT) and fully fine-tuned end-to-end using the same training data. For each model, the checkpoint achieving the best validation-set performance was retained for comparison. TCRT5[18], TCRdesign[33] and GRATCR[34] were used as comparators for pMHC-conditioned TRB sequence generation.

#### S3.2 Evaluation of recognition prediction

**Pretrained-model scoring**. OmniTCR(Base) was evaluated without an added classification head. Each record was formatted in the canonical peptide-first order as $x = (x_1, \ldots, x_L)$. Perplexity was calculated over the target positions retained by the causal language-modeling loss:

$$\mathrm{PPL}(x) = \exp\left[-\frac{1}{|\mathcal{M}(x)|}\sum_{t\in\mathcal{M}(x)} \log p_\theta\,(x_t \mid x_{<t})\right] \tag{S7}$$

Amino-acid residues, component-boundary markers, and the final [EOS] token contributed to the score, whereas [BOS] and padding positions did not. Lower perplexity indicated greater language-model support for the complete formatted record. For analyses requiring a score that increased with model support, negative perplexity was used. Perplexity distributions were summarized by their medians, and separation between supported and negative records was quantified using the rank-biserial correlation.

**Receptor clustering.** OmniTCR(Base) representations were extracted from the closing [TRB] token for 28,820 unique TRB CDR3β sequences. The representations were L2-normalized and partitioned into 50 clusters using K-means with the squared-Euclidean objective, k-means++ initialization, 20 initializations and a random seed of 42. Peptide identities and binding labels were not used during clustering. The 34,261 supported peptide-TRB associations were then mapped to these clusters. Association between peptide identity and cluster membership was quantified using bias-corrected Cramer's V. For each peptide, the proportions of its associated TRB sequences assigned to the 50 clusters defined a receptor-cluster profile. Similarity between two receptor-cluster profiles was calculated as one minus their Jensen-Shannon distance, and its association with peptide sequence similarity was summarized using Spearman's rank correlation.

For visualization, clusters were ranked separately for each of the ten selected peptides by the observed-to-expected association ratio, $R_{ec} = O_{ec}/(n_e n_c/N)$, where $O_{ec}$ is the number of records linking peptide $e$ to cluster $c$, $n_e$ and $n_c$ are the corresponding marginal record counts, and $N$ is the total number of merged positive input records. Only clusters with observed associations for the relevant peptide were ranked. Selection proceeded through the first-ranked candidates for all peptides, followed by second-ranked and subsequent candidates, using the peptide order shown in **Fig. 2b** and skipping previously selected clusters until ten distinct clusters had been obtained.

**Performance.** OmniTCR(FFT) assigned each record a probability of belonging to the supported class. Overall performance was measured using the area under the receiver-operating-characteristic curve (AUROC) and the area under the precision-recall curve (AUPRC). Performance among the highest-ranked predictions was evaluated using positive predictive value at rank k (PPV@k), calculated as the number of supported records among the k highest-scoring predictions divided by k. Values of k were 100, 200, 500, 1,000, 2,000, and 5,000 when permitted by the test-set size.

#### S3.3 Evaluation of cancer-associated receptor and repertoire predictions

**OmniTCR-CES.** For each cancer and healthy evaluation repertoire, TRB clonotypes were first clustered with GLIPH2 and filtered using the same clustering and retention criteria applied during construction of the receptor-level classifier dataset. To ensure adequate data quality and sufficient repertoire depth for downstream analysis, only repertoires containing more than 50 GLIPH2-retained TRB clonotypes were included. Each retained clonotype was then assigned a candidate cancer-associated probability by OmniTCR. Retained clonotypes were ranked within each repertoire by clonotype frequency, and the 1,000 most abundant clonotypes were selected; all retained clonotypes were used when fewer than 1,000 remained. OmniTCR-CES was defined as the median predicted probability across the selected clonotypes. The GLIPH2 filtering, frequency-based selection and aggregation procedures were fixed before evaluation and applied identically to all repertoires, without cohort-specific fitting or recalibration.

**Repertoire classification performance.** OmniTCR-CES or the corresponding comparator score was used as a continuous prediction score. AUROC and AUPRC were calculated for the held-out internal repertoire and the external multiregional NSCLC repertoire. For the DeepCAT-derived evaluation, AUROC and AUPRC were calculated separately for each cancer cohort against its corresponding control collection.

**Receptor-level transfer performance.** Transfer to independently annotated tumour-antigen-associated TRB sequences was evaluated using the VDJdb dataset described in **Supplementary Section S1.3**. Direct receptor-level performance was assessed using the cancer-associated probabilities produced by OmniTCR(FFT). In a complementary representation-level analysis, a linear discriminant model was fitted to fixed OmniTCR(FFT) representations of candidate cancer-associated and healthy-background TRBs from the receptor-level training data. The resulting discriminant axis was then applied without refitting to the VDJdb tumour-antigen-associated and healthy-control TRB sequences. Each TRB was projected onto this fixed axis to obtain a discriminant score.

#### S3.4 Evaluation of pMHC-conditioned TRB sequence generation

**General evaluation protocol**. OmniTCR(Base), OmniTCR(SFT), OmniTCR(PMI), and the comparator generators were evaluated on the internal, TRAIT, and unseen-pMHC target sets described in **Supplementary Section S1.4**. Each evaluation used the 20 pMHCs with the largest experimentally observed reference TRB collections, and each method returned 100 valid, unique TRB CDR3 sequences per target. OmniTCR(SFT) and OmniTCR(PMI) used the fixed-pool ranking procedure described before, whereas comparator methods used their released decoding configurations under the same final output budget.

**Sequence validity.** A generated TRB CDR3 sequence was valid when it contained 7-24 amino acids, began with cysteine, ended with phenylalanine or tryptophan, and contained no stop codon or unsupported residue. Generated sequences were compared with the corresponding experimentally observed reference collections in terms of CDR3β length distributions, OLGA[35] human TRB generation probability, global amino-acid frequencies, and position-specific sequence logos. OLGA values were reported as log10 generation probability. Global amino-acid agreement was calculated as Pearson's correlation across the 20 standard amino-acid frequencies. Sequence logos were constructed separately for the dominant 13- and 15-residue classes using all valid sequences of the corresponding length.

**Dependence on pMHC conditioning.** We first assessed whether each generated CDR3β sequence received greater model support from its conditioning pMHC than from alternative pMHCs (**Fig. 5e**). For each of the 20 target pMHCs, the 100 unique CDR3β sequences generated under the complete pMHC prompt were held fixed. For each candidate, the mean per-residue log-likelihood was calculated under its target pMHC and under each of the other 19 complete pMHC prompts. The target-versus-alternative margin was defined as the log-likelihood under the target prompt minus the mean log-likelihood across the 19 alternative prompts. Candidate-level margins are shown in **Fig. 5e** and were averaged within each target pMHC for statistical inference.

We next separated the contributions of peptide and HLA-I context using complementary likelihood- and generation-based perturbation analyses (**Fig. 5f**). For the likelihood analysis, the same top 100 unique CDR3β sequences generated under each complete pMHC prompt were held fixed. For each target, the mean per-residue log-likelihood of this fixed candidate set was calculated under four prompt conditions: the complete pMHC prompt, a peptide-swapped prompt retaining the original HLA-I pseudo-sequence, an HLA-I-swapped prompt retaining the original peptide and a null prompt omitting both conditioning sequences. For each condition, the prompt-support margin was calculated by subtracting a common noncognate baseline, defined as the mean log-likelihood of the same fixed candidate set under the other 19 complete pMHC prompts. The noncognate baseline was therefore identical across the four conditions for each target.

Generated-set retention was evaluated separately by comparing the Top-100 unique sequences generated under each condition, $Y_q$, with those generated under the corresponding full pMHC prompt, $Y_{full}$:

$$R(q) = 100 \times \frac{|Y_q \cap Y_{\text{full}}|}{|Y_{\text{full}}|} \quad \text{(S8)}$$

Each target pMHC contributed one paired observation per condition. The displayed summary values are means with 95% confidence intervals estimated from 10,000 pMHC-level bootstrap resamples. Directional comparisons were performed using raw, one-sided paired Wilcoxon signed-rank tests across the 20 target pMHCs, without multiple-testing correction.

**Cross-target sharing of generated sequences.** For each method $m \in \{\text{SFT}, \text{PMI}\}$,

$Y_i^{(m)}$ denoted the retained Top-100 unique CDR3β sequences for target pMHC $i$. Cross-target sharing was quantified separately for SFT and PMI using the pairwise Jaccard similarity

$$J_{ij}^{(m)} = \frac{\left|Y_i^{(m)} \cap Y_j^{(m)}\right|}{\left|Y_i^{(m)} \cup Y_j^{(m)}\right|} \tag{S9}$$

This produced a symmetric $20 \times 20$ (20 target pMHCs) matrix for each method, with diagonal elements equal to 1. The reported mean Jaccard similarity was calculated across the $\binom{20}{2} = 190$ unique unordered target pairs. For visualization, the SFT values were displayed in the upper triangle and the PMI values in the lower triangle using a common scale from 0 to 1. Target ordering derived from previous hierarchical clustering of the SFT matrix was applied identically to both matrices solely to organize the heatmap. This ordering did not affect the Jaccard calculations, and no clustering topology was displayed or interpreted.

**Reference correspondence and output diversity.** For each of the 20 target pMHCs, 100 generated CDR3β sequences were evaluated against the corresponding deduplicated experimentally observed reference collection. Char-BLEU, sequence recovery, Precision@100, Recall@100 and F1@100 were calculated separately for each target and macro-averaged across the 20 pMHCs.

Char-BLEU was calculated at the character level using NLTK corpus_bleu. For each generated sequence, up to 20 cognate references with the smallest Levenshtein distances were selected. Character one- to four-grams were assigned uniform weights of 0.25, and Chen-Cherry smoothing method 1 was applied.

$$\mathrm{CharBLEU}_i = \mathrm{BP}_i \exp\left(\frac{1}{4}\sum_{n=1}^{4} \log \tilde{p}_{i,n}\right) \tag{S10}$$

Here, $\tilde{p}_{i,n}$ is the smoothed modified precision for character n-grams, and $\mathrm{BP}_i$ is the corpus-level brevity penalty. Char-BLEU was assigned a value of zero when no valid sequence was retained.

Sequence recovery quantified similarity to the closest experimentally observed sequence. For each generated CDR3β sequence, the experimentally observed reference with the smallest Levenshtein distance was selected from the same-length references when available. Otherwise, the closest reference across all lengths was used. Per-sequence recovery was calculated as:

$$\rho_i(g) = \begin{cases} \frac{1}{|g|}\sum_{t=1}^{|g|} \mathbf{1}\left(g_t = r_{i,t}^*(g)\right), & |r_i^*(g)| = |g| \\ 1 - \frac{d_{\mathrm{Lev}}\left(g, r_i^*(g)\right)}{\max(|g|, |r_i^*(g)|)}, & |r_i^*(g)| \neq |g| \end{cases} \quad \text{(S11)}$$

The target-level sequence-recovery score was the mean across the 100 generated sequences, where $\mathcal{G}_i$ denotes the collection of 100 sequences generated for target $i$:

$$\mathrm{Recovery}_i = \frac{1}{100}\sum_{g\in\mathcal{G}_i} \rho_i(g) \quad \text{(S12)}$$

Exact experimentally observed reference recovery was evaluated using Precision@100, Recall@100 and F1@100. For pMHC target $i$, let $c_i$ denote the number of generated sequences that exactly matched an experimentally observed reference sequence. Let $u_i$ denote the number of unique experimentally observed reference sequences recovered. Precision@100, Recall@100 and F1@100 were calculated as:

$$P_i^{(100)} = \frac{c_i}{100}, \ R_i^{(100)} = \frac{u_i}{\min(100, |\mathcal{R}_i|)} \quad \text{(S13)}$$

$$F_{1,i}^{(100)} = \frac{2P_i^{(100)}R_i^{(100)}}{P_i^{(100)} + R_i^{(100)}} \quad \text{(S14)}$$

where $\mathcal{R}_i$ is the experimentally observed reference collection for target $i$.

**Physicochemical properties.** Four sequence-level physicochemical properties were calculated for each experimentally observed reference sequence and generated CDR3β sequence. For each property, sequence-level values were pooled across the 20 target pMHCs separately for generated and experimentally observed sequences. Differences between the resulting distributions were summarized using the two-sample Kolmogorov-Smirnov statistic and the Wasserstein distance.

For a sequence $x = (a_1, \dots, a_L)$ of length $L$, GRAVY was the mean residue hydropathy on the Kyte-Doolittle scale.

$$\mathrm{GRAVY}(x) = \frac{1}{L}\sum_{t=1}^{L} h_{\mathrm{KD}}(a_t) \quad \text{(S15)}$$

Here, $h_{\mathrm{KD}}(a_t)$ denotes the Kyte-Doolittle hydropathy. GRAVY was calculated using ProteinAnalysis.gravy() from Biopython[36].

Polarity was the mean of the residue-specific Grantham polarity values:

$$\mathrm{Polarity}(x) = \frac{1}{L}\sum_{t=1}^{L} p_{\mathrm{G}}(a_t) \quad \text{(S16)}$$

Here, $p_{\mathrm{G}}(a_t)$ denotes the Grantham polarity values assigned to residue $a_t$.

Aromaticity was defined as the fraction of phenylalanine, tryptophan and tyrosine residues.

$$\mathrm{Aromaticity}(x) = \frac{1}{L}\sum_{t=1}^{L} \mathbf{1}(a_t \in \{\mathrm{F}, \mathrm{W}, \mathrm{Y}\}) \quad \text{(S17)}$$

Aromaticity was calculated using ProteinAnalysis.aromaticity().

The hydrophobic-residue fraction was defined using alanine, valine, isoleucine, leucine, methionine, phenylalanine, tryptophan and tyrosine:

$$\mathrm{HF}(x) = \frac{1}{L}\sum_{t=1}^{L} \mathbf{1}(a_t \in \{\mathrm{A}, \mathrm{V}, \mathrm{I}, \mathrm{L}, \mathrm{M}, \mathrm{F}, \mathrm{W}, \mathrm{Y}\}) \quad \text{(S18)}$$

where $\mathbf{1}(\cdot)$ is the indicator function. The hydrophobic fraction (HF) was calculated directly from the specified residue set.

**AlphaFold 3 structural evaluation.** For each structural evaluation target, 100 generated CDR3β sequences per method were inserted into the experimentally observed TRB framework and combined with the corresponding TRA chain, peptide, MHC-I heavy chain, and β2-microglobulin. Then they were modeled by AlphaFold 3. The experimentally observed TRB was processed through the same AlphaFold 3 workflow. Structural confidence was evaluated using mean TRB pLDDT, TRB pTM, TRB ipTM and TRB interface predicted alignment error (iPAE). Mean TRB pLDDT summarized residue-level confidence of TRB, TRB pTM summarized confidence in the TRB structure, TRB ipTM summarized confidence in the positioning of TRB relative to the remaining complex, and TRB iPAE summarized predicted alignment error across the defined TRB interface. Higher pLDDT, pTM, and ipTM and lower iPAE indicated greater AF3-predicted confidence. A generated complex satisfied the composite AF3 structural-confidence criterion only when it simultaneously exceeded the experimentally observed TRB model in pLDDT, pTM, and ipTM and had lower iPAE. For each target and method, the pass rate was defined as the proportion of the 100 generated candidates satisfying all four conditions.

#### S3.5 Statistical analysis

Exact *P* values were reported where numerical precision permitted, and $P < 0.05$ was considered statistically significant. The analysis unit was defined according to each analysis as an individual recognition record, unique TRB CDR3 sequence, peptide pair, repertoire or target pMHC.

For recognition-prediction benchmarks, differences in AUROC between models evaluated on the same records were assessed using DeLong's method. Differences in AUPRC were assessed by nonparametric bootstrap testing with 10,000 resamples of the evaluation records. Bootstrap-derived 95% confidence intervals for AUROC and AUPRC were defined by the 2.5th and 97.5th percentiles of the corresponding bootstrap distributions. For the paired-chain analysis in **Fig. 3c**, AUROC and AUPRC were recalculated from 10,000 bootstrap resamples for each input condition. Empirical *P* values were derived from the bootstrap distributions of the metric differences between inputs with and without the paired TCRα chain across 2,535 matched TRAIT records.

For **Fig. 2a**, perplexity distributions for supported and negative records were compared using Mann-Whitney U tests separately for the P-M, P-T and P-M-T tasks. The three resulting *P* values were adjusted using the Benjamini-Hochberg procedure, and distributional separation was summarized using rank-biserial correlations. The association between peptide identity and TCR-cluster membership was quantified using bias-corrected Cramér's V. Significance was assessed using 10,000 permutations of cluster assignments among unique TCRs while retaining their observed peptide associations (random seed, 42). Peptide-specific cluster enrichment was assessed using hypergeometric tests, followed by Benjamini-Hochberg correction across all tested peptide-cluster combinations. In **Supplementary Fig. 2b**, the difference in mean Jensen-Shannon profile similarity between within-SARS and SARS-reference peptide pairs was assessed using 10,000 permutations of peptide-category labels. In **Supplementary Fig. 2d**, the association between peptide-sequence similarity and TCR-cluster profile similarity was evaluated using Spearman's rank correlation across all 45 unique peptide pairs. Comparisons among clonotype-expansion groups and pMHC-barcode UMI tertiles in **Fig. 3e** used Mann-Whitney U tests, with Benjamini-Hochberg adjustment within each set of comparisons.

For the VDJdb receptor-level transfer analysis in **Fig. 4e**, fixed linear-discriminant scores for the 29,829 tumour-antigen-associated TRB sequences and 29,829 healthy-control TRB sequences were compared using Welch's *t*-test. Repertoire-level comparisons in **Fig. 4g** and **Supplementary Fig. 4d** used Mann-Whitney U tests.

For generation analyses, differences between experimentally observed and generated sequence distributions were summarized using two-sample Kolmogorov-Smirnov statistics and Wasserstein distances for sequence length, OLGA log10 generation probability and physicochemical properties. Global amino-acid usage in **Fig. 5c** and **Supplementary Fig. 5c** was assessed using Pearson's correlation across the frequencies of the 20 standard amino acids. For **Fig. 5e**, target-level mean likelihood margins were compared with zero using a two-sided Wilcoxon signed-rank test. For **Supplementary Fig. 5e**, margins were averaged across the 100 retained PMI-ranked candidates for each of the 20 pMHC targets. These target-level means were tested against zero using a one-sided Wilcoxon signed-rank test with the alternative hypothesis of positive margins. Prespecified directional comparisons in **Fig. 5f** and **Supplementary Fig. 5f, g** used one-sided paired Wilcoxon signed-rank tests across 20 pMHCs without multiple-testing correction. **Supplementary Fig. 5f, g** were evaluated separately at top-*k* values of 10, 20, 50 and 100. Confidence intervals for pMHC-level summaries were estimated using 10,000 target-level bootstrap resamples. Char-BLEU, sequence-recovery and F1@100 values were summarized as mean ± s.e.m. across the 20 target pMHCs. Unique-sequence counts and AF3 structural-confidence measurements were treated as descriptive summaries.

Benjamini-Hochberg correction was applied only to the comparisons specified above. Other *P* values were unadjusted unless stated otherwise. Random seed 42 was used for TCR clustering and the associated permutation analyses.

### S4. Additional analyses of recognition

#### S4.1 Peptide-associated organization of TCR-cluster profiles

We further examined the receptor-cluster profiles underlying the peptide-associated organization reported in main **Fig. 2b**. As described in Section S3.2, each of the ten peptides was represented by the distribution of its associated TRB sequences across the

50 receptor clusters. Principal coordinates analysis based on Jensen-Shannon distance provided an overall view of the relationships among these peptide-specific profiles, with SARS-CoV-1 and SARS-CoV-2 peptides occupying relatively similar regions of the profile space compared with the reference viral peptides (**Supplementary Fig. 2a**).

Consistent with the quantitative comparison reported in the main text, receptor-cluster profile similarity was higher for peptide pairs within the same SARS group than for SARS-reference pairs (mean, 0.8876 versus 0.4862; $P = 5.0 \times 10^{-4}$, category-label permutation test with 10,000 permutations; **Supplementary Fig. 2b**). The complete pairwise similarity matrix resolved these relationships across all ten peptides and showed the corresponding high-similarity structure among SARS-associated peptides (**Supplementary Fig. 2c**). Across all 45 unique peptide pairs, receptor-cluster profile similarity was positively associated with peptide-sequence similarity (Spearman's $\rho = 0.4233$, $P = 0.0038$; **Supplementary Fig. 2d**).

#### S4.2 Embedding visualization across external and unseen-epitope recognition datasets of OmniTCR(FFT)

To further visualize the representation space from OmniTCR(FFT) underlying the external and unseen-epitope evaluations, we extracted the final-layer hidden state of the penultimate token for each record and projected the resulting OmniTCR(FFT) embeddings into two dimensions using UMAP. The penultimate token was the component-closing token immediately preceding [EOS], corresponding to the closing [TRB] token for TCR-containing inputs. Points were coloured by supported or negative association status.

NetMHCpan P-M records and TRAIT P-T and P-M-T records showed task-dependent, partial separation between supported and negative associations (**Supplementary Fig.**

**2e**). Similar organization was observed in the independent SARS-CoV-2 P-T and P-M-T datasets (**Supplementary Fig. 2f**). In the unseen-epitope evaluations, P-M records remained broadly separated, whereas P-T and P-M-T records were more intermixed but retained structured distributions (**Supplementary Fig. 2g**). These projections complement the AUROC, AUPRC and TCRα-chain analyses in main **Fig. 3**.

#### S4.3 Layer-wise redistribution of attribution in the NR1C recognition context

We examined how token-level attribution patterns changed across Transformer layers for the NR1C recognition context analyzed in main **Fig. 3**. Layer-wise integrated-gradient attribution was calculated for the YLQPRTFLL-HLA-A*02:01 input and for YLQPRTFLL paired with the NR1C CDR3β sequence CAGQVTNTGELFF (**Supplementary Fig. 3a, b**).

Layer-wise integrated gradients were computed for the pre-sigmoid recognition logit using Captum's LayerIntegratedGradients. Attribution was calculated separately for the output hidden states of each decoder block in the task-specific recognition classifier. The baseline replaced every token with the padding-token embedding while retaining the original attention mask. This replacement included both sequence-boundary and component-boundary tokens, and the classifier readout position remained fixed. Attributions were computed using 300 integration steps and summarized at each token position by the L2 norm across hidden dimensions. For visualization, each layer's token-level magnitudes were divided by its maximum magnitude with a stabilizing constant of $10^{-9}$. The heatmaps therefore display relative attribution magnitudes within each layer.

In early and intermediate layers, relatively high attribution was distributed across multiple peptide, HLA-I and CDR3β positions together with component-boundary tokens. In later layers, the relative attribution pattern shifted towards component-closing and [EOS] tokens while selected residue positions remained prominent. Similar redistribution was observed for both the peptide-HLA-I and peptide-TRB inputs. These patterns are consistent with progressive consolidation of residue-level information into the component-level representations used for recognition prediction.

### S5. Additional analyses of cancer-associated repertoires

#### S5.1 Differences between the candidate cancer-associated and the healthy-background TRBs

We characterized sequence-level differences between the 1,489,111 candidate cancer-associated TRB CDR3 sequences used for classifier training and the equally sized, clonotype-frequency-matched healthy-background training set. We compared CDR3β length, global amino-acid usage and sequence-level physicochemical descriptors. Distributional differences in the physicochemical descriptors were summarized using rank-biserial correlations.

The two receptor sets had strongly overlapping length distributions, with both concentrated between 14 and 16 residues (**Supplementary Fig. 4a**). Differences in global amino-acid usage were distributed across several residues: valine, leucine, arginine and isoleucine were relatively more frequent among the candidate cancer-associated set, whereas serine, alanine and phenylalanine were more frequent in the healthy-background set (**Supplementary Fig. 4b**). Differences in physicochemical descriptors were also small in magnitude. The largest shifts towards the candidate cancer-associated set involved sequence entropy, basic-residue fraction and net-charge proxy, with effect sizes of 0.082, 0.081 and 0.066, respectively. Aromatic- and polar-residue fractions showed the largest shifts towards the healthy-background set, with effect sizes of −0.129 and −0.080 (**Supplementary Fig. 4c**). These results indicate that the two receptor sets differed through multiple small sequence-level shifts rather than a single dominant length, residue or bulk physicochemical property. This supports the use of a learned sequence representation to capture the more distributed sequence differences between the two sets.

##### S5.2 Distribution of OmniTCR-CES across control, early-stage and established cancer repertoires

We further computed the distribution of OmniTCR-CES after grouping repertoires into control, early-stage cancer and established cancer categories. The three groups contained 690, 60 and 323 repertoires, respectively (**Supplementary Fig. 4d**). CES values were higher in early-stage cancer than in control repertoires ($P < 2.2 \times 10^{-16}$) and were further increased in established cancer relative to early-stage cancer ($P =$ 0.0029; Mann-Whitney U tests). These comparisons extend the repertoire-level classification in main **Fig. 4** by showing that receptor-level predictions accumulate into a graded repertoire-level signal that is already detectable in early-stage disease by OmniTCR-CES.

#### S6. Additional analyses of conditional generation and structural evaluation

##### S6.1 Sequence characteristics of PMI-ranked TRB candidates

OmniTCR(PMI) reranked a fixed pool of up to 200 candidates generated by OmniTCR(SFT) for each pMHC and retained the top 100 according to the PMI-inspired score. We compared the resulting 2,000 candidates across 20 pMHCs with their corresponding experimentally observed reference TRB collections using CDR3β length, OLGA generation probability, global amino-acid usage and length-specific sequence logos. The pMHC-conditioned likelihood margins were also evaluated for the PMI-ranked TRB candidates.

PMI-ranked candidates retained the dominant 13-16-residue length range and had higher OLGA generation probabilities than the experimentally observed reference sequences (**Supplementary Fig. 5a, b**). Their global amino-acid frequencies closely matched the experimentally observed reference collection (Pearson's $r = 0.982$, $P = 1.6 \times 10^{-14}$; **Supplementary Fig. 5c**). Sequence logos for the dominant 13- and 15-residue classes showed similar N-terminal CASS-like and C-terminal EQYF- or NEQYF-like organization, together with diversity at central junctional positions (**Supplementary Fig. 5d**). PMI-ranked candidates generally received higher mean per-token log-likelihoods under their designated pMHC prompts than under the other 19 pMHC prompts. Candidate-level margins were averaged within each target, yielding 20 target-level means. These means were shifted above zero (one-sided Wilcoxon signed-rank test, $P = 9.5 \times 10^{-7}$; **Supplementary Fig. 5e**). PMI reranking therefore changed candidate prioritization while retaining the principal sequence properties established by SFT.

#### S6.2 PMI reranking increases target-relative support and reduces cross-pMHC sharing

We compared the original SFT ranking and PMI ranking of the same candidate pools at top-$k$ values of 10, 20, 50 and 100. Across the 20 target pMHCs, PMI-ranked candidates had larger target-versus-mean-noncognate likelihood margins at every cutoff (all $P < 1.0 \times 10^{-4}$; paired Wilcoxon signed-rank tests; **Supplementary Fig. 5f**). Cross-pMHC sharing was also consistently reduced after PMI reranking. Mean pairwise Jaccard similarity among pMHC-conditioned candidate sets was also lower at each cutoff (all $P < 1.0 \times 10^{-4}$; paired Wilcoxon signed-rank tests; **Supplementary Fig. 5g**). At $k =$

100, mean Jaccard similarity decreased from 0.0870 to 0.0323, corresponding to a 62.9% reduction in cross-pMHC sharing.

PMI reranking also changed which candidates entered the retained top-k sets. The proportion of PMI-ranked candidates absent from the corresponding original SFT top-k set decreased as *k* increased but remained substantial when the top 100 candidates were retained (**Supplementary Fig. 5h**). These results show that PMI reranking selected a different subset from the fixed SFT candidate pool, with greater target-relative model support and less sequence sharing across pMHC conditions.

#### S6.3 Generation performance on unseen pMHC targets

Generation was evaluated on 20 unseen pMHC targets from a test set expanded with IMMREP25[19] records (Supplementary Section S1.4). All target peptide–HLA pairs were absent from both pretraining and generation SFT. Of the 1,036 target-specific reference TRB CDR3 sequences, 986 were associated with peptides absent from both training stages. The remaining 50 were associated with peptides encountered during pretraining but absent from SFT. Each method generated 100 valid, unique CDR3β sequences per target. Performance was summarized using Char-BLEU, sequence recovery, exact-reference F1@100 and the total number of unique outputs.

OmniTCR(SFT) exceeded TCRT5 by 0.0148 in Char-BLEU and by 0.0145 in sequence recovery (**Supplementary Fig. 5i**). OmniTCR(PMI) retained similar reference correspondence after reranking, and OmniTCR(Base) remained competitive with the task-specific generators. Exact-reference F1@100 values were low for all methods, indicating that most outputs were not direct copies of observed receptors. TCRdesign produced the largest number of unique sequences but lower Char-BLEU and sequence recovery, showing that output diversity did not directly track experimentally observed reference TRB sequences.

#### S6.4 Ablation of the TCR-specific SFT objective

To assess the contribution of the proposed training objective, we compared the models trained with standard token-level SFT and our TCR-specific objective described in Section S2.3. The TCR-specific objective increased Char-BLEU from 0.8117 to 0.8272 on the internal set, from 0.7266 to 0.7328 on TRAIT and from 0.7947 to 0.7991 on the unseen-pMHC benchmark (**Supplementary Table S9**). Sequence recovery increased from 0.7342 to 0.7456, from 0.6624 to 0.6724 and from 0.6717 to 0.6876 across the same evaluations. The largest recovery gain occurred on the unseen-pMHC benchmark, whereas F1@100 changed little across the three evaluations. The objective therefore improved sequence-level correspondence.

**Table S9**. Ablation of the TCR-specific supervised fine-tuning objective.

| | Char-BLEU | | Sequence recovery | | F1@100 | |
|---|---|---|---|---|---|---|
| | Std. SFT | **TCR-SFT** | Std. SFT | **TCR-SFT** | Std. SFT | **TCR-SFT** |
| Internal | 0.8117 | **0.8272** | 0.7342 | **0.7456** | **0.0147** | 0.0143 |
| TRAIT | 0.7266 | **0.7328** | 0.6624 | **0.6724** | **0.0049** | 0.0048 |
| Unseen-pMHC | 0.7947 | **0.7991** | 0.6717 | **0.6876** | 0 | **0.0005** |

#### S6.5 Best-of-100 AlphaFold 3 structural-confidence analysis

We also examined the highest-ipTM candidate obtained within the fixed generation budget of 100 CDR3β sequences per method for each of the seven post-cutoff, epitope-excluded pMHC targets. Each generated CDR3β sequence was inserted into the matched TCRβ framework and modeled using AlphaFold 3 (AF3). The experimentally observed CDR3β sequence for each target was processed using the same modeling workflow. For each method and target, the generated candidate with the highest TRB ipTM was retained and evaluated using TRB pTM, ipTM, iPAE and mean pLDDT.

Across the seven targets, OmniTCR(PMI) achieved the highest mean TCRβ ipTM among all groups (0.6086), compared with 0.5814 for the experimentally observed sequence models, and exceeded the matched observed sequence in four targets (**Supplementary Fig. 6a**). OmniTCR(PMI) also achieved the lowest mean iPAE among generated methods, at 16.55 Å compared with 17.41 Å for OmniTCR(SFT). The experimentally observed sequence models still had a lower mean iPAE of 14.43 Å. Mean TCRβ pTM was similar between the selected OmniTCR(PMI) candidates and the experimentally observed sequence models (0.8529 versus 0.8643).

The FSGEYIPTV-HLA-A*02:01 complex (PDB ID: 9K2I) provided another representative example. The selected OmniTCR(PMI) candidate, CASSLGDSSYEQYF, had a TRB pTM of 0.8600, while ipTM increased from 0.47 in the experimentally observed sequence model to 0.77, iPAE decreased from 17.23 Å to 9.20 Å and mean pLDDT increased from 89.36 to 90.88. In the AF3-predicted complex, its central LGDSSY segment contacted the exposed YIP peptide core through 19 heavy-atom contacts within 4.0 Å, concentrated around peptide Tyr5 and Pro7 (**Supplementary Fig. 6b**). Tyr101 formed the closest contact with Pro7 at 3.06 Å. No direct CDR3β-peptide hydrogen bonds were detected; instead, the predicted interface was dominated by van der Waals contacts, hydrophobic and aromatic packing and shape complementarity. This example shows that a sequence-divergent generated CDR3β can be accommodated at the peptide-facing interface in an AF3-predicted complex with a contact arrangement distinct from that of the matched experimentally observed sequence model.

### References

1. Yue, T. et al. TCRdb 2.0: an updated T-cell receptor sequence database. *Nucleic Acids Res* **54**, D504–D510 (2026).
2. Corrie, B.D. et al. iReceptor: A platform for querying and analyzing antibody/B-cell and T-cell receptor repertoire data across federated repositories. *Immunol Rev* **284**, 24–41 (2018).
3. Vita, R. et al. The Immune Epitope Database (IEDB): 2018 update. *Nucleic Acids Res* **47**, D339–D343 (2019).
4. Shugay, M. et al. VDJdb: a curated database of T-cell receptor sequences with known antigen specificity. *Nucleic Acids Res* **46**, D419–D427 (2018).
5. Tickotsky, N., Sagiv, T., Prilusky, J., Shifrut, E. & Friedman, N. McPAS-TCR: a manually curated catalogue of pathology-associated T cell receptor sequences. *Bioinformatics* **33**, 2924–2929 (2017).
6. Albert, B.A. et al. Deep neural networks predict class I major histocompatibility complex epitope presentation and transfer learn neoepitope immunogenicity. *Nat Mach Intell* **5**, 861–872 (2023).
7. Reynisson, B., Alvarez, B., Paul, S., Peters, B. & Nielsen, M. NetMHCpan-4.1 and NetMHCIIpan-4.0: improved predictions of MHC antigen presentation by concurrent motif deconvolution and integration of MS MHC eluted ligand data. *Nucleic Acids Res* **48**, W449–W454 (2020).
8. Zhang, W. et al. A framework for highly multiplexed dextramer mapping and prediction of T cell receptor sequences to antigen specificity. *Sci Adv* **7**, eabf5835 (2021).
9. Lu, T.S. et al. Deep learning-based prediction of the T cell receptor-antigen binding specificity. *Nat Mach Intell* **3**, 864–875 (2021).
10. Wei, M.M. et al. TRAIT: A Comprehensive Database for T-cell Receptor-antigen Interactions. *Genom Proteom Bioinf* **23**, qzaf033 (2025).
11. Yu, C.P., Fang, X., Tian, S.Y. & Liu, H. A unified cross-attention model for predicting antigen binding specificity to both HLA and TCR molecules. *Nat Mach Intell* **7**, 278–292 (2025).
12. Gao, Y.C. et al. Pan-Peptide Meta Learning for T-cell receptor-antigen binding recognition. *Nat Mach Intell* **5**, 236–249 (2023).
13. Minervina, A.A. et al. SARS-CoV-2 antigen exposure history shapes phenotypes and specificity of memory CD8+ T cells. *Nat Immunol* **23**, 781–790 (2022).
14. Huang, H., Wang, C.L., Rubelt, F., Scriba, T.J. & Davis, M.M. Analyzing the Mycobacterium tuberculosis immune response by T-cell receptor clustering with GLIPH2 and genome-wide antigen screening. *Nat Biotechnol* **38**, 1194–1202 (2020).
15. Jia, Q.Z. et al. Local mutational diversity drives intratumoral immune

heterogeneity in non-small cell lung cancer. *Nat Commun* **9**, 5361 (2018).

16. Beshnova, D. et al. De novo prediction of cancer-associated T cell receptors for noninvasive cancer detection. *Sci Transl Med* **12**, eaaz3738 (2020).
17. Hoye, E. et al. T cell receptor repertoire sequencing reveals chemotherapy-driven clonal expansion in colorectal liver metastases. *Gigascience* **12**, giad032 (2023).
18. Karthikeyan, D., Bennett, S.N., Reynolds, A.G., Vincent, B.G. & Rubinsteyn, A. Conditional generation of real antigen-specific T cell receptor sequences. *Nat Mach Intell* **7**, 1494–1509 (2025).
19. IMMREP25 Organizers. IMMREP25: TCR Specificity Prediction Challenge. https://kaggle.com/competitions/immrep25, 2025. Kaggle.
20. Lin, V.L.R. et al. TCR3d 2.0: expanding the T cell receptor structure database with new structures, tools and interactions. *Nucleic Acids Res* **53**, D604–D608 (2025).
21. Davis, M.M. & Bjorkman, P.J. T-Cell Antigen Receptor Genes and T-Cell Recognition. *Nature* **334**, 395–402 (1988).
22. Glanville, J. et al. Identifying specificity groups in the T cell receptor repertoire. *Nature* **547**, 94–98 (2017).
23. Dao, T. FlashAttention-2: Faster Attention with Better Parallelism and Work Partitioning. In *The Twelfth International Conference on Learning Representations* (2024).
24. Rajbhandari, S., Rasley, J., Ruwase, O. & He, Y.X. ZeRO: Memory Optimizations Toward Training Trillion Parameter Models. *Proceedings of Sc20: The International Conference for High Performance Computing, Networking, Storage and Analysis (Sc20)* (2020).
25. Rasley, J., Rajbhandari, S., Ruwase, O. & He, Y.X. DeepSpeed: System Optimizations Enable Training Deep Learning Models with Over 100 Billion Parameters. *Kdd '20: Proceedings of the 26th Acm Sigkdd International Conference on Knowledge Discovery & Data Mining*, 3505–3506 (2020).
26. Nilsson, J.B., Greenbaum, J., Peters, B. & Nielsen, M. NetMHCpan-4.2: improved prediction of CD8+epitopes by use of transfer learning and structural features. *Front Immunol* **16**, 1616113 (2025).
27. Peng, X.A. et al. Characterizing the interaction conformation between T-cell receptors and epitopes with deep learning. *Nat Mach Intell* **5**, 395–407 (2023).
28. Zhang, Y.M. et al. Epitope-anchored contrastive transfer learning for paired CD8+ T cell receptor-antigen recognition. *Nat Mach Intell* **6**, 1344–1358 (2024).
29. Lin, Z.M. et al. Evolutionary-scale prediction of atomic-level protein structure with a language model. *Science* **379**, 1123–1130 (2023).
30. Wu, K. et al. TCR-BERT: learning the grammar of T-cell receptors for flexible antigen-binding analyses. *Pr Mach Learn Res* **240**, pp. 194–229 (2024).
31. Zhang, M. et al. BertTCR: a Bert-based deep learning framework for predicting cancer-related immune status based on T cell receptor repertoire. *Brief*

*Bioinform* **25**, bbae420 (2024).

32. Cai, Y.D. et al. The Deep Learning Framework iCanTCR Enables Early Cancer Detection Using the T-cell Receptor Repertoire in Peripheral Blood. *Cancer Res* **84**, 1915–1928 (2024).
33. Li, X.K. et al. TCRdesign: an antigen-specific generative language model for de novo design of T-cell receptors. *Brief Bioinform* **26**, bbaf691 (2025).
34. Zhou, Z.H. et al. GRATCR: Epitope-Specific T Cell Receptor Sequence Generation With Data-Efficient Pre-Trained Models. *Ieee J Biomed Health* **29**, 2271–2283 (2025).
35. Sethna, Z., Elhanati, Y., Callan, C.G., Jr., Walczak, A.M. & Mora, T. OLGA: fast computation of generation probabilities of B- and T-cell receptor amino acid sequences and motifs. *Bioinformatics* **35**, 2974–2981 (2019).
36. Cock, P.J.A. et al. Biopython: freely available Python tools for computational molecular biology and bioinformatics. *Bioinformatics* **25**, 1422–1423 (2009).

### Supplementary figure legends

**Supplementary Fig. 1 | Coverage and sequence composition of the OmniTCR pretraining corpus. a,b**, Distributions of the number of unique TCRβ (TRB) CDR3 sequences associated with individual peptides (**a**; *n* = 2,271; median, 4; 90th percentile, 390) and peptide-HLA-I combinations (pMHCs) (**b**; *n* = 1,797; median, 2; 90th percentile, 17). **c**, Number of unique pMHCs associated with each of 70,192 TRB CDR3 sequences. The upper bar summarizes associations with one, two or at least three pMHCs; the lower bars resolve associations with at least three pMHCs. **d**, Numbers of unique peptides and TRB CDR3 sequences associated with the 20 HLA-I alleles having the greatest peptide coverage. **e**, Length distributions of unique peptides (*n* = 434,143), TCRα (TRA) CDR3 sequences (*n* = 5,095,003) and TRB CDR3 sequences (*n* = 319,410,970). **f**, Number of unique HLA-I alleles associated with each peptide (*n* = 415,686 peptides with at least one HLA-I annotation). **g, h**, Sequence logos for the two most abundant TRA CDR3 length classes (13 and 14 residues; **g**) and TRB CDR3 length classes (15 and 16 residues; **h**). Sequence counts are shown above each logo, and letter height denotes information content in bits.

**Supplementary Fig. 2 | Peptide-associated receptor-cluster profiles and embedding structure across recognition datasets. a**, Principal coordinates analysis (PCoA) of peptide-specific receptor-cluster profiles. Each profile represents the normalized distribution of associated TRB sequences across the 50 clusters, and pairwise distances are base-2 Jensen-Shannon distances. Colours and symbols denote SARS-CoV-1, SARS-CoV-2 and reference viral peptides. Peptide identity was associated with cluster membership (bias-corrected Cramér's V = 0.2251; $P = 9.99 \times 10^{-5}$, unique-TCR-level permutation test with 10,000 permutations). **b**, Receptor-cluster profile similarity for peptide pairs within the same SARS group and for SARS-reference pairs. Similarity was defined as one minus the Jensen-Shannon distance. Points denote peptide pairs.

$P = 5.0 \times 10^{-4}$, category-label permutation test with 10,000 permutations. **c**, Pairwise receptor-cluster profile similarity matrix for the ten peptides. **d**, Association between length-normalized Levenshtein peptide-sequence similarity and receptor-cluster profile similarity across all 45 unique peptide pairs. The line is an ordinary least-squares fit used as a visual guide, with shading showing its 95% confidence interval. Spearman's $\rho = 0.4233$; $P = 0.0038$. **e**, UMAP projections of NetMHCpan-4.1 P-M (n = 6,042), TRAIT P-T (n = 6,842) and TRAIT P-M-T (n = 5,945) records. **f**, Corresponding projections for SARS-CoV-2 P-T (n = 6,470) and P-M-T (n = 6,488) records. **g**, Projections for unseen-epitope P-M (n = 34,750), P-T (n = 642) and P-M-T (n = 141) records. Embeddings in **e-g** were extracted from the component-closing token immediately preceding [EOS]. Gold and purple denote negative and supported records, respectively. P-M, peptide-MHC; P-T, peptide-TCRβ; P-M-T, peptide-MHC-TCRβ; UMAP, uniform manifold approximation and projection.

**Supplementary Fig. 3 | Layer-wise attribution patterns in the NR1C recognition complex.** a, YLQPRTFLL–HLA-A*02:01 input. b, YLQPRTFLL paired with the NR1C CDR3β sequence CAGQVTNTGELFF. Layer-wise integrated gradients attributed the pre-sigmoid recognition logit to the output hidden states of each decoder block in the corresponding task-specific classifier. At each token position, attribution magnitude was calculated as the L2 norm across hidden dimensions and scaled by the maximum within its layer. Rows denote layers L0–L11, and columns denote token positions in model input order. Colour intensity indicates relative attribution magnitude within each layer. [BOS] and [EOS] denote sequence-boundary tokens; [EPI], [HLA] and [TRB] denote component-boundary tokens.

**Supplementary Fig. 4 | Sequence characteristics of candidate cancer-associated TRBs and repertoire-level CES distributions. a**, Kernel-density estimates of CDR3β length for 1,489,111 candidate cancer-associated TCRs and an equally sized, clonotype-frequency-matched healthy-background set. **b**, Differences in global amino-acid frequency between the candidate cancer-associated and healthy-background sets. Positive values indicate higher frequency in the candidate cancer-associated set, and negative values indicate higher frequency in the healthy-background set. **c**, Rank-biserial correlations for sequence-level physicochemical descriptors. Positive values indicate higher values in the candidate cancer-associated set, and negative values indicate higher values in the healthy-background set. **d**, OmniTCR cancer-associated TCR enrichment scores (OmniTCR-CES) for control (n = 690), early-stage cancer (n = 60) and established cancer (n = 323) repertoires. Each point denotes one repertoire. Boxes show the median and interquartile range, and whiskers extend to 1.5 times the interquartile range. Group differences were assessed using Mann-Whitney U tests. **, $P < 0.01$; ****, $P < 0.0001$.

**Supplementary Fig. 5 | PMI reranking and TCRβ generation for unseen pMHCs.** **a-e,** Comparison of 3,222 experimentally observed reference TRB CDR3 sequences with 2,000 OmniTCR(PMI) candidates retained across 20 internal pMHC targets. **a,** CDR3β length distributions. **b,** OLGA $log_{10}$ generation-probability distributions. **c,** Global frequencies of the 20 standard amino acids in experimentally observed reference and generated sequences. The solid grey line denotes identity, and the dashed line is the fitted linear relationship (Pearson's $r$ = 0.982; $P = 1.6 \times 10^{-14}$). **d,** Sequence logos for experimentally observed reference and generated CDR3β sequences of lengths 13 and 15. **e,** Distribution of the difference between each candidate's mean per-residue log-likelihood under its target pMHC and its mean log-likelihood under the other 19 pMHC prompts. For each pMHC, margins were averaged across the 100 retained candidates and compared with zero using a one-sided Wilcoxon signed-rank test ($P = 9.5 \times 10^{-7}$). **f,** Target-versus-mean-noncognate likelihood margins for candidates retained under the original SFT ranking and PMI ranking at top-$k$ values of 10, 20, 50 and 100. **g,** Mean Jaccard similarity between each pMHC-conditioned candidate set and the other 19 target sets at the same cutoffs. In **f** and **g**, grey lines connect rankings for the same pMHC; comparisons used paired Wilcoxon signed-rank tests across 20 pMHCs. **h,** Percentage of PMI-ranked top-*k* candidates absent from the corresponding top-*k* set under the original SFT ranking. In **f-h**, small points denote individual target pMHCs, and summary points and error bars show the mean and 95% confidence interval estimated from 10,000 target-level bootstrap resamples. **i,** Char-BLEU, sequence recovery, F1@100 and pooled unique-sequence counts on 20 unseen pMHC targets comprising 1,036 experimentally observed reference TRB CDR3 sequences. All target peptide–HLA pairs were absent from pretraining and generation SFT. Each method generated 100 valid unique candidates per target, yielding 2,000 candidates per method. Bars show mean ± s.e.m. across the 20 pMHCs, except pooled unique-sequence counts. PMI, pointwise mutual-information-inspired; ****, $P < 0.0001$.

**Supplementary Fig. 6 | Best-of-100 AlphaFold 3 structural-confidence analysis of generated TCRβ candidates. a,** AlphaFold 3 (AF3) structural-confidence metrics for candidates selected from 100 generated CDR3β sequences per method for each of seven post-cutoff, epitope-excluded pMHCs. For each method and target, the candidate with the highest TCRβ ipTM was retained. The experimentally observed CDR3β sequence for each target was processed through the same modeling workflow. Points denote target-specific selected candidates, and bars show mean ± s.d. across the seven targets. Arrows indicate the favourable direction for each metric. **b,** AF3-predicted structure of the OmniTCR(PMI) CDR3β candidate CASSLGDSSYEQYF in the FSGEYIPTV-HLA-A*02:01 complex (PDB ID: 9K2I). The generated CDR3β loop is orange, the peptide is magenta and the remaining complex is shown in muted colours. The inset highlights selected heavy-atom contacts between the central LGDSSY segment and the peptide YIP core; dashed lines denote selected contacts within 4.0 Å. The predicted complex contained 19 CDR3β-peptide heavy-atom contacts within 4.0 Å, with the closest contact between TCRβ Tyr101 and peptide Pro7 at 3.06 Å. PMI, pointwise mutual-information-inspired; pTM, predicted template modelling score; ipTM, interface predicted template modelling score; iPAE, interface predicted alignment error; pLDDT, predicted local distance difference test.

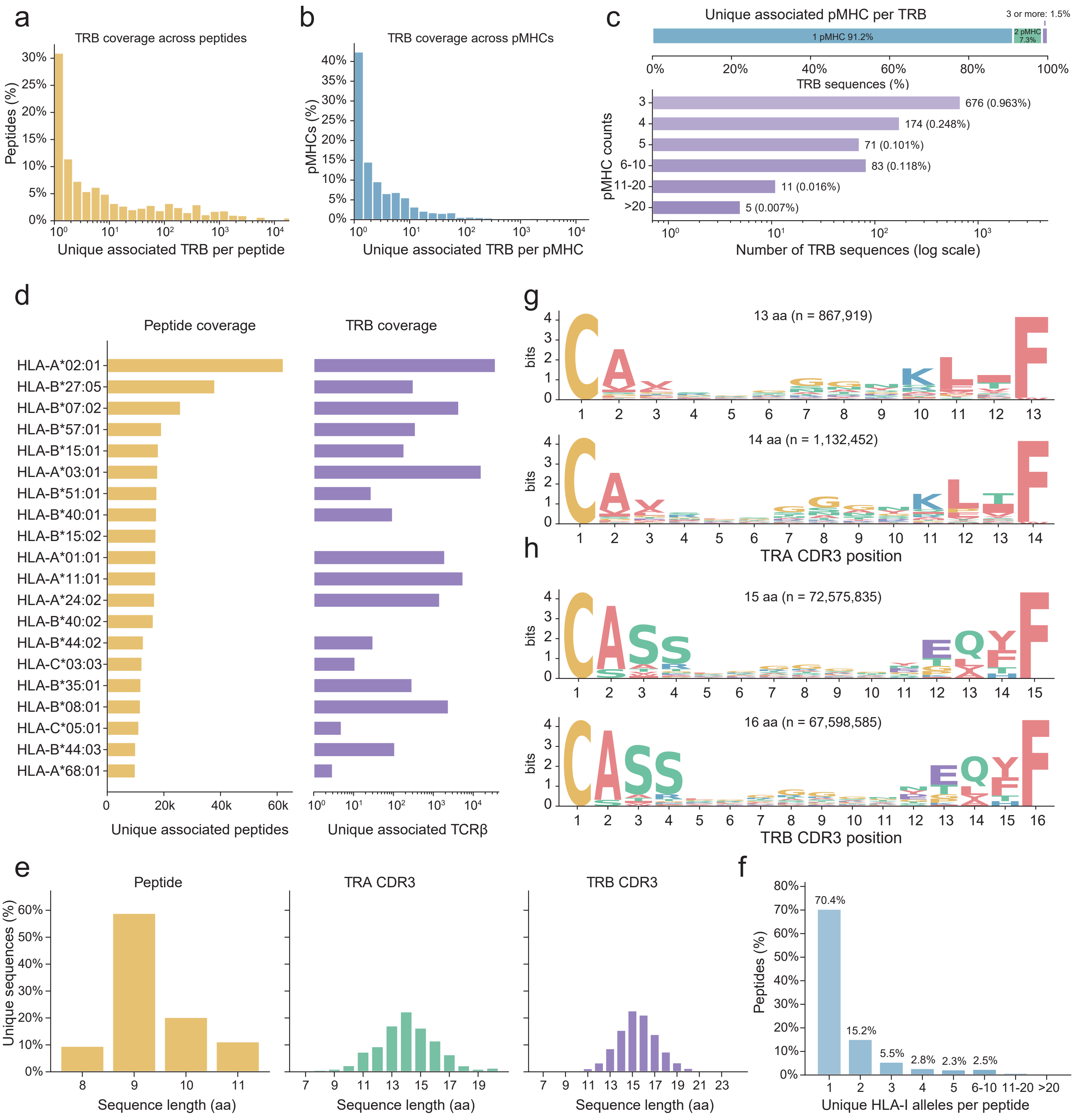
a
TRB coverage across peptides
Peptides (%)
Unique associated TRB per peptide
b
TRB coverage across pMHCs
pMHCs (%)
Unique associated TRB per pMHC
c
Unique associated pMHC per TRB
1 pMHC 91.2%
2 pMHC 7.3%
3 or more: 1.5%
TRB sequences (%)
pMHC counts
676 (0.963%)
174 (0.248%)
71 (0.101%)
83 (0.118%)
11 (0.016%)
5 (0.007%)
Number of TRB sequences (log scale)
d
Peptide coverage
TRB coverage
HLA-A*02:01
HLA-B*27:05
HLA-B*07:02
HLA-B*57:01
HLA-B*15:01
HLA-A*03:01
HLA-B*51:01
HLA-B*40:01
HLA-B*15:02
HLA-A*01:01
HLA-A*11:01
HLA-A*24:02
HLA-B*40:02
HLA-B*44:02
HLA-C*03:03
HLA-B*35:01
HLA-B*08:01
HLA-C*05:01
HLA-B*44:03
HLA-A*68:01
Unique associated peptides
Unique associated TCRβ
e
Peptide
TRA CDR3
TRB CDR3
Unique sequences (%)
Sequence length (aa)
f
Peptides (%)
70.4%
15.2%
5.5%
2.8%
2.3%
2.5%
Unique HLA-I alleles per peptide
g
13 aa (n = 867,919)
14 aa (n = 1,132,452)
TRA CDR3 position
h
15 aa (n = 72,575,835)
16 aa (n = 67,598,585)
TRB CDR3 position
bits

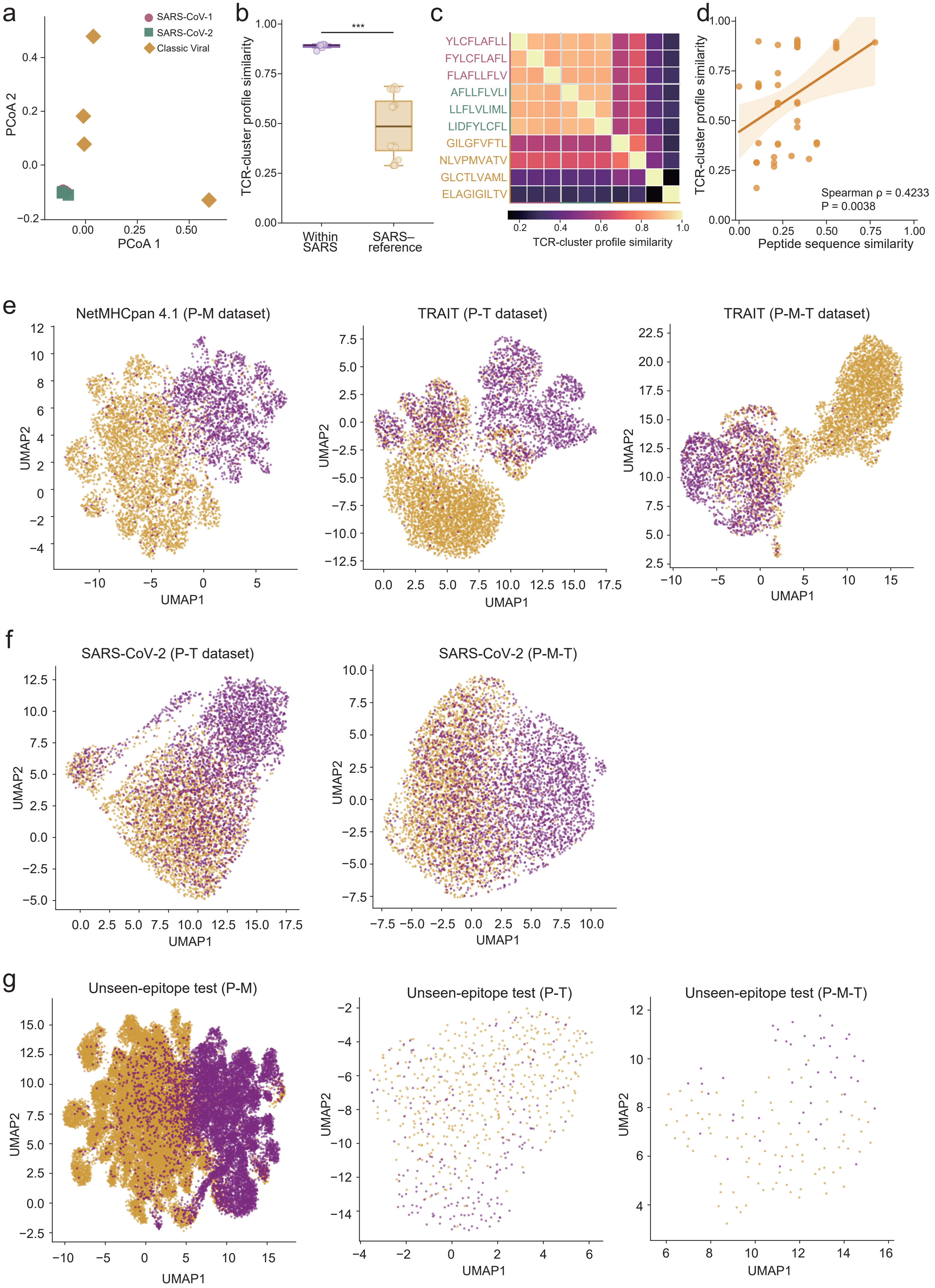

a
SARS-CoV-1
SARS-CoV-2
Classic Viral
PCoA 1
PCoA 2
b
***
TCR-cluster profile similarity
Within SARS
SARS–reference
c
YLCFLAFLL
FYLCFLAFL
FLAFLLFLV
AFLLFLVLI
LLFLVLIML
LIDFYLCFL
GILGFVFTL
NLVPMVATV
GLCTLVAML
ELAGIGILTV
TCR-cluster profile similarity
d
TCR-cluster profile similarity
Peptide sequence similarity
Spearman ρ = 0.4233
P = 0.0038
e
NetMHCpan 4.1 (P-M dataset)
TRAIT (P-T dataset)
TRAIT (P-M-T dataset)
UMAP1
UMAP2
f
SARS-CoV-2 (P-T dataset)
SARS-CoV-2 (P-M-T)
UMAP1
UMAP2
g
Unseen-epitope test (P-M)
Unseen-epitope test (P-T)
Unseen-epitope test (P-M-T)
UMAP1
UMAP2

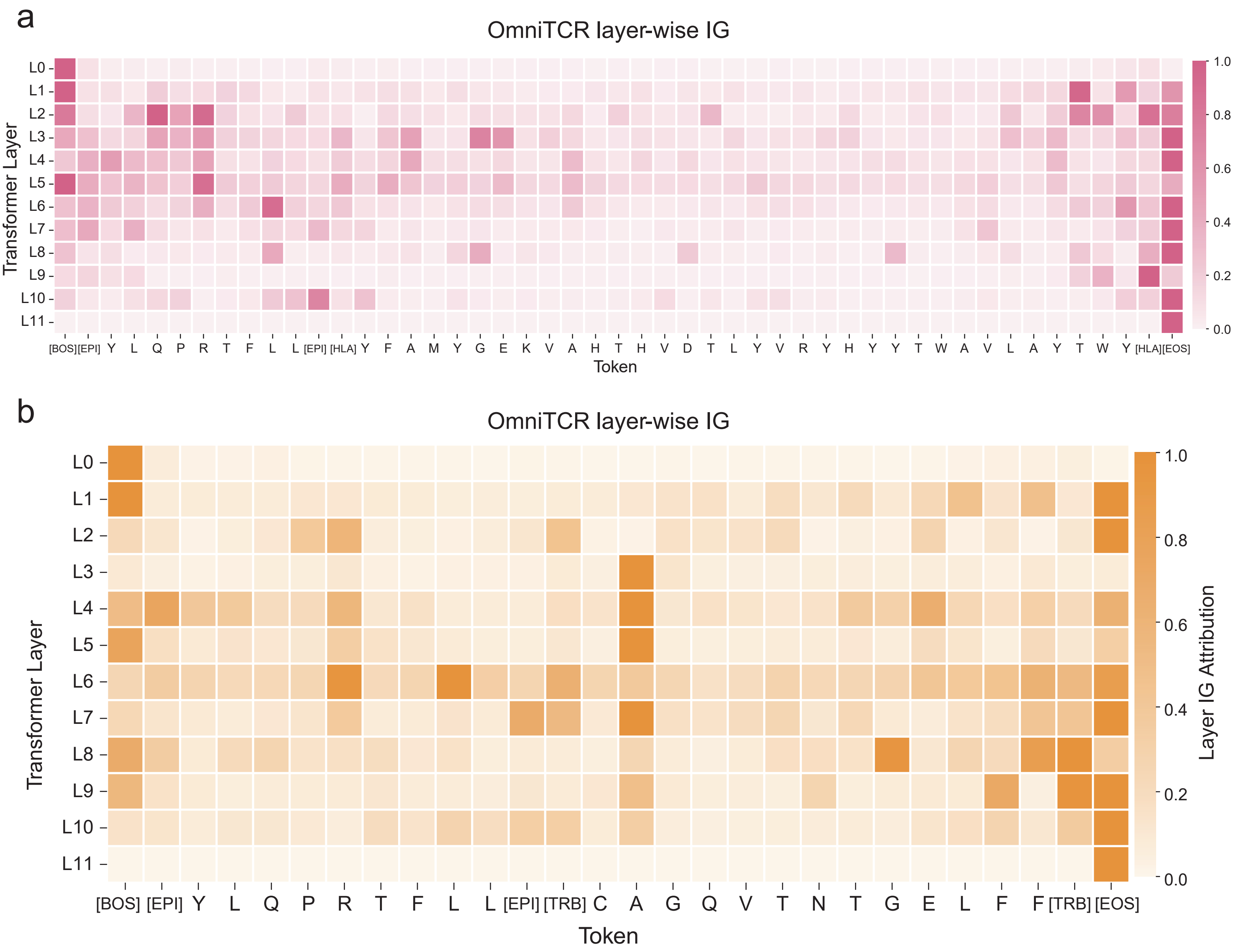
a
OmniTCR layer-wise IG
Transformer Layer
L0
L1
L2
L3
L4
L5
L6
L7
L8
L9
L10
L11
[BOS][EPI] Y L Q P R T F L L [EPI] [HLA] Y F A M Y G E K V A H T H V D T L Y V R Y H Y Y T W A V L A Y T W Y [HLA][EOS]
Token
1.0
0.8
0.6
0.4
0.2
0.0
b
OmniTCR layer-wise IG
Transformer Layer
L0
L1
L2
L3
L4
L5
L6
L7
L8
L9
L10
L11
[BOS] [EPI] Y L Q P R T F L L [EPI] [TRB] C A G Q V T N T G E L F F [TRB] [EOS]
Token
Layer IG Attribution
1.0
0.8
0.6
0.4
0.2
0.0

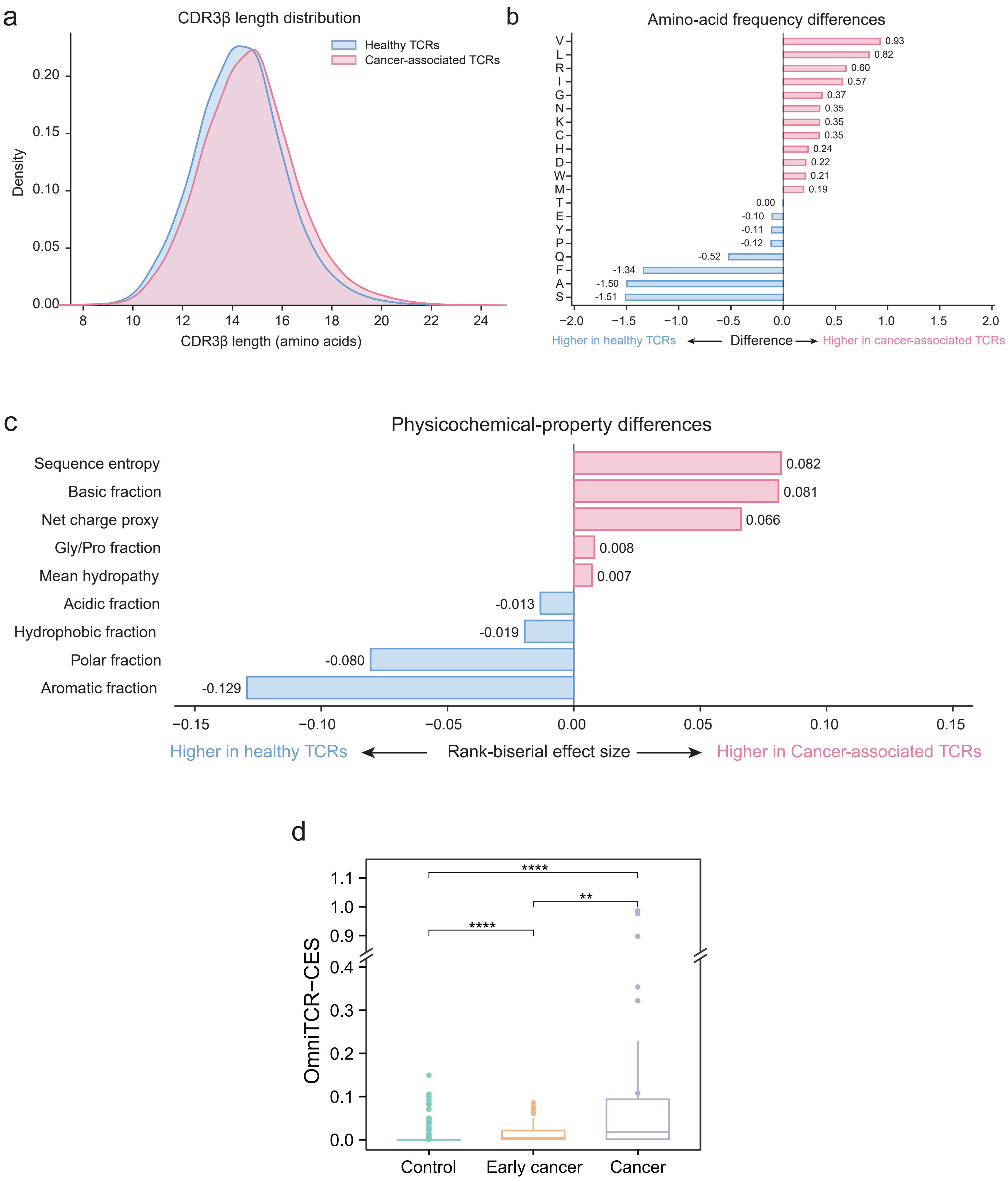
a
CDR3β length distribution
Healthy TCRs
Cancer-associated TCRs
Density
CDR3β length (amino acids)
b
Amino-acid frequency differences
V 0.93
L 0.82
R 0.60
I 0.57
G 0.37
N 0.35
K 0.35
C 0.35
H 0.24
D 0.22
W 0.21
M 0.19
T 0.00
E -0.10
Y -0.11
P -0.12
Q -0.52
F -1.34
A -1.50
S -1.51
Higher in healthy TCRs ← Difference → Higher in cancer-associated TCRs
c
Physicochemical-property differences
Sequence entropy 0.082
Basic fraction 0.081
Net charge proxy 0.066
Gly/Pro fraction 0.008
Mean hydropathy 0.007
Acidic fraction -0.013
Hydrophobic fraction -0.019
Polar fraction -0.080
Aromatic fraction -0.129
Higher in healthy TCRs ← Rank-biserial effect size → Higher in Cancer-associated TCRs
d
OmniTCR-CES
****
**
****
Control
Early cancer
Cancer

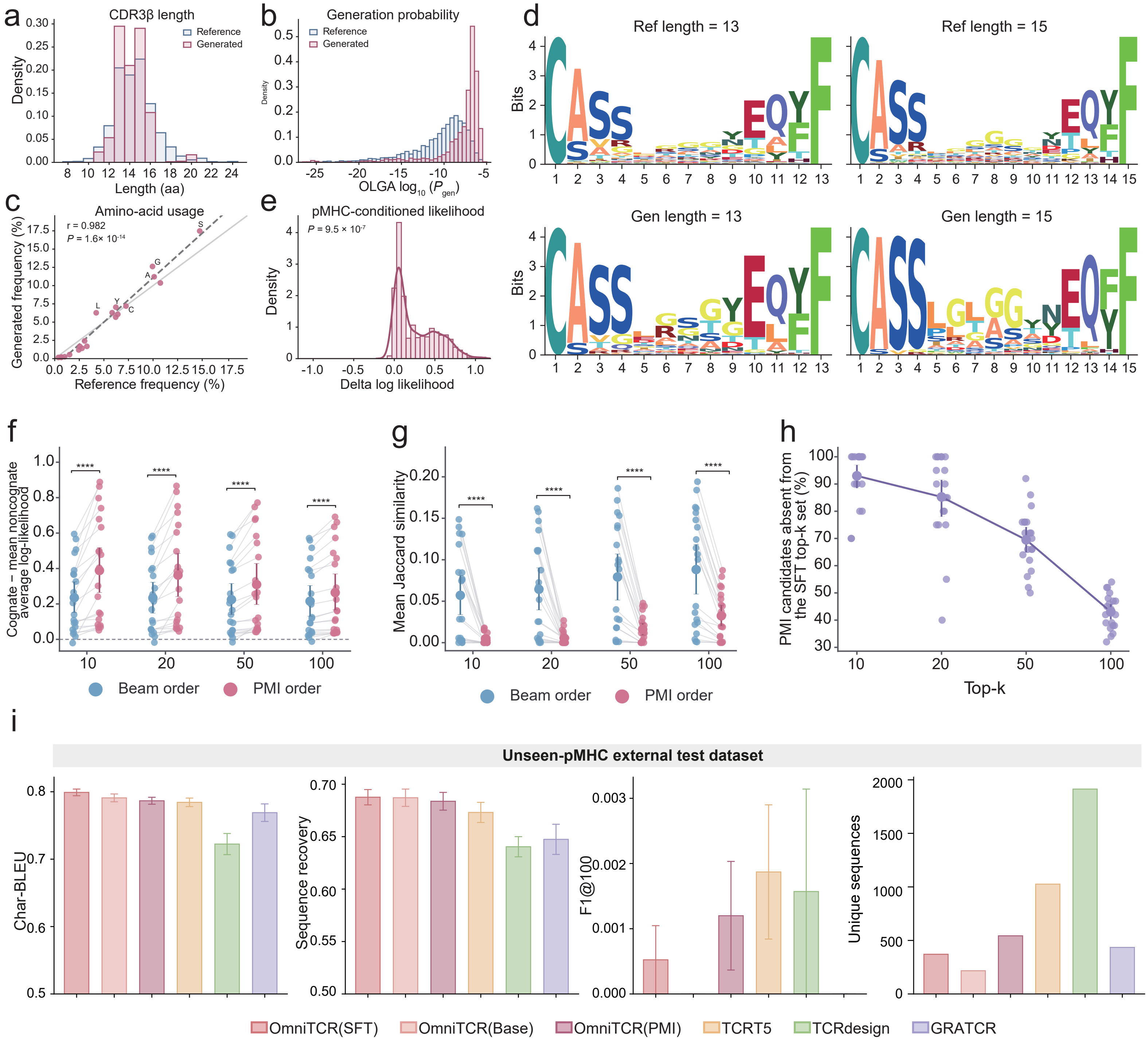

a
CDR3β length
Reference
Generated
Density
Length (aa)
b
Generation probability
Reference
Generated
Density
OLGA $\log_{10}(P_{gen})$
c
Amino-acid usage
r = 0.982
P = 1.6× 10-14
Generated frequency (%)
Reference frequency (%)
e
pMHC-conditioned likelihood
P = 9.5 × 10-7
Density
Delta log likelihood
d
Ref length = 13
Ref length = 15
Gen length = 13
Gen length = 15
Bits
f
Cognate − mean noncognate average log-likelihood
Beam order
PMI order
g
Mean Jaccard similarity
Beam order
PMI order
h
PMI candidates absent from the SFT top-k set (%)
Top-k
i
Unseen-pMHC external test dataset
Char-BLEU
Sequence recovery
F1@100
Unique sequences
OmniTCR(SFT)
OmniTCR(Base)
OmniTCR(PMI)
TCRT5
TCRdesign
GRATCR

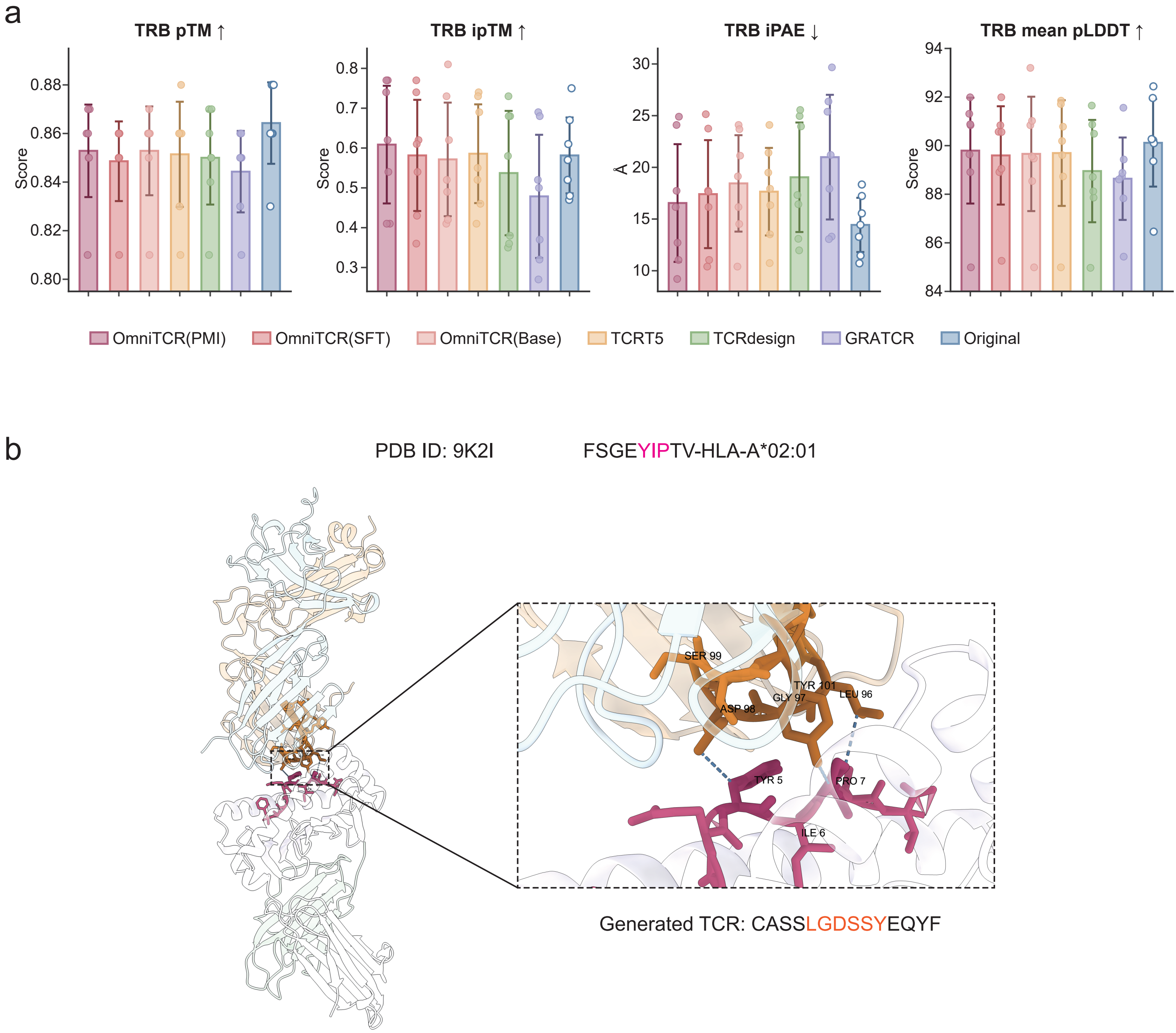
a
TRB pTM ↑
TRB ipTM ↑
TRB iPAE ↓
TRB mean pLDDT ↑
Score
Å
OmniTCR(PMI)
OmniTCR(SFT)
OmniTCR(Base)
TCRT5
TCRdesign
GRATCR
Original
b
PDB ID: 9K2I
FSGEYIPTV-HLA-A*02:01
SER 99
TYR 101
GLY 97
LEU 96
ASP 98
TYR 5
PRO 7
ILE 6
Generated TCR: CASSLGDSSYEQYF
